# In silico thermodynamic evaluation of the effectiveness of RT-LAMP primers to SARS-CoV-2 variants detection

**DOI:** 10.1101/2023.08.08.552530

**Authors:** Pâmella Miranda, Pedro A. Alves, Rubens L. do Monte-Neto, Gerald Weber

## Abstract

Viral mutations are the primary cause of mismatches in primer-target hybridisation, affecting the sensibility of molecular techniques, potentially leading to detection dropouts. Despite its importance, little is known about the quantitative effect of mismatches in primer-target hybridisation. We use up-to-date and highly detailed thermodynamic model parameters of DNA mismatches to evaluate the sensibility to variants of SARS-CoV-2 RT-LAMP primers. We aligned 18 RT-LAMP primer sets, which were underwent clinical validation, to the genomes of Wuhan strain (ws), 7 variants and 4 subvariants, and calculated hybridisation temperatures allowing up to three consecutive mismatches. We calculate the coverage when the mismatched melting temperature falls by more than 5^°^C in comparison to the matched alignments. If no mismatches are considered, the average coverage found would be 94% for ws, falling the lowest value for Omicron: 84%. However, considering mismatches the coverage is much higher: 97% (ws) to 88% (Omicron). Stabilizing mismatches (higher melting temperatures), account for roughly 1/3 of this increase. The number of primer dropouts increases for new each variant, however the effect is much less severe if mismatches are considered. We suggest using melting temperature calculations to continuously assess the trend of primer dropouts.

## 1. Introduction

There are eight possible mismatched (MM) base pairs in DNA: AA, AC, AG, CC, CT, GG, GT and TT. They may arise from DNA replication [1], genetic recombination [2] and primer-template hybridisation in PCR reactions [3]. Their presence may influence the stability and structural properties of DNA duplex changing hydrogen bonds conformation and stacking interactions. MM pairs may be found in anti-syn or syn-anti conformations differently from DNA pairs which are naturally in an anti-anti conformation. Mismatch impact varies from weakly bound (CC pair) to strongly bound (GG pair) in a local conformation, while molecular dynamics and NMR experiments have shown no impact in a global conformation such as for AA and TT pairs [1, 4]. Although it is known that mismatches typically destabilise the primer-target duplex, some types of mismatches are more stable than others, some even more than AT base pairs, which may contribute towards the stability of the duplex [1, 5, 6]. A few mismatches in PCR primers may contribute to design of antisense oligonucleotides [7], SNP [8] and allele-specific identification [6].

The isothermal PCR known as RT-LAMP (reverse transcription loop-mediated isothermal amplification) is a robust, fast, inexpensive molecular technique and which can be carried out in less than an hour [9, 10]. It has been used as molecular diagnostic test for several diseases such as ebola [11], zika [12], HIV [13], SARS [14], MERS-CoV [15] and SARS-CoV-2 [16], the causative agent of COVID-19. To detect those diseases, it is necessary to design specific primers to identify the target agent. Unlike PCR primers, which usually uses a single pair of primers, RT-LAMP uses 2 or 3 pairs: F3 and B3 (outer primers), FIP and BIP (inner primers) and LF and LB (loop primers). The outer and inner primers act in the beginning of the reaction, but just the inner ones act in later cycles. FIP and BIP primers are long primers which contain two parts: F1c and F2 for FIP and B1c and B2 for BIP, which correspond to sense and antisense sequences of the target [17]. Finally, the loop primers are included to accelerate the reaction [18].

For LAMP, in the same way as for PCR [19], mismatches may appear between target and primers due to mutations potentially causing false-negative results [20, 21]. In fact, SARS-CoV-2 PCR primers designed by Corman et al. [21] during the earlier stages of pandemic have mismatches which have not hindered the detection of the coronavirus [22].

In a previous study [23], we evaluated the impact of mismatches on RT-PCR primers and probes, where we showed that the mismatches do not always have a negative impact on the thermodynamic stability. The reason for this is that there is a number of mismatch configurations that may actually increase the melting temperatures. This was confirmed recently by Scapaticci et al. [24] who found that mutations may have higher melting temperatures and suggested that the melting temperature analysis could be used to detect specific variants. As for PCR, it is expected that mismatches in primer-target hybridisation will appear for LAMP primers. Specially for both FIP and BIP primers in which mismatched base pairs in either 5^′^ or 3′ ends may prevent the elongation by Bst DNA polymerase leading to a low amplification efficiency [25]. Although one or two mismatches were shown to be tolerable for LAMP [26, 27], studies with three or more consecutive mismatches are, to our knowledge, not available.

Here we show the evaluation of DNA mismatches in 18 RT-LAMP primer sets [28–45], which were designed for SARS-CoV-2 original genomes. One of those sets [29] was previously successfully evaluated by us for a few variants and now for an amplified genome sets. We applied a previous workflow [23] to analyse those primers for the detection of SARS-CoV-2 variants as Alpha (B.1.1.7), Beta (B.1.351), Gamma (P.1), Delta (B.1.617.2), Lambda (C.37), Mu (B.1.621), Omicron (B.1.1.529) variants and BA.2 to BA.5 subvariants. The outcomes show if those primers may still be effective to detect the variants and how the presence of mismatches may contribute to cover more genomes, consequently, detect the coronavirus. Furthermore, we reinforce the fact that a continuous evaluation of RT-LAMP primer sets is needed to cover variants that may arise as already suggested [39].

## 2. Materials and Methods

### 2.1. Genome sets

We collected randomly 21665 genomes of original SARS-CoV-2 (Wuhan strain) in 8 October 2020, at NCBI [46]; 7247 genomes of Alpha variant, 7497 of Beta variant and 2308 of Gamma variant in 7 April 2021; 7943 of Delta variant in 5 June 2021; 7029 of Omicron variant in 16 December 2021; 6610 of Mu variant, 9340 of Lambda variant, 7393 and 348 of Omicron subvariants BA.2 and BA.3 in 11 February 2022; 629 and 1231 of Omicron subvariants BA.4 and BA.5 in 19 September 2022, at GISAID [47] (data not shown).

### 2.2. Primer sets

We collected 18 different RT-LAMP primer sets designed to SARS-CoV-2 original genomes that underwent clinical validation from Refs. [28–45] in a total of 436 primers. Their details are shown in Supplementary Table S1. FIP and BIP primers were divided in F1c/F2 and B1c/B2 primers, respectively, except those from three sets [29, 31, 42], for which the division of primers was already given. We found all possible combinations of primer pairs and selected those according to the temperatures of the same type pair from the three sets just mentioned.

### 2.3. Evaluation workflow

All primers were aligned to each genome set using a Smith-Waterman algorithm, as described in [23]. Fully matched alignments were called strictly matched and those with single, double and triple consecutive mismatches were termed partially matched. Alignments with four or more consecutive mismatches were considered as not aligned. The limit of the maximal number of consecutive mismatches is due to fact that the available parameter only covers up to three contiguous mismatches [5]. In addition, it is very likely that four or more mismatches will destabilize the primers far beyond the limits considered here. Also deletions in the viral genome, as in Omicron variant [48, 49], may lead to no alignment of the primers.

Hybridisation temperatures for matched (*T*_ref._) and mismatched (*T*_MM_) alignments are calculated from a mesoscopic model with the parameters from [5].

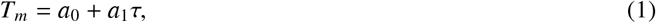

The reference hybridisation temperature *T*_ref._ for each primer is shown in Supplementary Table S1. Note that the parameters from Ref. [5] are for a sodium buffer which is different from those typically used in PCR reactions which contain Mg^+^. Therefore, the absolute temperatures *T*_ref._ will be different from the actual melting temperatures of the primers. However, since our analysis deals with temperature differences, which are not strongly buffer dependent, we expect them to be sufficiently accurate for our purposes.

We define a strictly matched (AT and CG only) alignment coverage for each primer as

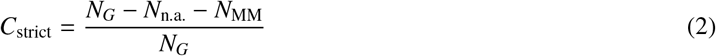

where *N*_*G*_ is the total number of genomes, *N*_n.a._ the number of genomes for which no alignment was found and *N*_MM_ the number of genomes for which a partial alignment containing mismatches was found.

The difference between reference hybridisation temperature *T*_ref._ and mismatched alignments *T*_MM_

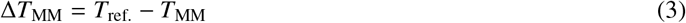

where *T*_MM_ is usually lower than *T*_ref._ [5]. The partial coverage for alignments with up to three contiguous mismatches

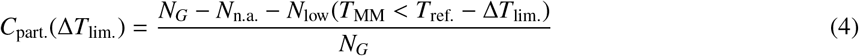

where *N*_low_ is the number of alignments where the mismatched melting temperature *T*_MM_ is lower by Δ*T*_lim._ than the reference *T*_ref._. Note that as there are many mismatch configurations that have an increased *T*_MM_, that is, there are situations where *C*_part._ > *C*_strict_ even for Δ*T*_lim._ = 0. For additional details of this workflow see Ref. [23].

All 18 primer sets were aligned against the genomes of SARS-CoV-2 variants. We calculated the hybridisation temperatures and coverages for both matched and mismatched alignments considering single, double and triple consecutive mismatches. The complete evaluation was carried out in approximately 120 h computing time.

### 2.4. Availability

The software packages used to carry out this work are freely available and can be found in https://bioinf.fisica.ufmg.br/software/, in the *analyse primer lamp.tar.gz* package. The authors will consider requests for the analysis of primer sets, please see contact details.

## 3. Results and Discussion

Given the continuous mutation of the SARS-CoV-2 genomes, it is expected that over time mismatches should increasingly occur within the primer regions. Figure 1, where we show the *C*_strict_ averaged over all 436 primers, illustrates this decreasing coverage as variants appear. In comparison to the ws coverage, all variants decrease their coverage. When we consider partial coverages in the presence of mismatches with Δ*T*_lim_ = 0^°^C, that is, primers with *T*_MM_ ≥ *T*_ref._, the curve is uniformly shifted upwards. For Δ*T*_lim_ = 5^°^C, Beta, Gamma, Delta and Mu partial coverage becomes slightly higher than the ws strict coverage. However, the rate of decrease is not uniform, and some variants have higher coverage than their presumed predecessor variants. For the Omicron variant, which has a larger number of mutations [50], we observe a sharp drop in the coverage. However for the subsequent subvariant the picture is mixed, the BA.3 subvariant shares the low coverage, but BA.2, BA.4 and BA.5 have a higher coverage. The reason for this oscillation is not clear.

**Figure 1:**
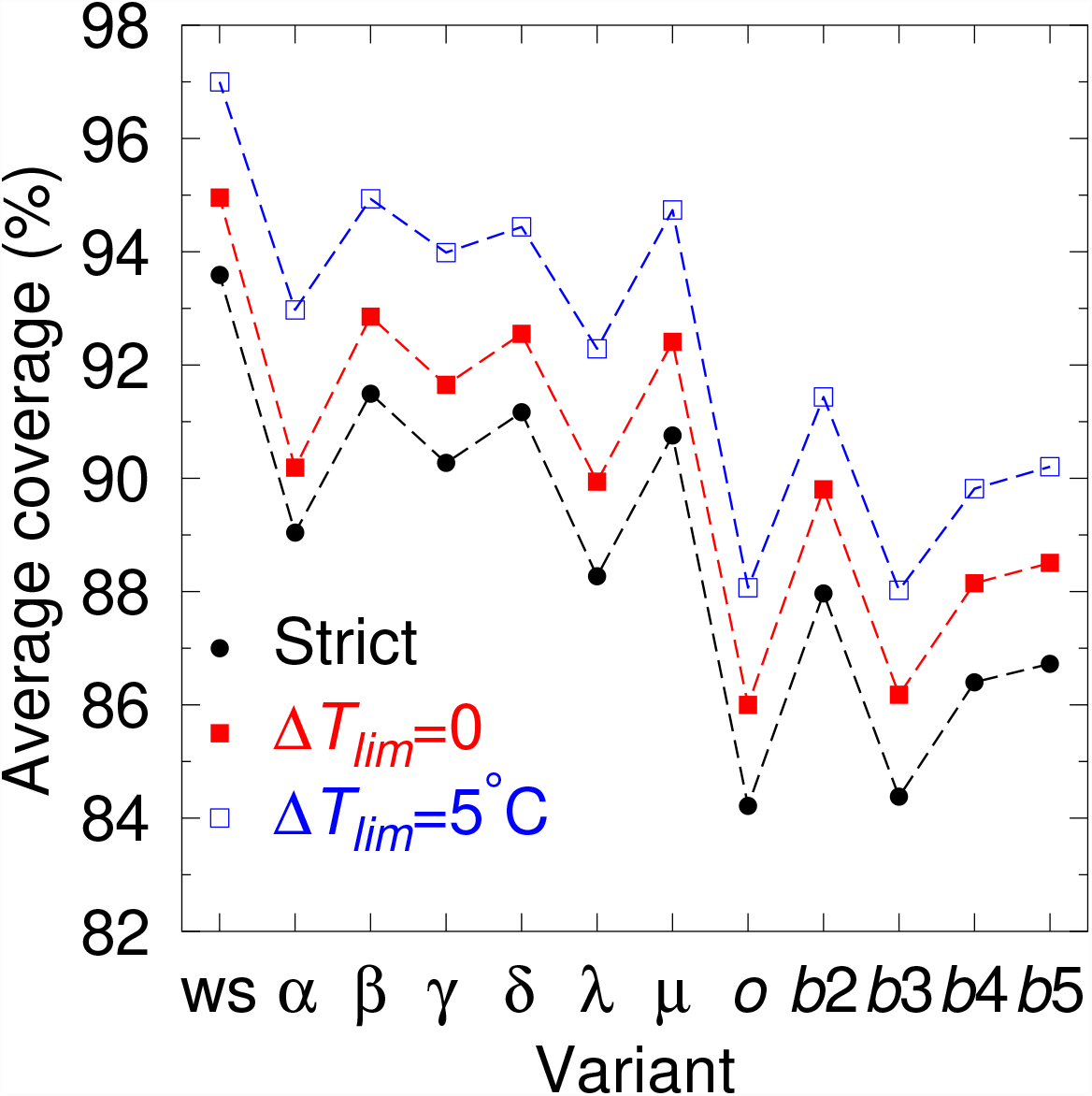
Coverage averaged over all primers as for SARS-CoV-2 original genomes (ws), Alpha (*α*), Beta (*β*), Gamma (*γ*), Delta (*δ*), Lambda (*#x03BB;*), Mu (*μ*), Omicron (*o*) variants and BA.2 (b2), BA.3 (b3), BA.4 (b4) and BA.5 (b5) subvariants. Black bullet are for *C*_strict_ and red (blue) boxes are for Δ*T*_lim_ = 0°C (5°C). The dashed line connecting the data point is only intended as a guide to the eye.

Similar to what we have seen for RT-PCR [51], many alignments that would result in a null strict coverage achieve partial coverage beyond 99% if mismatches are considered. In some cases, a large partial coverage is already obtained for Δ*T*_lim_ = 0°C, that is, if we consider only mismatches that do not destabilize the duplex. In Table 1 we show a few examples of primers that have zero strict coverage but go beyond 90% if stabilizing mismatches are considered. It is somewhat surprising that some primers achieve high coverages for Omicron only and not for the other variants, despite the fact that all were designed for the Wuhan strain. While this seems to be an opposite trend to the overall decline for Omicron, one should note that a higher coverage for Omicron is rather exceptional and only occurs for very few primers. On the other hand, it highlights quite clearly that the assessment of mismatch influence is far from trivial.

**Table 1:**
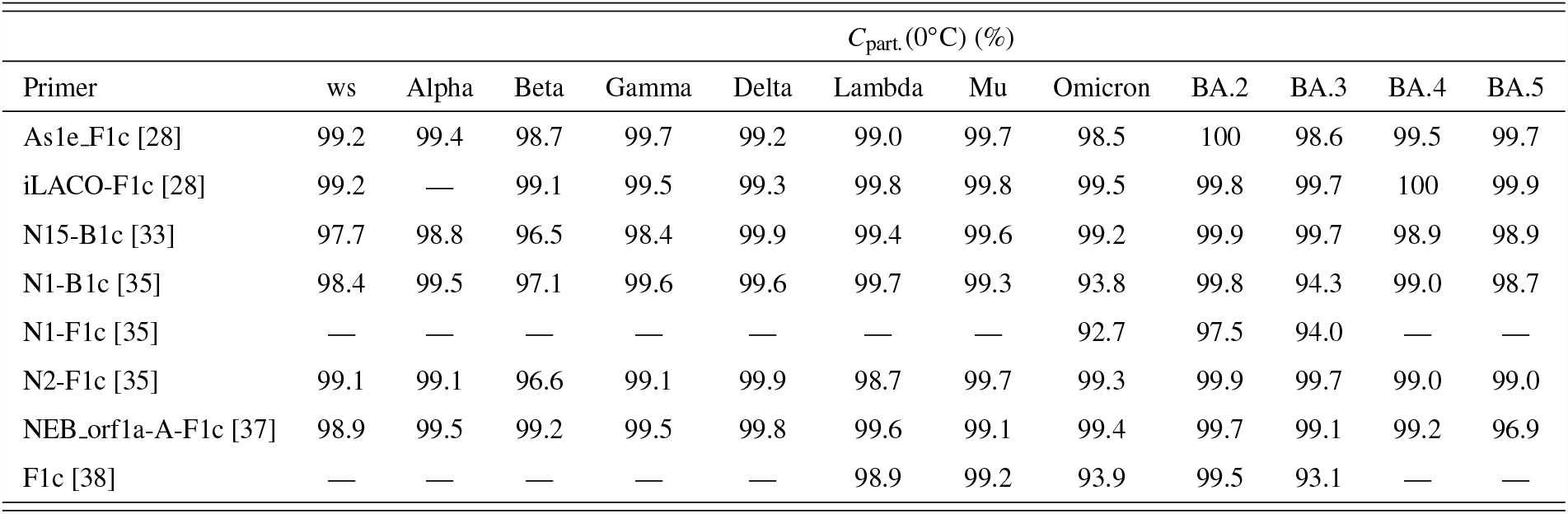
Examples of primer coverages with stabilizing mismatches, *T*_lim._ = 0°C, who have *C*_part._(0°C) > 90% while having *C*_strict_ = 0. Only those primers which do have stabilizing mismatches for given variant shown.

While most primers have large coverages, an important amount of primers fail to achieve significant coverage for at least one variant and may represent a potential dropout. A summary of the amount of primers found that could potential represent dropouts are shown in Table 2. Here we are considering a very stringent threshold of 5% at Δ*T*_lim_ = 5°C, that is, primers where even considering a maximal 5°C melting temperature below the reference temperature covered less than 5% of the available genomes for a given variant. Only Alekseenko et al. [28] set had no potential dropout primers at all. The complete list of potential dropout primers for each variant is shown in supplementary Tables S56–S67.

**Table 2:** Sets that have at least one potential drop-out primer for any of the variants. Only the reference number is shown for each set. Drop-out primers are considered as those with a partial coverage (Δ*T*_lim._ = 5°C) below 5%, N_drop_, for at least one variant. *N*_primers_ is the number of separate primers for each set.

| Ref. | $N_{\text{primers}}$ | $N_{\text{drop}}$ | ws | Alpha | Beta | Gamma | Delta | Lambda | Mu | Omicron | BA.2 | BA.3 | BA.4 | BA.5 |
| --- | --- | --- | --- | --- | --- | --- | --- | --- | --- | --- | --- | --- | --- | --- |
| [29] | 32 | 6 | 0 | 0 | 0 | 1 | 0 | 1 | 1 | 3 | 3 | 3 | 4 | 3 |
| [30] | 40 | 7 | 0 | 1 | 1 | 1 | 1 | 0 | 0 | 3 | 3 | 3 | 3 | 3 |
| [31] | 81 | 28 | 4 | 11 | 5 | 5 | 5 | 6 | 12 | 9 | 5 | 10 | 7 | 6 |
| [32] | 8 | 1 | 0 | 0 | 0 | 0 | 0 | 1 | 0 | 0 | 0 | 0 | 0 | 0 |
| [33] | 32 | 6 | 0 | 0 | 0 | 1 | 0 | 3 | 1 | 3 | 2 | 3 | 2 | 2 |
| [34] | 15 | 3 | 0 | 0 | 1 | 0 | 0 | 0 | 0 | 1 | 2 | 2 | 2 | 2 |
| [35] | 32 | 5 | 1 | 3 | 1 | 3 | 2 | 2 | 1 | 2 | 2 | 2 | 3 | 2 |
| [36] | 8 | 1 | 0 | 0 | 0 | 0 | 0 | 0 | 0 | 1 | 1 | 1 | 1 | 1 |
| [37] | 32 | 5 | 0 | 0 | 0 | 0 | 0 | 1 | 0 | 1 | 2 | 2 | 4 | 2 |
| [38] | 10 | 6 | 0 | 2 | 1 | 0 | 2 | 3 | 0 | 3 | 3 | 3 | 3 | 3 |
| [39] | 48 | 12 | 0 | 3 | 0 | 5 | 3 | 6 | 0 | 7 | 7 | 7 | 8 | 7 |
| [40] | 8 | 5 | 0 | 2 | 0 | 1 | 0 | 0 | 2 | 0 | 0 | 0 | 0 | 0 |
| [41] | 16 | 5 | 0 | 1 | 2 | 0 | 1 | 1 | 0 | 1 | 1 | 1 | 1 | 1 |
| [42] | 8 | 2 | 1 | 1 | 1 | 2 | 1 | 1 | 1 | 1 | 1 | 1 | 1 | 1 |
| [43] | 15 | 3 | 0 | 1 | 0 | 0 | 0 | 1 | 0 | 1 | 1 | 1 | 1 | 1 |
| [44] | 15 | 2 | 0 | 1 | 0 | 1 | 1 | 2 | 0 | 1 | 1 | 1 | 1 | 1 |
| [45] | 16 | 1 | 1 | 1 | 1 | 1 | 1 | 1 | 1 | 1 | 1 | 1 | 1 | 1 |

Mismatched pairs in 5′ or 3′ terminals of FIP and BIP primers may hamper the amplification by Bst DNA polymerase. However we found a few alignments with either 5′ and 3′ terminal mismatches which have a hybridisation temperature within the threshold and contribute to the increase in the coverage when mismatches are taken into account. Clearly, in some cases, mismatches in both terminals reduced the temperature. FIP and BIP primers with terminal mismatches which have an increase in their coverage are shown in supplementary Tables S68–S79 for each genome set. Note that FIP and BIP primers were divided into F1c/F2 and B1c/B2, respectively, and as such treated individually. Since for the LAMP technique the F1c and B1c depend on their respective F2 and B2 complements, the dropout may in practice be higher.

## 4. Conclusion

We evaluated the coverage of 18 RT-LAMP primer sets considering single, double and triple mismatches in primer-target hybridisation to SARS-CoV-2 variants. In general, the average coverage of these primer sets decreased for the new variants, when compared to the Wuhan strain. Overall, the coverage was lowest for the Omicron and BA.3 variants. However, a clear monotonic decrease of the coverage is not observed, instead for some variants the coverage increases when compared to its putative predecessor, as exemplified most notably by the Mu variant which showed one of the highest coverages. Coverage uniformly increases if mismatches are taken into account, while not enough to completely compensate the loss in comparison to the Wuhan strain, it shifts the worst case from 84% to 88%. Similarly, the number potential dropout primers increased with each new variant, and only one out of 18 sets did show no potential primer drop-out. We suggest the use of the methodology described here to continuously evaluate the effectiveness of RT-LAMP primer as new variants emerge. Furthermore, our method could be applied to the detection of other infectious diseases.

## Supplementary data

Table S1 show all primers used and their reference temperatures. Tables S2–S55 show the strict and partial coverages of all primers. Tables S56–S67 show the potential drop-out primers for each variant. Tables S68–S79 show the terminal mismatched positions of FIP and BIP primers for each genome.

