## supplementary material for "In silico thermodynamic evaluation of the effectiveness of RT-LAMP primers to SARS-CoV-2 variants detection"

---

---

*Corresponding author

*August 8, 2023*

S27 Strict and Partial Coverages for Jiang et al. set. . . . . . . . . . . . . . . . . . . . . . . . . . . 49
S28 Strict and Partial Coverages for Jiang et al. set. . . . . . . . . . . . . . . . . . . . . . . . . . . 50
S29 Strict and Partial Coverages for Lalli et al. set. . . . . . . . . . . . . . . . . . . . . . . . . . . 51
S30 Strict and Partial Coverages for Lalli et al. set. . . . . . . . . . . . . . . . . . . . . . . . . . . 52
S31 Strict and Partial Coverages for Lalli et al. set. . . . . . . . . . . . . . . . . . . . . . . . . . . 53
S32 Strict and Partial Coverages for Lau et al. set. . . . . . . . . . . . . . . . . . . . . . . . . . . 54
S33 Strict and Partial Coverages for Lau et al. set. . . . . . . . . . . . . . . . . . . . . . . . . . . 55
S34 Strict and Partial Coverages for Lau et al. set. . . . . . . . . . . . . . . . . . . . . . . . . . . 56
S35 Strict and Partial Coverages for Luo et al. set. . . . . . . . . . . . . . . . . . . . . . . . . . . 57
S36 Strict and Partial Coverages for Luo et al. set. . . . . . . . . . . . . . . . . . . . . . . . . . . 59
S37 Strict and Partial Coverages for Luo et al. set. . . . . . . . . . . . . . . . . . . . . . . . . . . 61
S38 Strict and Partial Coverages for Mautner et al. set. . . . . . . . . . . . . . . . . . . . . . . . . 63
S39 Strict and Partial Coverages for Mautner et al. set. . . . . . . . . . . . . . . . . . . . . . . . . 64
S40 Strict and Partial Coverages for Mautner et al. set. . . . . . . . . . . . . . . . . . . . . . . . . 65
S41 Strict and Partial Coverages for Mohon et al. set. . . . . . . . . . . . . . . . . . . . . . . . . 66
S42 Strict and Partial Coverages for Mohon et al. set. . . . . . . . . . . . . . . . . . . . . . . . . 67
S43 Strict and Partial Coverages for Mohon et al. set. . . . . . . . . . . . . . . . . . . . . . . . . 68
S44 Strict and Partial Coverages for Rodriguez-Manzano et al. set. . . . . . . . . . . . . . . . . . 69
S45 Strict and Partial Coverages for Rodriguez-Manzano et al. set. . . . . . . . . . . . . . . . . . 70
S46 Strict and Partial Coverages for Rodriguez-Manzano et al. set. . . . . . . . . . . . . . . . . . 71
S47 Strict and Partial Coverages for Yan et al. set. . . . . . . . . . . . . . . . . . . . . . . . . . . 72
S48 Strict and Partial Coverages for Yan et al. set. . . . . . . . . . . . . . . . . . . . . . . . . . . 73
S49 Strict and Partial Coverages for Yan et al. set. . . . . . . . . . . . . . . . . . . . . . . . . . . 74
S50 Strict and Partial Coverages for Yang et al. set. . . . . . . . . . . . . . . . . . . . . . . . . . . 75
S51 Strict and Partial Coverages for Yang et al. set. . . . . . . . . . . . . . . . . . . . . . . . . . . 76
S52 Strict and Partial Coverages for Yang et al. set. . . . . . . . . . . . . . . . . . . . . . . . . . . 77
S53 Strict and Partial Coverages for Yoshikawa et al. set. . . . . . . . . . . . . . . . . . . . . . . 78
S54 Strict and Partial Coverages for Yoshikawa et al. set. . . . . . . . . . . . . . . . . . . . . . . 79
S55 Strict and Partial Coverages for Yoshikawa et al. set. . . . . . . . . . . . . . . . . . . . . . . 80
S56 Potential drop-out primers for SARS-CoV-2. . . . . . . . . . . . . . . . . . . . . . . . . . . . 81
S57 Potential drop-out primers for Alpha variant. . . . . . . . . . . . . . . . . . . . . . . . . . . . 82
S58 Potential drop-out primers for Beta variant. . . . . . . . . . . . . . . . . . . . . . . . . . . . 83
S59 Potential drop-out primers for Gamma variant. . . . . . . . . . . . . . . . . . . . . . . . . . . 84
S60 Potential drop-out primers for Delta variant. . . . . . . . . . . . . . . . . . . . . . . . . . . . 85
S61 Potential drop-out primers for Lambda variant. . . . . . . . . . . . . . . . . . . . . . . . . . . 86
S62 Potential drop-out primers for Mu variant. . . . . . . . . . . . . . . . . . . . . . . . . . . . . 87
S63 Potential drop-out primers for Omicron variant. . . . . . . . . . . . . . . . . . . . . . . . . . 88
S64 Potential drop-out primers for BA.2 subvariant. . . . . . . . . . . . . . . . . . . . . . . . . . 90
S65 Potential drop-out primers for BA.3 subvariant. . . . . . . . . . . . . . . . . . . . . . . . . . 91
S66 Potential drop-out primers for BA.4 subvariant. . . . . . . . . . . . . . . . . . . . . . . . . . 93
S67 Potential drop-out primers for BA.5 subvariant. . . . . . . . . . . . . . . . . . . . . . . . . . 95
S68 Mismatched position in FIP and BIP primers for SARS-CoV-2. . . . . . . . . . . . . . . . . 96
S69 Mismatched position in FIP and BIP primers for Alpha variant. . . . . . . . . . . . . . . . . 97
S70 Mismatched position in FIP and BIP primers for Beta variant. . . . . . . . . . . . . . . . . . 98
S71 Mismatched position in FIP and BIP primers for Gamma variant. . . . . . . . . . . . . . . . 99

S72 Mismatched position in FIP and BIP primers for Delta variant. . . . . . . . . . . . . . . . . 100
S73 Mismatched position in FIP and BIP primers for Lambda variant. . . . . . . . . . . . . . . . 101
S74 Mismatched position in FIP and BIP primers for Mu variant. . . . . . . . . . . . . . . . . . 102
S75 Mismatched position in FIP and BIP primers for Omicron variant. . . . . . . . . . . . . . . 103
S76 Mismatched position in FIP and BIP primers for BA.2 subvariant. . . . . . . . . . . . . . . 104
S77 Mismatched position in FIP and BIP primers for BA.3 subvariant. . . . . . . . . . . . . . . 105
S78 Mismatched position in FIP and BIP primers for BA.4 subvariant. . . . . . . . . . . . . . . 106
S79 Mismatched position in FIP and BIP primers for BA.5 subvariant. . . . . . . . . . . . . . . 107

Table S1: Summary of primers for each set and their calculated hybridisation temperature $T_{ref.}$. We only considered new sequences from each publication, that is, those which to our knowledge have not been reported previously.

| Name of Set | Primer | Sequence $5' \rightarrow 3'$ | $T_{ref.}$ (°C) |
|---|---|---|---|
| Alekseenko et al. [1] | iLACO-F3 | CCACTAGAGGAGCTACTGTA | 59.5 |
| Alekseenko et al. [1] | iLACO-B3 | TGACAAGCTACAACACGT | 54.8 |
| Alekseenko et al. [1] | iLACO-F1c | AGGTGAGGGTTTTCTACATCACTATAT | 69.1 |
| Alekseenko et al. [1] | iLACO-F2 | TGGAACAAGCAAATTCTATGG | 60.3 |
| Alekseenko et al. [1] | iLACO-B1c | ATGGGTTGGGATTATCCTAAATGTG | 68.0 |
| Alekseenko et al. [1] | iLACO-B2 | TGCGAGCAAGAACAAGTG | 59.2 |
| Alekseenko et al. [1] | iLACO-LF | CAGTTTTTAACATGTTGTGCCAACC | 67.4 |
| Alekseenko et al. [1] | iLACO-LB | TAGAGCCATGCCTAACATGCT | 65.1 |
| Alekseenko et al. [1] | As1_F3 | CGGTGGACAAATTGTCAC | 56.7 |
| Alekseenko et al. [1] | As1_B3 | CTTCTCTGGATTTAACACACTT | 59.3 |
| Alekseenko et al. [1] | As1_LF | TTACAAGCTTAAAGAATGTCTGAACACT | 70.0 |
| Alekseenko et al. [1] | As1_LB | TTGAATTTAGGTGAAACATTTGTCACG | 69.4 |
| Alekseenko et al. [1] | As1_F1c | TCAGCACACAAAGCCAAAAATTTATC | 69.6 |
| Alekseenko et al. [1] | As1_F2 | TGTGCAAAGGAAATTAAGGAG | 60.2 |
| Alekseenko et al. [1] | As1_B1c | TATTGGTGGAGCTAAACTTAAAGCC | 68.6 |
| Alekseenko et al. [1] | As1_B2 | CTGTACAATCCCTTTGAGTG | 58.0 |
| Alekseenko et al. [1] | As1e_F1c | TCAGCACACAAAGCCAAAAATTTATTTTT | 72.3 |
| Alekseenko et al. [1] | As1e_F2 | CTGTGCAAAGGAAATTAAGGAG | 62.6 |
| Alekseenko et al. [1] | As1e_B1c | TATTGGTGGAGCTAAACTTAAAGCCT | 70.2 |
| Alekseenko et al. [1] | As1e_B2 | TTTCTGTACAATCCCTTTGAGTG | 63.4 |
| Alves et al. [2] | N_Set1_F3 | TGGCTACTACCGAAGAGCT | 61.6 |
| Alves et al. [2] | N_Set1_B3 | TGCAGCATTGTTAGCAGGAT | 62.5 |
| Alves et al. [2] | N_Set1_F1c | TCTGGCCCAGTTCCTAGGTAGT | 68.1 |
| Alves et al. [2] | N_Set1_F2 | GACGAATTCGTGGTGGTGA | 61.9 |
| Alves et al. [2] | N_Set1_B1c | AGACGGCATCATATGGGTTGCA | 69.2 |
| Alves et al. [2] | N_Set1_B2 | CGGGTGCCAATGTGATCT | 60.6 |
| Alves et al. [2] | N_Set1_LF | TGGACTGAGATCTTTCATTTTACCG | 69.3 |
| Alves et al. [2] | N_Set1_LB | ACTGAGGGAGCCTTGAATACA | 64.2 |
| Alves et al. [2] | N_Set2_F3 | TGGACCCCAAAATCAGCG | 61.7 |
| Alves et al. [2] | N_Set2_B3 | GCCTTGTCCTCGAGGGAAT | 64.6 |
| Alves et al. [2] | N_Set2_F1c | CCACTGCGTTCTCCATTCTGGT | 69.4 |
| Alves et al. [2] | N_Set2_F2 | AAATGCACCCCGCATTACG | 62.8 |
| Alves et al. [2] | N_Set2_B1c | CGCGATCAAAACAACGTCGGC | 70.1 |
| Alves et al. [2] | N_Set2_B2 | CCTTGCCATGTTGAGTGAGA | 62.9 |
| Alves et al. [2] | N_Set2_LF | TTGAATCTGAGGGTCCACCAAA | 66.4 |
| Alves et al. [2] | N_Set2_LB | GGTTTACCCAATAATACTGCGTCTT | 67.6 |
| Alves et al. [2] | RdRp_F3 | CTGTCAAATTACAGAATAATGAGC | 62.4 |

(continued on next page)

(Table S1 continued)

| Name of Set | Primer | Sequence 5′ → 3′ | $T_{\text{ref.}}$ (°C) |
|---|---|---|---|
| Alves et al. [2] | RdRp_B3 | TCCATCACTCTTAGGGAATC | 60.2 |
| Alves et al. [2] | RdRp_F1c | TGTCATCAGTGCAAGCAGTTTG | 65.7 |
| Alves et al. [2] | RdRp_F2 | CTGTTGCACTACGACAGA | 56.6 |
| Alves et al. [2] | RdRp_B1c | ATGCGTTAGCTTACTACAACACA | 63.4 |
| Alves et al. [2] | RdRp_B2 | CCCATTTCAAATCCTGTAAATCG | 64.6 |
| Alves et al. [2] | RdRp_LF | ACCGGCAGCACAAGACA | 60.2 |
| Alves et al. [2] | RdRp_LB | ACAAAGGGAGGTAGGTTTGTACT | 63.8 |
| Alves et al. [2] | E_Set1_F3 | TGATGAGCCTGAAGAACATG | 61.3 |
| Alves et al. [2] | E_Set1_B3 | CGCTATTAACTATTAACGTACCT | 59.5 |
| Alves et al. [2] | E_Set1_F1c | TCGGTTCATCATAAATTGGTTCCAT | 68.6 |
| Alves et al. [2] | E_Set1_F2 | CAAATTCACACAATCGACGG | 60.8 |
| Alves et al. [2] | E_Set1_B1c | ACGACTACTAGCGTGCCTTT | 62.5 |
| Alves et al. [2] | E_Set1_B2 | GTCTCTTCCGAAACGAATG | 59.7 |
| Alves et al. [2] | E_Set1_LF | ACTGGATTAACAACTCCGGATGA | 66.9 |
| Alves et al. [2] | E_Set1_LB | GTAAGCACAAGCTGATGAGTACGAA | 70.6 |
| Diego et al. [3] | ORF1b-F3 | CACAGACTTTGTGAATGAGTT | 58.0 |
| Diego et al. [3] | ORF1b-B3 | GTCAGTCTCAGTCCAACAT | 57.3 |
| Diego et al. [3] | ORF1b-F1c | CTATTGAAACACACAACAGCATCG | 66.7 |
| Diego et al. [3] | ORF1b-F2 | CATATTTGCGTAAACATTTCTCA | 60.8 |
| Diego et al. [3] | ORF1b-B1c | TATGCATCTCAAGGTCTAGTGGCTA | 70.9 |
| Diego et al. [3] | ORF1b-B2 | TGCTTCAGACATAAAAACATTG | 59.1 |
| Diego et al. [3] | S447-F3 | GTTTCTGCCTTTCCAACAA | 57.8 |
| Diego et al. [3] | S447-B3 | AACAGGGACTTCTGTGCA | 58.1 |
| Diego et al. [3] | S447-F1c | TCAAGAATCTCAAGTGTCTGTGG | 65.7 |
| Diego et al. [3] | S447-F2 | TGGCAGAGACATTGCTGA | 60.1 |
| Diego et al. [3] | S447-B1c | ACCATGTTCTTTTGGTGGTGTCAACAT | 72.5 |
| Diego et al. [3] | S447-B2 | CCTGATAAAGAACAGC | 49.2 |
| Diego et al. [3] | S447-LF | TCACGGACAGCATCAGTAGTG | 65.1 |
| Diego et al. [3] | S447-LB | CAGGAACAAATACTTCTAACCAGGT | 66.4 |
| Diego et al. [3] | S555-F3 | CTATGCAAATGGCTTATAGGTT | 60.3 |
| Diego et al. [3] | S555-B3 | AGTTGTTTAACAAGCGTGTT | 55.5 |
| Diego et al. [3] | S555-F1c | GCACTATTAAATTGGTTGGCAATCA | 67.7 |
| Diego et al. [3] | S555-F2 | TAATGGTATTGGAGTTACACAGA | 60.5 |
| Diego et al. [3] | S555-B1c | ATTGGCAAAATTCAAGACTCACTT | 65.0 |
| Diego et al. [3] | S555-B2 | TTGTGCATTTTGGTTGACC | 57.5 |
| Diego et al. [3] | E-F3 | TCATTCGTTTCGGAAGAGA | 60.0 |
| Diego et al. [3] | E-B3 | AGGAACTCTAGAAGAATTCAGAT | 62.1 |
| Diego et al. [3] | E-F1c | TGTAACTAGCAAGAATACCACGAAAC | 68.6 |
| Diego et al. [3] | E-F2 | AGGTACGTTAATAGTTAATAGCG | 59.5 |
| Diego et al. [3] | E-B1c | GCTTCGATTGTGTGCGTACT | 63.0 |
| Diego et al. [3] | E-B2 | CGAGAGTAAACGTAAAAAGAAGG | 63.1 |
| Diego et al. [3] | M-F3 | GTTTCCTATTCCTTACATGGATT | 61.2 |

(continued on next page)

(Table S1 continued)

| Name of Set | Primer | Sequence 5′ → 3′ | $T_{ref.}$ (°C) |
|---|---|---|---|
| Diego et al. [3] | M-B3 | AGCCACATCAAGCCTACA | 58.4 |
| Diego et al. [3] | M-F1c | CCATAACAGCCAGAGGAAAATTAAC | 67.9 |
| Diego et al. [3] | M-F2 | CTTCTACAATTTGCCTATGCC | 61.3 |
| Diego et al. [3] | M-B1c | AACTTTAGCTTGTTTTGTGCTTGC | 66.3 |
| Diego et al. [3] | M-B2 | ACAAGCCATTGCGATAGC | 59.5 |
| Diego et al. [3] | N5-F3 | CCAGAATGGAGAACGCAGTG | 64.7 |
| Diego et al. [3] | N5-B3 | CCGTCACCACCACGAATT | 59.9 |
| Diego et al. [3] | N5-F1c | AGCGGTGAACCAAGACGCAG | 67.9 |
| Diego et al. [3] | N5-F2 | GGCGCGATCAAAACAACG | 62.4 |
| Diego et al. [3] | N5-B1c | AATTCCCTCGAGGACAAGGCG | 69.8 |
| Diego et al. [3] | N5-B2 | AGCTCTTCGGTAGTAGCCAA | 63.1 |
| Diego et al. [3] | N5-LF | ATTATTGGGTAAACCTTGGGGC | 64.6 |
| Diego et al. [3] | N5-LB | ATTAACACCAATAGCAGTCCAGATG | 67.6 |
| Ganguli et al. [4] | Orf1a-P1-F3 | ATCTAGTTGTAATGGCCTACA | 58.1 |
| Ganguli et al. [4] | Orf1a-P1-B3 | ACAAGCACAGGTTGAGAT | 55.4 |
| Ganguli et al. [4] | Orf1a-P1-F1c | TGAGTTTTTCATAAACAGTGCCAAA | 66.1 |
| Ganguli et al. [4] | Orf1a-P1-F2 | CAGGTGGTGTTGTTCAGT | 55.3 |
| Ganguli et al. [4] | Orf1a-P1-B1c | AACCCGTCCTTGATTGGCTT | 64.2 |
| Ganguli et al. [4] | Orf1a-P1-B2 | AATTTAACAATTTCCCAACCGTC | 62.5 |
| Ganguli et al. [4] | Orf1a-P1-Loop-F | TGTTAGTTAGCCACTGCGAAGT | 65.1 |
| Ganguli et al. [4] | Orf1a-P2-F3 | AAACCCGTCCTTGATTGG | 58.1 |
| Ganguli et al. [4] | Orf1a-P2-B3 | CTTTAAGTTTAGCTCCACCAAT | 59.8 |
| Ganguli et al. [4] | Orf1a-P2-F1c | TCACAAGCACAGGTTGAGATAAATT | 67.4 |
| Ganguli et al. [4] | Orf1a-P2-F2 | GAAGGTGTAGAGTTTCTTAGAGAC | 64.4 |
| Ganguli et al. [4] | Orf1a-P2-B1c | GTCACCTGTGCAAAGGAAATTAAG | 66.7 |
| Ganguli et al. [4] | Orf1a-P2-B2 | AGAGTCAGCACACAAAGC | 57.5 |
| Ganguli et al. [4] | Orf1a-P3-F3 | CGGTGGACAAATTGTCAC | 56.7 |
| Ganguli et al. [4] | Orf1a-P3-B3 | CTTCTCTGGATTTAACACACTT | 59.3 |
| Ganguli et al. [4] | Orf1a-P3-F1c | TCAGCACACAAAGCCAAAAATTTAT | 66.9 |
| Ganguli et al. [4] | Orf1a-P3-F2 | CTGTGCAAAGGAAATTAAGGAG | 62.6 |
| Ganguli et al. [4] | Orf1a-P3-B1c | TATTGGTGGAGCTAAACTTAAAGCC | 68.6 |
| Ganguli et al. [4] | Orf1a-P3-B2 | CTGTACAATCCCTTTGAGTG | 58.0 |
| Ganguli et al. [4] | S-P1-F3 | TCTTTCACACGTGGTGTT | 55.2 |
| Ganguli et al. [4] | S-P1-B3 | CAGTGGAAGCAAAATAAACAC | 58.5 |
| Ganguli et al. [4] | S-P1-F1c | GAAAGGTAAGAACAAGTCCTGAGT | 65.7 |
| Ganguli et al. [4] | S-P1-F2 | TTACCCTGACAAAGTTTTCAG | 58.4 |
| Ganguli et al. [4] | S-P1-B1c | TTCCAATGTTACTTGGTTCCATGC | 67.9 |
| Ganguli et al. [4] | S-P1-B2 | GACAGGGTTATCAAACCTCT | 58.8 |
| Ganguli et al. [4] | S-P1-Loop-B | TATACATGTCTCTGGGACCAATGG | 68.1 |
| Ganguli et al. [4] | S-P2-F3 | GGTGTTTATTACCCTGACAAAG | 60.1 |
| Ganguli et al. [4] | S-P2-B3 | GTACCAAAAATCCAGCCTC | 58.3 |
| Ganguli et al. [4] | S-P2-F1c | TGGAACCAAGTAACATTGGAAAAGA | 67.5 |

(continued on next page)

(Table S1 continued)

| Name of Set | Primer | Sequence 5′ → 3′ | $T_{\text{ref.}}$ (°C) |
|---|---|---|---|
| Ganguli et al. [4] | S-P2-F2 | TTTTCAGATCCTCAGTTTTACATTC | 64.9 |
| Ganguli et al. [4] | S-P2-B1c | CTCTGGGACCAATGGTACTAAGAG | 68.7 |
| Ganguli et al. [4] | S-P2-B2 | GACTTCTCAGTGGAAGCA | 57.9 |
| Ganguli et al. [4] | S-P2-Loop-B | AACCCTGTCCTACCATTTAATGATG | 67.3 |
| Ganguli et al. [4] | S-P3-F3 | CCTGACAAAGTTTTCAGATCC | 61.0 |
| Ganguli et al. [4] | S-P3-B3 | GTACCAAAAATCCAGCCTC | 58.3 |
| Ganguli et al. [4] | S-P3-F1c | GCATGGAACCAAGTAACATTGGAAA | 69.3 |
| Ganguli et al. [4] | S-P3-F2 | TCAGTTTTACATTCAACTCAGGA | 62.4 |
| Ganguli et al. [4] | S-P3-B1c | CTCTGGGACCAATGGTACTAAGAG | 68.7 |
| Ganguli et al. [4] | S-P3-B2 | GACTTCTCAGTGGAAGCA | 57.9 |
| Ganguli et al. [4] | S-P3-Loop-B | AACCCTGTCCTACCATTTAATGATG | 67.3 |
| Ganguli et al. [4] | Orf8-P1-F3 | ACGCCTAAACGAACATGAA | 58.5 |
| Ganguli et al. [4] | Orf8-P1-B3 | AGAACCAGCCTCATCCAG | 60.0 |
| Ganguli et al. [4] | Orf8-P1-F1c | GGTTGATGTTGAGTACATGACTGTA | 66.1 |
| Ganguli et al. [4] | Orf8-P1-F2 | CTTGTTTTCTTAGGAATCATCACA | 63.3 |
| Ganguli et al. [4] | Orf8-P1-B1c | ATATGTAGTTGATGACCCGTGTCC | 67.7 |
| Ganguli et al. [4] | Orf8-P1-B2 | TAAAGGTGCTGATTTTCTAGCT | 61.8 |
| Ganguli et al. [4] | Orf8-P1-Loop-F | CTACATTCTTGGTGAAATGCAGCTA | 68.9 |
| Ganguli et al. [4] | Orf8-P2-F3 | CTCAACATCAACCATATGTAGT | 58.2 |
| Ganguli et al. [4] | Orf8-P2-B3 | CAATTTAGGTTCCTGGCAATT | 60.0 |
| Ganguli et al. [4] | Orf8-P2-F1c | GGTGCTGATTTTCTAGCTCCTACTC | 71.8 |
| Ganguli et al. [4] | Orf8-P2-F2 | GACCCGTGTCCTATTCAC | 58.0 |
| Ganguli et al. [4] | Orf8-P2-B1c | TGCTGGATGAGGCTGGTTCTA | 67.6 |
| Ganguli et al. [4] | Orf8-P2-B2 | TGTAAAAGGTAACAGGAAACTG | 58.6 |
| Ganguli et al. [4] | Orf8-P2-Loop-B | ATCACCCATTCAGTACATCGATATC | 68.1 |
| Ganguli et al. [4] | Orf8-P3-F3 | AGCTGCATTTCACCAAGAA | 59.2 |
| Ganguli et al. [4] | Orf8-P3-B3 | CGATATCGATGTACTGAATGG | 61.0 |
| Ganguli et al. [4] | Orf8-P3-F1c | TGAATAGGACACGGGTCATCA | 64.6 |
| Ganguli et al. [4] | Orf8-P3-F2 | GTAGTTTACAGTCATGTACTCAA | 58.2 |
| Ganguli et al. [4] | Orf8-P3-B1c | GAGTAGGAGCTAGAAAATCAGCAC | 68.6 |
| Ganguli et al. [4] | Orf8-P3-B2 | TGATTTAGAACCAGCCTCATC | 62.3 |
| Ganguli et al. [4] | N-P1-F3 | GTTCCTCATCACGTAGTCG | 59.8 |
| Ganguli et al. [4] | N-P1-B3 | GTTTGGCCTTGTTGTTGTT | 56.4 |
| Ganguli et al. [4] | N-P1-F1c | GCCAGCCATTCTAGCAGGAG | 67.3 |
| Ganguli et al. [4] | N-P1-F2 | CAACAGTTAAGAAATTCAACTCC | 60.6 |
| Ganguli et al. [4] | N-P1-B1c | GATGCTGCTCTTGCTTTGCT | 65.1 |
| Ganguli et al. [4] | N-P1-B2 | ACCAGACATTTTGCTCTCAA | 59.5 |
| Ganguli et al. [4] | N-P1-Loop-B | GCTGCTTGACAGATTGAACCAG | 67.4 |
| Ganguli et al. [4] | N-P2-F3 | AGACGAATTCGTGGTGGT | 58.3 |
| Ganguli et al. [4] | N-P2-B3 | TTGTTAGCAGGATTGCGG | 59.2 |
| Ganguli et al. [4] | N-P2-F1c | TGGCCCAGTTCCTAGGTAGT | 63.5 |
| Ganguli et al. [4] | N-P2-F2 | GACGGTAAAATGAAAGATCTCAG | 63.8 |

(continued on next page)

(Table S1 continued)

| Name of Set | Primer | Sequence 5′ → 3′ | $T_{\text{ref.}}$ (°C) |
|---|---|---|---|
| Ganguli et al. [4] | N-P2-B1c | CTTCCCTATGGTGCTAACAAAGAC | 67.7 |
| Ganguli et al. [4] | N-P2-B2 | TGGTGTATTCAAGGCTCC | 57.5 |
| Ganguli et al. [4] | N-P2-Loop-B | GGCATCATATGGGTTGCAACTGAG | 71.6 |
| Ganguli et al. [4] | N-P3-F3 | GTCATTTTGCTGAATAAGCATAT | 60.8 |
| Ganguli et al. [4] | N-P3-B3 | GAGTCAGCACTGCTCATG | 59.6 |
| Ganguli et al. [4] | N-P3-F1c | TAAGGCTTGAGTTTCATCAGCCTT | 69.3 |
| Ganguli et al. [4] | N-P3-F2 | ACGCATACAAAACATTCCCA | 59.5 |
| Ganguli et al. [4] | N-P3-B1c | CAGAGACAGAAGAAACAGCAAACT | 67.1 |
| Ganguli et al. [4] | N-P3-B2 | GATTGTTGCAATTGTTTGGAG | 59.9 |
| Ganguli et al. [4] | N-P3-Loop-B | GTGACTCTTCTTCCTGCTGCAGATT | 73.4 |
| Garcia-Venzor et al. [5] | N-geneF3 | AACACAAGCTTTCGGCAG | 58.9 |
| Garcia-Venzor et al. [5] | N-geneB3 | GAAATTTGGATCTTTGTCATCC | 61.5 |
| Garcia-Venzor et al. [5] | N-geneF1c | TGCGGCCAATGTTTGTAATCAG | 66.7 |
| Garcia-Venzor et al. [5] | N-geneF2 | CCAAGGAAATTTTGGGGAC | 59.2 |
| Garcia-Venzor et al. [5] | N-geneB1c | CGCATTGGCATGGAAGTCAC | 65.8 |
| Garcia-Venzor et al. [5] | N-geneB2 | TTTGATGGCACCTGTGTAG | 58.2 |
| Garcia-Venzor et al. [5] | N-geneLF | TTCCTTGTCTGATTAGTTC | 53.8 |
| Garcia-Venzor et al. [5] | N-geneLB | ACCTTCGGGAACGTGGTT | 60.5 |
| Huang et al. [6] | N1-F3 | TGGACCCCAAAATCAGCG | 61.7 |
| Huang et al. [6] | N1-B3 | GCCTTGTCCTCGAGGGAAT | 64.6 |
| Huang et al. [6] | N1-F1c | CCACTGCGTTCTCCATTCTGGT | 69.4 |
| Huang et al. [6] | N1-F2 | AAATGCACCCCGCATTACG | 62.8 |
| Huang et al. [6] | N1-B1c | CGCGATCAAAACAACGTCGGC | 70.1 |
| Huang et al. [6] | N1-B2 | CCTTGCCATGTTGAGTGAGA | 62.9 |
| Huang et al. [6] | N1-LF | TGAATCTGAGGGTCCACCAAA | 65.0 |
| Huang et al. [6] | N1-LB | GGTTTACCCAATAATACTGCGTCTT | 67.6 |
| Huang et al. [6] | N15-F3 | AGATCACATTGGCACCCG | 60.6 |
| Huang et al. [6] | N15-B3 | CCATTGCCAGCCATTCTAGC | 65.6 |
| Huang et al. [6] | N15-F1c | TGCTCCCTTCTGCGTAGAAGC | 69.5 |
| Huang et al. [6] | N15-F2 | CAATGCTGCAATCGTGCTAC | 63.3 |
| Huang et al. [6] | N15-B1c | GGCGGCAGTCAAGCCTCTTCCC | 76.6 |
| Huang et al. [6] | N15-B2 | TACTGCTGCCTGGAGTT | 56.7 |
| Huang et al. [6] | N15-LF | GCAATGTTGTTCCTTGAGGAAGTT | 67.9 |
| Huang et al. [6] | N15-LB | GTTCCTCATCACGTAGTCGCAACA | 71.7 |
| Huang et al. [6] | S17-F3 | TCTTTCACACGTGGTGTT | 55.2 |
| Huang et al. [6] | S17-B3 | GTACCAAAAATCCAGCCTC | 58.3 |
| Huang et al. [6] | S17-F1c | CATGGAACCAAGTAACATTGGAAAA | 66.6 |
| Huang et al. [6] | S17-F2 | CCTGACAAAGTTTTCAGATCC | 61.0 |
| Huang et al. [6] | S17-B1c | CTCTGGGACCAATGGTACTAAGAG | 68.7 |
| Huang et al. [6] | S17-B2 | GACTTCTCAGTGGAAGCA | 57.9 |
| Huang et al. [6] | S17-LF | GAAAGGTAAGAACAAGTCCTGAGT | 65.7 |
| Huang et al. [6] | S17-LB | CTGTCCTACCATTTAATGATGGTGT | 66.7 |

(continued on next page)

(Table S1 continued)

| Name of Set | Primer | Sequence 5′ → 3′ | $T_{\text{ref.}}$ (°C) |
|---|---|---|---|
| Huang et al. [6] | O117-F3 | CCCCAAAATGCTGTTGTT | 55.8 |
| Huang et al. [6] | O117-B3 | TAGCACGTGGAACCCAAT | 58.4 |
| Huang et al. [6] | O117-F1c | GGTTTTCAAGCCAGATTCATTATGG | 69.2 |
| Huang et al. [6] | O117-F2 | ATGTCACAATTCAGAAGTAGGA | 60.5 |
| Huang et al. [6] | O117-B1c | TCTTCGTAAGGGTGGTCGCA | 66.3 |
| Huang et al. [6] | O117-B2 | GCACACTTGTTATGGCAAC | 57.9 |
| Huang et al. [6] | O117-LF | TCGGCAAGACTATGCTCAGG | 65.8 |
| Huang et al. [6] | O117-LB | TTGCCTTTGGAGGCTGTGT | 62.8 |
| Jang et al. [7] | RdRP-F3 | CGATAAGTATGTCCGCAATT | 58.6 |
| Jang et al. [7] | RdRP-B3 | GCTTCAGACATAAAAACATTGT | 58.3 |
| Jang et al. [7] | RdRP-F1c | ATGCGTAAAACTCATTCACAAAGTC | 66.9 |
| Jang et al. [7] | RdRP-F2 | CAACACAGACTTTATGAGTGTC | 59.9 |
| Jang et al. [7] | RdRP-B1c | TGATACTCTCTGACGATGCTGTT | 67.0 |
| Jang et al. [7] | RdRP-B2 | TAAAGTTCTTTATGCTAGCCAC | 60.2 |
| Jang et al. [7] | RdRP-BLP | TCAATAGCACTTATGCATCTCAAGG | 69.3 |
| Jang et al. [7] | RdRP-FLP | TGTGTCAACATCTCTATTTCTATAG | 62.4 |
| Jang et al. [7] | E-F3 | TCATTCGTTTCGGAAGAGA | 60.0 |
| Jang et al. [7] | E-B3 | AGGAACTCTAGAAGAATTCAGAT | 62.1 |
| Jang et al. [7] | E-F1c | TGTAACTAGCAAGAATACCACGAAA | 66.6 |
| Jang et al. [7] | E-F2 | CAGGTACGTTAATAGTTAATAGCG | 62.4 |
| Jang et al. [7] | E-B1c | GCTTCGATTGTGTGCGT | 57.8 |
| Jang et al. [7] | E-B2 | ACTCGAGAGTAAACGTAAAAAGAAGG | 69.0 |
| Jang et al. [7] | E-BLP | GCTGCAATATTGTTAACGTGAGTC | 67.0 |
| Ji et al. [8] | N1-F3 | CCCCAAAATCAGCGAAATGC | 65.0 |
| Ji et al. [8] | N1-B3 | CCACCACGAATTCGTCTGG | 63.2 |
| Ji et al. [8] | N1-F1c | CGTTGTTTTGATCGCGCCCCTTT | 73.3 |
| Ji et al. [8] | N1-F2 | CATTACGTTTGGTGGACCCT | 61.2 |
| Ji et al. [8] | N1-B1c | AATTCCCTCGAGGACAAGGCGTTT | 73.9 |
| Ji et al. [8] | N1-B2 | TAGCTCTTCGGTAGTAGCCA | 62.7 |
| Ji et al. [8] | N1-LF | TGGTTACTGCCAGTTGAATC | 60.0 |
| Ji et al. [8] | N1-LB | TAACACCAATAGCAGTCCAGATG | 65.2 |
| Ji et al. [8] | N2-F3 | CACCCGCAATCCTGCTAAC | 63.5 |
| Ji et al. [8] | N2-B3 | TTTGCTCTCAAGCTGGTTCA | 62.9 |
| Ji et al. [8] | N2-F1c | CCTCTGCTCCCTTCTGCGTAGATTT | 75.6 |
| Ji et al. [8] | N2-F2 | AATGCTGCAATCGTGCTACA | 62.6 |
| Ji et al. [8] | N2-B1c | AACTCCAGGCAGCAGTAGGGTT | 69.0 |
| Ji et al. [8] | N2-B2 | TGTCAAGCAGCAGCAAAGC | 63.7 |
| Ji et al. [8] | N2-LF | GCCTTTTGGCAATGTTGTTCCTT | 67.9 |
| Ji et al. [8] | N2-LB | CTCCTGCTAGAATGGCTGGC | 67.3 |
| Ji et al. [8] | ORFlab-1-F3 | GCTGCACTTACTAACAATGT | 56.5 |
| Ji et al. [8] | ORFlab-1-B3 | GGTTGTTGACGATGACTTG | 58.0 |
| Ji et al. [8] | ORFlab-1-F1c | TCCTTCCTTAAAGAAACCCTTAGACTT | 71.0 |

(continued on next page)

(Table S1 continued)

| Name of Set | Primer | Sequence 5′ → 3′ | $T_{\text{ref.}}$ (°C) |
|---|---|---|---|
| Ji et al. [8] | ORFlab-1-F2 | TTTTCAAACTGTCAAACCCG | 58.8 |
| Ji et al. [8] | ORFlab-1-B1c | CCAACAATGTGTGATATCAGACAACT | 68.1 |
| Ji et al. [8] | ORFlab-1-B2 | TTAATACAGCCACCATCGTAAC | 61.8 |
| Ji et al. [8] | ORFlab-1-LF | GCAGCTACTGAAAAGCACGT | 63.0 |
| Ji et al. [8] | ORFlab-1-LB | TTGTAGTTGAGTTGTTGATAAGT | 58.0 |
| Ji et al. [8] | ORFlab-2-F3 | GACATCACATACAGTAATGCC | 59.4 |
| Ji et al. [8] | ORFlab-2-B3 | TGTATACACTATGCGAGCAG | 59.2 |
| Ji et al. [8] | ORFlab-2-F1c | CTCATCTGAGATATTGAGTGTTGGGT | 70.7 |
| Ji et al. [8] | ORFlab-2-F2 | ACCTACACTAGTGCCACA | 55.2 |
| Ji et al. [8] | ORFlab-2-B1c | AAGTATTCTACACTCCAGGGACC | 65.7 |
| Ji et al. [8] | ORFlab-2-B2 | GGTAGTAGAGAGCTAGGCC | 60.7 |
| Ji et al. [8] | ORFlab-2-LF | GCCAGTAATTCTAACATAGTGCT | 62.7 |
| Ji et al. [8] | ORFlab-2-LB | GGTACTGGTAAGAGTCATTTTGC | 64.2 |
| Jiang et al. [9] | nCoV-N-F3 | CCAGAATGGAGAACGCAGTG | 64.7 |
| Jiang et al. [9] | nCoV-N-B3 | CCGTCACCACCACGAATT | 59.9 |
| Jiang et al. [9] | nCoV-N-F1c | AGCGGTGAACCAAGACGCAG | 67.9 |
| Jiang et al. [9] | nCoV-N-F2 | GGCGCGATCAAAACAACG | 62.4 |
| Jiang et al. [9] | nCoV-N-B1c | AATTCCCTCGAGGACAAGGCG | 69.8 |
| Jiang et al. [9] | nCoV-N-B2 | AGCTCTTCGGTAGTAGCCAA | 63.1 |
| Jiang et al. [9] | nCoV-N-LF | TTATTGGGTAAACCTTGGGGC | 63.6 |
| Jiang et al. [9] | nCoV-N-LB | TTCCAATTAACACCAATAGCAGTCC | 68.4 |
| Lalli et al. [10] | NEB_orf1a-A-F3 | CTGCACCTCATGGTCATGTT | 62.1 |
| Lalli et al. [10] | NEB_orf1a-A-B3 | AGCTCGTCGCCTAAGTCAA | 63.1 |
| Lalli et al. [10] | NEB_orf1a-A-F1c | GAGGGACAAGGACACCAAGTGTA | 68.4 |
| Lalli et al. [10] | NEB_orf1a-A-F2 | TGGTTGAGCTGGTAGCAGA | 62.2 |
| Lalli et al. [10] | NEB_orf1a-A-B1c | CCAGTGGCTTACCGCAAGGTT | 68.0 |
| Lalli et al. [10] | NEB_orf1a-A-B2 | TTAGATCGGCGCCGTAAC | 60.7 |
| Lalli et al. [10] | NEB_orf1a-A-LF | CCGTACTGAATGCCTTCGAGT | 66.1 |
| Lalli et al. [10] | NEB_orf1a-A-LB | TTCGTAAGAACGGTAATAAAGGAGC | 68.9 |
| Lalli et al. [10] | NEB_geneN-A-F3 | TGGCTACTACCGAAGAGCT | 61.6 |
| Lalli et al. [10] | NEB_geneN-A-B3 | TGCAGCATTGTTAGCAGGAT | 62.5 |
| Lalli et al. [10] | NEB_geneN-A-F1c | TCTGGCCCAGTTCCTAGGTAGT | 68.1 |
| Lalli et al. [10] | NEB_geneN-A-F2 | CCAGACGAATTCGTGGTGG | 63.2 |
| Lalli et al. [10] | NEB_geneN-A-B1c | AGACGGCATCATATGGGTTGCA | 69.2 |
| Lalli et al. [10] | NEB_geneN-A-B2 | CGGGTGCCAATGTGATCT | 60.6 |
| Lalli et al. [10] | NEB_geneN-A-LF | GGACTGAGATCTTTCATTTTACCGT | 68.6 |
| Lalli et al. [10] | NEB_geneN-A-LB | ACTGAGGGAGCCTTGAATACA | 64.2 |
| Lalli et al. [10] | NEB_N2-F3 | ACCAGGAACTAATCAGACAAG | 59.8 |
| Lalli et al. [10] | NEB_N2-B3 | GACTTGATCTTTGAAATTTGGATCT | 65.7 |
| Lalli et al. [10] | NEB_N2-F1c | TTCCGAAGAACGCTGAAGCG | 67.7 |
| Lalli et al. [10] | NEB_N2-F2 | GAACTGATTACAAACATTGGCC | 62.4 |
| Lalli et al. [10] | NEB_N2-B1c | CGCATTGGCATGGAAGTCAC | 65.8 |

(continued on next page)

(Table S1 continued)

| Name of Set | Primer | Sequence 5′ → 3′ | $T_{ref.}$ (°C) |
|---|---|---|---|
| Lalli et al. [10] | NEB_N2-B2 | AATTTGATGGCACCTGTGTA | 58.5 |
| Lalli et al. [10] | NEB_N2-LF | GGGGGCAAATTGTGCAATTTG | 65.4 |
| Lalli et al. [10] | NEB_N2-LB | CTTCGGGAACGTGGTTGACC | 66.4 |
| Lalli et al. [10] | NEB_E1-F3 | TGAGTACGAACTTATGTACTCAT | 60.1 |
| Lalli et al. [10] | NEB_E1-B3 | TTCAGATTTTTAACACGAGAGT | 59.2 |
| Lalli et al. [10] | NEB_E1-F1c | ACCACGAAAGCAAGAAAAAGAAG | 65.4 |
| Lalli et al. [10] | NEB_E1-F2 | TTCGTTTCGGAAGAGACAG | 60.2 |
| Lalli et al. [10] | NEB_E1-B1c | TTGCTAGTTACACTAGCCATCCTT | 66.9 |
| Lalli et al. [10] | NEB_E1-B2 | AGGTTTTACAAGACTCACGT | 56.8 |
| Lalli et al. [10] | NEB_E1-LB | GCGCTTCGATTGTGTGCGT | 65.2 |
| Lalli et al. [10] | NEB_E1-LF | CGCTATTAACTATTAACG | 48.4 |
| Lau et al. [11] | F1c | TGGGGTCCATTATCAGACATTTTAGT | 69.7 |
| Lau et al. [11] | F2 | TTTAGAGTATCATGACGTTCG | 59.2 |
| Lau et al. [11] | B1c | CGAAATGCACCCCGCATTAC | 65.8 |
| Lau et al. [11] | B2 | CCACTGCGTTCTCCATTC | 60.2 |
| Lau et al. [11] | FLP | TGTTCGTTTAGATGAAATC | 52.7 |
| Lau et al. [11] | BLP | TGGTGGACCCTCAGATTCAA | 63.6 |
| Lau et al. [11] | FP | GTTGTTCGTTCTATGAAGACT | 58.3 |
| Lau et al. [11] | BP | GACGTTGTTTTGATCGCG | 59.2 |
| Lau et al. [11] | FSP | TTGGGGTCCATTATCAG | 53.7 |
| Lau et al. [11] | BSP | ATCAGCGAAATGCACC | 54.6 |
| Luo et al. [12] | PS1-F3 | CACCCGCAATCCTGCTAAC | 63.5 |
| Luo et al. [12] | PS1-B3 | CAGCCATTCTAGCAGGAGA | 62.0 |
| Luo et al. [12] | PS1-F1c | TGCTCCCTTCTGCGTAGAAGC | 69.5 |
| Luo et al. [12] | PS1-F2 | AATGCTGCAATCGTGCTACA | 62.6 |
| Luo et al. [12] | PS1-B1c | GGCGGCAGTCAAGCCTCTT | 67.2 |
| Luo et al. [12] | PS1-B2 | CCCTACTGCTGCCTGGAGTT | 66.1 |
| Luo et al. [12] | PS1-LF | GCAATGTTGTTCCTTGAGGAAGT | 66.5 |
| Luo et al. [12] | PS1-LB | TTCCTCATCACGTAGTCGCAACAGT | 73.3 |
| Luo et al. [12] | PS2-F3 | CCAGAATGGAGAACGCAGTG | 64.7 |
| Luo et al. [12] | PS2-B3 | CCGTCACCACCACGAATT | 59.9 |
| Luo et al. [12] | PS2-F1c | AGCGGTGAACCAAGACGCAG | 67.9 |
| Luo et al. [12] | PS2-F2 | GGCGCGATCAAAACAACG | 62.4 |
| Luo et al. [12] | PS2-B1c | AATTCCCTCGAGGACAAGGCG | 69.8 |
| Luo et al. [12] | PS2-B2 | AGCTCTTCGGTAGTAGCCAA | 63.1 |
| Luo et al. [12] | PS2-LF | TTATTGGGTAAACCTTGGGGC | 63.6 |
| Luo et al. [12] | PS2-LB | TCCAATTAACACCAATAGCAGTCCA | 69.2 |
| Luo et al. [12] | PS3-F3 | CCAGAATGGAGAACGCAGTG | 64.7 |
| Luo et al. [12] | PS3-B3 | CCGTCACCACCACGAATT | 59.9 |
| Luo et al. [12] | PS3-F1c | AGCGGTGAACCAAGACGCAG | 67.9 |
| Luo et al. [12] | PS3-F2 | GGCGCGATCAAAACAACG | 62.4 |
| Luo et al. [12] | PS3-B1c | AATTCCCTCGAGGACAAGGCG | 69.8 |

(continued on next page)

(Table S1 continued)

| Name of Set | Primer | Sequence 5′ → 3′ | $T_{\text{ref.}}$ (°C) |
|---|---|---|---|
| Luo et al. [12] | PS3-B2 | AGCTCTTCGGTAGTAGCCAA | 63.1 |
| Luo et al. [12] | PS3-LF | ATTATTGGGTAAACCTTGGGGC | 64.6 |
| Luo et al. [12] | PS3-LB | CCAATTAACACCAATAGCAGTCCA | 66.8 |
| Luo et al. [12] | PS4-F3 | AGATCACATTGGCACCCG | 60.6 |
| Luo et al. [12] | PS4-B3 | CCATTGCCAGCCATTCTAGC | 65.6 |
| Luo et al. [12] | PS4-F1c | TGCTCCCTTCTGCGTAGAAGC | 69.5 |
| Luo et al. [12] | PS4-F2 | CAATGCTGCAATCGTGCTAC | 63.3 |
| Luo et al. [12] | PS4-B1c | GGCGGCAGTCAAGCCTCTT | 67.2 |
| Luo et al. [12] | PS4-B2 | CCCTACTGCTGCCTGGAGTT | 66.1 |
| Luo et al. [12] | PS4-LF | GCAATGTTGTTCCTTGAGGAAGTT | 67.9 |
| Luo et al. [12] | PS4-LB | GTTCCTCATCACGTAGTCGCAACA | 71.7 |
| Luo et al. [12] | PS5-F3 | AGATCACATTGGCACCCG | 60.7 |
| Luo et al. [12] | PS5-B3 | CCATTGCCAGCCATTCTAGC | 65.6 |
| Luo et al. [12] | PS5-F1c | TGCTCCCTTCTGCGTAGAAGC | 69.5 |
| Luo et al. [12] | PS5-F2 | CAATGCTGCAATCGTGCTAC | 63.3 |
| Luo et al. [12] | PS5-B1c | GGCGGCAGTCAAGCCTCTT | 67.2 |
| Luo et al. [12] | PS5-B2 | CCCTACTGCTGCCTGGAGTT | 66.1 |
| Luo et al. [12] | PS5-LF | GGCAATGTTGTTCCTTGAGGAAGTT | 71.1 |
| Luo et al. [12] | PS5-LB | GTTCCTCATCACGTAGTCGCAACA | 71.7 |
| Luo et al. [12] | PS6-F3 | TGGACCCCAAAATCAGCG | 61.7 |
| Luo et al. [12] | PS6-B3 | GCCTTGTCCTCGAGGGAAT | 64.6 |
| Luo et al. [12] | PS6-F1c | CCACTGCGTTCTCCATTCTGGT | 69.4 |
| Luo et al. [12] | PS6-F2 | AAATGCACCCCGCATTACG | 62.8 |
| Luo et al. [12] | PS6-B1c | CGCGATCAAAACAACGTCGGC | 70.1 |
| Luo et al. [12] | PS6-B2 | CCTTGCCATGTTGAGTGAGA | 62.9 |
| Luo et al. [12] | PS6-LF | TTGAATCTGAGGGTCCACCA | 63.6 |
| Luo et al. [12] | PS6-LB | TACCCAATAATACTGCGTCTTGGT | 67.0 |
| Mautner et al. [13] | ORF8-F3 | ACTTGTCACGCCTAAACG | 57.2 |
| Mautner et al. [13] | ORF8-B3 | CTACCCAATTTAGGTTCCTGG | 61.5 |
| Mautner et al. [13] | OFR8-F1c | AGGACACGGGTCATCAACTACA | 66.4 |
| Mautner et al. [13] | OFR8-F2 | AGCTGCATTTCACCAAGAA | 59.2 |
| Mautner et al. [13] | ORF8-B1c | AGGAGCTAGAAAATCAGCACCT | 66.0 |
| Mautner et al. [13] | OFR8-B2 | ATGGGTGATTTAGAACCAGC | 60.1 |
| Mautner et al. [13] | ORF8-LF | TGGTTGATGTTGAGTACATGAC | 61.2 |
| Mautner et al. [13] | ORF8-LB | AATTGAATTGTGCGTGGATGAG | 64.4 |
| Mohon et al. [14] | S2-F3 | ATTCTAAGCACACGCCTAT | 57.3 |
| Mohon et al. [14] | S2-B3 | GAAGATAACCCACATAATAAGCT | 60.8 |
| Mohon et al. [14] | S2-F1c | ACCTATTGGCAAATCTACCAATGG | 67.0 |
| Mohon et al. [14] | S2-F2 | TTTAGTGCGTGATCTCCCT | 60.1 |
| Mohon et al. [14] | S2-B1c | ATCACTAGGTTTCAAACTTTACTTGC | 66.7 |
| Mohon et al. [14] | S2-B2 | CTGTCCAACCTGAAGAAGA | 58.6 |
| Mohon et al. [14] | S2-LPF | TTCTAAAGCCGAAAAACCCTG | 63.0 |

(continued on next page)

(Table S1 continued)

| Name of Set | Primer | Sequence 5′ → 3′ | $T_{ref.}$ (°C) |
|---|---|---|---|
| Mohon et al. [14] | S2-LPB | CATAGAAGTTATTTGACTCCTGGTG | 66.3 |
| Mohon et al. [14] | S3-F3 | CACCTTATGGGTTGGGATT | 58.0 |
| Mohon et al. [14] | S3-B3 | AACATATAGTGAACCGCCA | 57.0 |
| Mohon et al. [14] | S3-F1c | GTTTGCGAGCAAGAACAAGTG | 64.4 |
| Mohon et al. [14] | S3-F2 | AATGTGATAGAGCCATGCC | 59.7 |
| Mohon et al. [14] | S3-B1c | ATACAACGTGTTGTAGCTTGTCA | 63.2 |
| Mohon et al. [14] | S3-B2 | CACATGACCATTTCACTCAA | 57.9 |
| Mohon et al. [14] | S3-LPF | GGCCATAATTCTAAGCATGTTA | 61.1 |
| Mohon et al. [14] | S3-LPB | ATTAGCTAATGAGTGTGCTCAAGTA | 66.2 |
| Rodriguez-Manzano et al. [15] | F3-N | ACCAATAGCAGTCCAGATGA | 60.8 |
| Rodriguez-Manzano et al. [15] | B3-N | CACGATTGCAGCATTGTTAGC | 64.8 |
| Rodriguez-Manzano et al. [15] | F1c-N | TCTGGCCCAGTTCCTAGGTAGT | 68.1 |
| Rodriguez-Manzano et al. [15] | F2-N | CCAGACAAATTCGTGGTGG | 60.5 |
| Rodriguez-Manzano et al. [15] | B1c-N | GGACTTCCCTATGGTGCTAACAAA | 68.8 |
| Rodriguez-Manzano et al. [15] | B2-N | CGGGTGCCAATGTGATCT | 60.6 |
| Rodriguez-Manzano et al. [15] | LF-N | GGACTGAGATCTTTCATTTTACCGT | 68.6 |
| Rodriguez-Manzano et al. [15] | LB-N | ACTGAGGGAGCCTTGAATACA | 64.2 |
| Yan et al. [16] | orf1ab-4F3 | GGTATGATTTTGTAGAAAACCCA | 61.2 |
| Yan et al. [16] | orf1ab-4B3 | CAACAGGAACTCCACTACC | 58.2 |
| Yan et al. [16] | orf1ab-4F1c | GGCATCACAGAATTGTACTGTTTTT | 66.2 |
| Yan et al. [16] | orf1ab-4F2 | GCGTATACGCCAACTTAGG | 60.0 |
| Yan et al. [16] | orf1ab-4B1c | AATGCTGGTATTGTTGGTGTACTGA | 67.8 |
| Yan et al. [16] | orf1ab-4B2 | GGTTTGTATGAAATCACCGAA | 59.9 |
| Yan et al. [16] | orf1ab-4LF | AACAAAGCTTGGCGTACACGTTCA | 71.0 |
| Yan et al. [16] | S-123F3 | TCTATTGCCATACCCACAA | 57.2 |
| Yan et al. [16] | S-123B3 | GGTGTTTTGTAAATTTGTTTGAC | 58.2 |
| Yan et al. [16] | S-123F1c | CATTCAGTTGAATCACCACAAATGT | 66.2 |
| Yan et al. [16] | S-123F2 | GTGTTACCACAGAAATTCTACC | 59.7 |
| Yan et al. [16] | S-123B1c | GTTGCAATATGGCAGTTTTTGTACA | 66.7 |
| Yan et al. [16] | S-123B2 | TTGGGTGTTTTTGTCTTGTT | 56.1 |
| Yan et al. [16] | S-123LF | ACTGATGTCTTGGTCATAGACACT | 65.7 |
| Yan et al. [16] | S-123LB | TAAACCGTGCTTTAACTGGAATAGC | 68.9 |
| Yang et al. [17] | CU-N2-F3 | CGGCAGTCAAGCCTCTTC | 63.0 |
| Yang et al. [17] | CU-N2-B3 | TTGCTCTCAAGCTGGTTCAA | 62.9 |
| Yang et al. [17] | CU-N2-Loop-B | ATGGCGGTGATGCTGCTCTT | 67.5 |
| Yang et al. [17] | CU-N2-F1c | TCCCCTACTGCTGCCTGGAG | 68.9 |
| Yang et al. [17] | CU-N2-F2 | CGTTCCTCATCACGTAGTCG | 63.8 |
| Yang et al. [17] | CU-N2-B1c | TCTCCTGCTAGAATGGCTGGC | 69.5 |
| Yang et al. [17] | CU-N2-B2 | ATCTGTCAAGCAGCAGCAAAG | 65.3 |
| Yang et al. [17] | ORF1e-F3 | GGCTAACTAACATCTTTGGC | 59.5 |
| Yang et al. [17] | ORF1e-B3 | GTCAGCACACAAAGCCAA | 58.4 |
| Yang et al. [17] | ORF1e-Loop-F | TCTTCAAGCCAATCAAGGAC | 61.6 |

(continued on next page)

(Table S1 continued)

| Name of Set | Primer | Sequence 5′ → 3′ | $T_{\text{ref.}}$ (°C) |
|---|---|---|---|
| Yang et al. [17] | ORF1e-Loop-B | TTGTCGGTGGACAAATTGT | 57.6 |
| Yang et al. [17] | ORF1e-F1c | TCTCTAAGAAACTCTACACCTTCCT | 67.4 |
| Yang et al. [17] | ORF1e-F2 | ACTGTTTATGAAAAACTCAAACC | 58.7 |
| Yang et al. [17] | ORF1e-B1c | TATCTCAACCTGTGCTTGTGAAAA | 66.4 |
| Yang et al. [17] | ORF1e-B2 | GAATGTCTGAACACTCTCCT | 59.2 |
| Yoshikawa et al. [18] | LAMP_ORF1b-1_F3 | AACCTGAGTTTTATGAGGCT | 58.2 |
| Yoshikawa et al. [18] | LAMP_ORF1b-1_B3 | TCCTAAGTAAAGTTGAGTCACA | 59.4 |
| Yoshikawa et al. [18] | LAMP_ORF1b-1_F1c | TGCAAGCACCACATCTTAATGAAGT | 69.9 |
| Yoshikawa et al. [18] | LAMP_ORF1b-1_F2 | CGCATACAGTCTTACAGGCT | 61.2 |
| Yoshikawa et al. [18] | LAMP_ORF1b-1_B1c | ACGACCATGTCATATCAACATCACA | 68.7 |
| Yoshikawa et al. [18] | LAMP_ORF1b-1_B2 | ACATCACAACCTGGAGCAT | 59.3 |
| Yoshikawa et al. [18] | LAMP_ORF1b-1_LF | CAAAGAACACAAGCCCCAAC | 62.0 |
| Yoshikawa et al. [18] | LAMP_ORF1b-1_LB | GTCTTGTCTGTTAATCCGTATGTTTG | 67.9 |
| Yoshikawa et al. [18] | LAMP_ORF1b-2_F3 | GGTTTTTTCACTTACATTTGTGG | 60.5 |
| Yoshikawa et al. [18] | LAMP_ORF1b-2_B3 | TCCTCCAAAATATGTAATTTGCA | 61.7 |
| Yoshikawa et al. [18] | LAMP_ORF1b-2_F1c | GCGAAGTGTCCCATGAGCTTA | 67.0 |
| Yoshikawa et al. [18] | LAMP_ORF1b-2_F2 | TAAACTAGCTCTTGGAGGTTCCG | 68.1 |
| Yoshikawa et al. [18] | LAMP_ORF1b-2_B1c | AATGCGTCATCATCTGAAGCAT | 66.0 |
| Yoshikawa et al. [18] | LAMP_ORF1b-2_B2 | TTTCATAACCATCTATTTGTTCGCG | 68.3 |
| Yoshikawa et al. [18] | LAMP_ORF1b-2_LF | TCAGCATTCCAAGAATGTTCTGT | 65.8 |
| Yoshikawa et al. [18] | LAMP_ORF1b-2_LB | ATTGGATGTAATTATCTTGGCAAACC | 68.2 |

Table S2: Matched and mismatched coverages for Alekseenko et al. [1] set.

| Primer | Gene | SARS-CoV-2 | | Alpha | | Beta | | Gamma | |
|---|---|---|---|---|---|---|---|---|---|
| | | $C_{\text{strict}}$ | $C_{\text{part.}}$ | $C_{\text{strict}}$ | $C_{\text{part.}}$ | $C_{\text{strict}}$ | $C_{\text{part.}}$ | $C_{\text{strict}}$ | $C_{\text{part.}}$ |
| As1_B1c | Orf1a | 99.3 | 99.3 | 99.4 | 99.5 | 98.6 | 98.7 | 99.7 | 99.7 |
| As1_B2 | Orf1a | 96.0 | 96.0 | 99.5 | 99.5 | 98.8 | 98.8 | 99.8 | 99.8 |
| As1_B3 | Orf1a | 99.1 | 99.1 | 99.4 | 99.4 | 97.9 | 97.9 | 99.7 | 99.7 |
| As1_F1c | Orf1a | 0.000 | 99.3 | 0.000 | 99.4 | 0.000 | 98.7 | 0.000 | 99.7 |
| As1_F2 | Orf1a | 99.1 | 99.1 | 99.0 | 99.0 | 98.5 | 98.5 | 99.8 | 99.8 |
| As1_F3 | Orf1a | 99.3 | 99.3 | 99.4 | 99.4 | 98.2 | 98.2 | 99.6 | 99.6 |
| As1_LB | Orf1a | 99.1 | 99.3 | 99.4 | 99.6 | 98.7 | 98.8 | 99.7 | 99.7 |
| As1_LF | Orf1a | 98.7 | 99.2 | 99.3 | 99.5 | 98.4 | 98.5 | 98.9 | 99.8 |
| As1e_B1c | Orf1a | 99.3 | 99.3 | 99.4 | 99.5 | 98.6 | 98.7 | 99.7 | 99.8 |
| As1e_B2 | Orf1a | 96.0 | 96.1 | 99.5 | 99.5 | 98.8 | 98.8 | 99.8 | 99.8 |
| As1e_F1c | Orf1a | 0.000 | 99.3 | 0.000 | 99.5 | 0.000 | 98.8 | 0.000 | 99.8 |
| As1e_F2 | Orf1a | 99.1 | 99.1 | 98.9 | 98.9 | 98.5 | 98.5 | 99.8 | 99.8 |
| iLACO-B1c | Orf1ab | 99.4 | 99.5 | 99.8 | 99.8 | 99.1 | 99.1 | 99.5 | 99.5 |
| iLACO-B2 | Orf1ab | 98.9 | 98.9 | 99.6 | 99.6 | 99.0 | 99.0 | 99.4 | 99.4 |
| iLACO-B3 | Orf1ab | 99.3 | 99.3 | 99.7 | 99.7 | 99.0 | 99.0 | 99.4 | 99.4 |
| iLACO-F1c | Orf1ab | 0.000 | 99.5 | 0.000 | 99.7 | 0.000 | 99.1 | 0.000 | 99.5 |
| iLACO-F2 | Orf1ab | 99.5 | 99.5 | 99.8 | 99.8 | 99.2 | 99.2 | 99.6 | 99.6 |
| iLACO-F3 | Orf1ab | 99.4 | 99.4 | 99.8 | 99.8 | 98.7 | 98.7 | 99.5 | 99.5 |
| iLACO-LB | Orf1ab | 97.1 | 97.1 | 99.6 | 99.6 | 99.0 | 99.0 | 99.5 | 99.5 |
| iLACO-LF | Orf1ab | 99.1 | 99.4 | 99.5 | 99.8 | 99.0 | 99.1 | 98.8 | 99.6 |

Table S3: Matched and mismatched coverages for Alekseenko et al. [1] set.

| Primer | Gene | Delta | | Lambda | | Mu | | Omicron | |
|---|---|---|---|---|---|---|---|---|---|
| | | $C_{\text{strict}}$ | $C_{\text{part.}}$ | $C_{\text{strict}}$ | $C_{\text{part.}}$ | $C_{\text{strict}}$ | $C_{\text{part.}}$ | $C_{\text{strict}}$ | $C_{\text{part.}}$ |
| As1_B1c | Orf1a | 99.9 | 100. | 99.8 | 99.9 | 99.8 | 99.9 | 98.1 | 98.5 |
| As1_B2 | Orf1a | 98.7 | 98.7 | 99.7 | 99.7 | 99.9 | 99.9 | 98.6 | 98.6 |
| As1_B3 | Orf1a | 99.9 | 99.9 | 99.5 | 99.5 | 97.7 | 97.7 | 98.7 | 98.7 |
| As1_F1c | Orf1a | 0.000 | 99.2 | 0.000 | 99.1 | 0.000 | 99.7 | 0.000 | 98.6 |
| As1_F2 | Orf1a | 99.9 | 99.9 | 99.3 | 99.3 | 99.8 | 99.8 | 98.5 | 98.5 |
| As1_F3 | Orf1a | 99.9 | 99.9 | 99.5 | 99.5 | 99.7 | 99.7 | 98.4 | 98.4 |
| As1_LB | Orf1a | 99.3 | 99.9 | 98.3 | 99.9 | 99.5 | 99.9 | 98.6 | 98.7 |
| As1_LF | Orf1a | 99.8 | 99.9 | 99.4 | 99.7 | 99.7 | 99.8 | 98.5 | 98.5 |
| As1e_B1c | Orf1a | 99.9 | 100. | 99.8 | 99.9 | 99.8 | 99.9 | 98.1 | 98.5 |
| As1e_B2 | Orf1a | 98.7 | 98.7 | 99.7 | 99.7 | 99.9 | 99.9 | 98.6 | 98.6 |
| As1e_F1c | Orf1a | 0.000 | 100. | 0.000 | 99.4 | 0.000 | 99.9 | 0.000 | 98.6 |
| As1e_F2 | Orf1a | 99.9 | 99.9 | 99.2 | 99.2 | 99.8 | 99.8 | 98.4 | 98.4 |
| iLACO-B1c | Orf1ab | 100. | 100. | 100. | 100. | 100. | 100. | 99.6 | 99.6 |
| iLACO-B2 | Orf1ab | 100. | 100. | 99.9 | 99.9 | 99.8 | 99.8 | 90.3 | 90.3 |
| iLACO-B3 | Orf1ab | 99.9 | 99.9 | 99.7 | 99.7 | 99.9 | 99.9 | 90.3 | 90.3 |
| iLACO-F1c | Orf1ab | 0.000 | 100. | 0.000 | 100. | 0.000 | 100. | 0.000 | 99.5 |
| iLACO-F2 | Orf1ab | 99.9 | 99.9 | 99.7 | 99.7 | 99.9 | 99.9 | 99.0 | 99.0 |
| iLACO-F3 | Orf1ab | 100. | 100. | 99.8 | 99.8 | 99.9 | 99.9 | 98.2 | 98.2 |
| iLACO-LB | Orf1ab | 99.9 | 99.9 | 99.7 | 99.7 | 90.0 | 90.2 | 99.6 | 99.6 |
| iLACO-LF | Orf1ab | 99.7 | 99.9 | 99.4 | 100. | 99.5 | 99.9 | 0.484 | 99.0 |

Table S4: Matched and mismatched coverages for Alekseenko et al. [1] set.

| Primer | Gene | BA.2 | | BA.3 | | BA.4 | | BA.5 | |
|---|---|---|---|---|---|---|---|---|---|
| | | $C_{\text{strict}}$ | $C_{\text{part.}}$ | $C_{\text{strict}}$ | $C_{\text{part.}}$ | $C_{\text{strict}}$ | $C_{\text{part.}}$ | $C_{\text{strict}}$ | $C_{\text{part.}}$ |
| As1_B1c | Orf1a | 99.9 | 100. | 99.1 | 99.1 | 99.7 | 99.7 | 99.8 | 99.8 |
| As1_B2 | Orf1a | 100. | 100. | 99.1 | 99.1 | 100. | 100. | 99.5 | 99.5 |
| As1_B3 | Orf1a | 99.8 | 99.8 | 99.1 | 99.1 | 99.8 | 99.8 | 99.6 | 99.6 |
| As1_F1c | Orf1a | 0.000 | 100. | 0.000 | 98.6 | 0.000 | 99.5 | 0.000 | 99.7 |
| As1_F2 | Orf1a | 100. | 100. | 98.9 | 98.9 | 99.8 | 99.8 | 99.7 | 99.7 |
| As1_F3 | Orf1a | 99.7 | 99.7 | 98.6 | 98.6 | 99.2 | 99.2 | 99.5 | 99.5 |
| As1_LB | Orf1a | 99.9 | 100. | 98.9 | 98.9 | 99.7 | 99.8 | 99.6 | 99.6 |
| As1_LF | Orf1a | 99.9 | 99.9 | 99.1 | 99.1 | 99.5 | 99.7 | 99.2 | 99.8 |
| As1e_B1c | Orf1a | 99.9 | 100. | 99.1 | 99.1 | 99.7 | 99.7 | 99.8 | 99.8 |
| As1e_B2 | Orf1a | 99.9 | 99.9 | 99.1 | 99.1 | 100. | 100. | 99.5 | 99.5 |
| As1e_F1c | Orf1a | 0.000 | 100. | 0.000 | 99.1 | 0.000 | 99.8 | 0.000 | 99.8 |
| As1e_F2 | Orf1a | 100. | 100. | 98.9 | 98.9 | 99.8 | 99.8 | 99.7 | 99.7 |
| iLACO-B1c | Orf1ab | 100. | 100. | 99.1 | 100. | 100. | 100. | 100. | 100. |
| iLACO-B2 | Orf1ab | 100. | 100. | 98.9 | 98.9 | 99.4 | 99.4 | 99.8 | 99.8 |
| iLACO-B3 | Orf1ab | 99.9 | 99.9 | 98.6 | 98.6 | 99.7 | 99.7 | 95.0 | 95.0 |
| iLACO-F1c | Orf1ab | 0.000 | 100. | 0.000 | 100. | 0.000 | 100. | 0.000 | 99.9 |
| iLACO-F2 | Orf1ab | 100. | 100. | 100. | 100. | 99.8 | 99.8 | 99.3 | 99.3 |
| iLACO-F3 | Orf1ab | 100. | 100. | 99.7 | 99.7 | 100. | 100. | 100. | 100. |
| iLACO-LB | Orf1ab | 100. | 100. | 100. | 100. | 99.8 | 99.8 | 95.0 | 95.0 |
| iLACO-LF | Orf1ab | 99.2 | 100. | 98.9 | 99.4 | 99.7 | 100. | 97.8 | 98.1 |

Table S5: Matched and mismatched coverages for Alves et al. [2] set.

| Primer | Gene Target | SARS-CoV-2 | | Alpha | | Beta | | Gamma | |
|---|---|---|---|---|---|---|---|---|---|
| | | $C_{strict}$ | $C_{part.}$ | $C_{strict}$ | $C_{part.}$ | $C_{strict}$ | $C_{part.}$ | $C_{strict}$ | $C_{part.}$ |
| E_Set1_B1c | E | 99.2 | 99.2 | 99.4 | 99.4 | 98.1 | 98.1 | 99.6 | 99.6 |
| E_Set1_B2 | E | 98.5 | 98.5 | 98.9 | 98.9 | 98.5 | 98.5 | 99.9 | 99.9 |
| E_Set1_B3 | E | 99.5 | 99.5 | 99.7 | 99.7 | 99.4 | 99.4 | 99.9 | 99.9 |
| E_Set1_F1c | E | 99.4 | 99.4 | 99.3 | 99.5 | 98.1 | 98.2 | 99.5 | 99.6 |
| E_Set1_F2 | E | 97.9 | 97.9 | 99.1 | 99.1 | 97.9 | 97.9 | 99.3 | 99.3 |
| E_Set1_F3 | E | 99.2 | 99.2 | 99.4 | 99.4 | 97.8 | 97.8 | 99.5 | 99.5 |
| E_Set1_LB | E | 98.2 | 99.5 | 99.5 | 99.6 | 98.2 | 98.5 | 99.7 | 99.8 |
| E_Set1_LF | E | 98.5 | 99.0 | 98.8 | 98.8 | 91.0 | 91.1 | 0.737 | 97.8 |
| N_Set1_B1c | N | 98.7 | 98.8 | 99.6 | 99.7 | 96.9 | 97.6 | 99.5 | 99.6 |
| N_Set1_B2 | N | 99.1 | 99.1 | 99.7 | 99.7 | 97.2 | 97.2 | 99.6 | 99.6 |
| N_Set1_B3 | N | 99.1 | 99.1 | 96.1 | 96.1 | 97.5 | 97.5 | 99.7 | 99.7 |
| N_Set1_F1c | N | 99.4 | 99.5 | 99.8 | 99.9 | 98.0 | 98.1 | 98.1 | 98.2 |
| N_Set1_F2 | N | 99.4 | 99.4 | 99.8 | 99.8 | 98.3 | 98.3 | 99.5 | 99.5 |
| N_Set1_F3 | N | 99.4 | 99.4 | 99.7 | 99.7 | 98.1 | 98.1 | 99.5 | 99.5 |
| N_Set1_LB | N | 98.9 | 99.2 | 97.9 | 97.9 | 97.3 | 97.3 | 99.5 | 99.5 |
| N_Set1_LF | N | 99.4 | 99.5 | 99.8 | 99.9 | 98.2 | 98.3 | 99.6 | 99.6 |
| N_Set2_B1c | N | 99.2 | 99.4 | 99.4 | 99.7 | 98.9 | 99.3 | 98.7 | 99.1 |
| N_Set2_B2 | N | 99.5 | 99.5 | 99.9 | 99.9 | 99.3 | 99.3 | 99.7 | 99.7 |
| N_Set2_B3 | N | 98.4 | 98.4 | 99.4 | 99.5 | 97.1 | 97.1 | 99.6 | 99.6 |
| N_Set2_F1c | N | 99.0 | 99.2 | 99.8 | 99.9 | 98.4 | 99.2 | 98.9 | 99.1 |
| N_Set2_F2 | N | 98.4 | 98.4 | 99.0 | 99.0 | 92.6 | 92.6 | 98.7 | 98.7 |
| N_Set2_F3 | N | 99.3 | 99.3 | 98.7 | 98.7 | 98.3 | 98.3 | 98.8 | 98.8 |
| N_Set2_LB | N | 99.5 | 99.5 | 99.9 | 99.9 | 99.1 | 99.3 | 99.1 | 99.2 |
| N_Set2_LF | N | 99.3 | 99.4 | 99.8 | 99.8 | 99.2 | 99.2 | 99.2 | 99.2 |
| RdRp_B1c | RdRp | 99.4 | 99.4 | 99.7 | 99.7 | 98.5 | 98.5 | 1.08 | 1.08 |
| RdRp_B2 | RdRp | 99.2 | 99.2 | 99.7 | 99.7 | 99.0 | 99.0 | 99.6 | 99.7 |
| RdRp_B3 | RdRp | 99.3 | 99.3 | 99.6 | 99.6 | 98.8 | 98.8 | 99.7 | 99.7 |
| RdRp_F1c | RdRp | 99.3 | 99.5 | 99.7 | 99.8 | 99.4 | 99.5 | 99.7 | 99.7 |
| RdRp_F2 | RdRp | 99.4 | 99.4 | 99.9 | 99.9 | 99.4 | 99.4 | 99.7 | 99.7 |
| RdRp_F3 | RdRp | 99.4 | 99.4 | 99.8 | 99.8 | 99.3 | 99.3 | 99.0 | 99.0 |
| RdRp_LB | RdRp | 99.1 | 99.1 | 98.6 | 98.6 | 98.7 | 98.7 | 99.6 | 99.6 |
| RdRp_LF | RdRp | 99.5 | 99.5 | 99.7 | 99.7 | 99.4 | 99.4 | 99.9 | 99.9 |

Table S6: Matched and mismatched coverages for Alves et al. [2] set.

| Primer | Gene Target | Delta | | Lambda | | Mu | | Omicron | |
|---|---|---|---|---|---|---|---|---|---|
| | | $C_{\text{strict}}$ | $C_{\text{part.}}$ | $C_{\text{strict}}$ | $C_{\text{part.}}$ | $C_{\text{strict}}$ | $C_{\text{part.}}$ | $C_{\text{strict}}$ | $C_{\text{part.}}$ |
| E_Set1_B1c | E | 99.7 | 99.7 | 99.2 | 99.2 | 99.6 | 99.6 | 99.2 | 99.2 |
| E_Set1_B2 | E | 99.8 | 99.8 | 99.6 | 99.6 | 99.7 | 99.7 | 0.199 | 0.199 |
| E_Set1_B3 | E | 100. | 100. | 99.9 | 99.9 | 99.9 | 99.9 | 97.5 | 97.5 |
| E_Set1_F1c | E | 99.9 | 99.9 | 99.7 | 99.8 | 99.4 | 99.6 | 99.4 | 99.4 |
| E_Set1_F2 | E | 98.8 | 98.8 | 99.9 | 99.9 | 99.8 | 99.8 | 99.5 | 99.5 |
| E_Set1_F3 | E | 98.0 | 98.0 | 97.4 | 97.4 | 99.4 | 99.4 | 99.5 | 99.5 |
| E_Set1_LB | E | 99.9 | 100. | 99.2 | 100. | 99.5 | 99.9 | 99.4 | 99.4 |
| E_Set1_LF | E | 99.4 | 99.4 | 98.7 | 99.0 | 0.756 | 0.953 | 99.4 | 99.4 |
| N_Set1_B1c | N | 99.9 | 99.9 | 99.3 | 99.4 | 85.5 | 85.7 | 94.2 | 94.3 |
| N_Set1_B2 | N | 99.5 | 99.5 | 97.7 | 97.7 | 99.5 | 99.5 | 99.2 | 99.2 |
| N_Set1_B3 | N | 99.5 | 99.5 | 99.3 | 99.3 | 99.6 | 99.6 | 99.2 | 99.2 |
| N_Set1_F1c | N | 99.9 | 99.9 | 99.6 | 100. | 85.2 | 99.7 | 94.2 | 94.4 |
| N_Set1_F2 | N | 99.9 | 99.9 | 99.9 | 99.9 | 99.5 | 99.5 | 94.6 | 94.6 |
| N_Set1_F3 | N | 99.9 | 99.9 | 99.2 | 99.2 | 99.6 | 99.6 | 94.6 | 94.6 |
| N_Set1_LB | N | 99.4 | 99.5 | 99.4 | 99.5 | 99.6 | 99.6 | 94.8 | 94.8 |
| N_Set1_LF | N | 99.9 | 99.9 | 99.8 | 99.9 | 99.1 | 99.6 | 94.4 | 94.5 |
| N_Set2_B1c | N | 99.8 | 99.9 | 99.2 | 99.9 | 88.7 | 99.7 | 94.7 | 94.8 |
| N_Set2_B2 | N | 100. | 100. | 99.8 | 99.8 | 99.8 | 99.8 | 94.7 | 94.7 |
| N_Set2_B3 | N | 99.5 | 99.6 | 99.6 | 99.7 | 97.2 | 97.2 | 93.8 | 93.8 |
| N_Set2_F1c | N | 99.8 | 99.9 | 99.5 | 99.7 | 99.0 | 99.4 | 0.555 | 0.555 |
| N_Set2_F2 | N | 99.7 | 99.7 | 0.664 | 0.664 | 99.1 | 99.1 | 0.640 | 0.640 |
| N_Set2_F3 | N | 99.8 | 99.8 | 99.3 | 99.3 | 99.5 | 99.5 | 94.6 | 94.6 |
| N_Set2_LB | N | 99.8 | 99.8 | 99.5 | 99.7 | 99.7 | 99.8 | 94.8 | 94.8 |
| N_Set2_LF | N | 98.8 | 98.8 | 98.6 | 98.7 | 99.7 | 99.8 | 94.8 | 94.8 |
| RdRp_B1c | RdRp | 99.8 | 99.8 | 98.5 | 98.5 | 99.8 | 99.8 | 97.9 | 97.9 |
| RdRp_B2 | RdRp | 100. | 100. | 99.8 | 99.8 | 100. | 100. | 99.8 | 99.8 |
| RdRp_B3 | RdRp | 100. | 100. | 98.8 | 98.8 | 99.9 | 99.9 | 99.6 | 99.6 |
| RdRp_F1c | RdRp | 97.4 | 99.9 | 99.1 | 99.8 | 98.9 | 99.9 | 99.7 | 99.9 |
| RdRp_F2 | RdRp | 98.9 | 98.9 | 99.8 | 99.8 | 99.7 | 99.7 | 99.6 | 99.6 |
| RdRp_F3 | RdRp | 100. | 100. | 99.7 | 99.7 | 99.7 | 99.7 | 99.6 | 99.6 |
| RdRp_LB | RdRp | 99.9 | 99.9 | 98.6 | 98.6 | 99.5 | 99.5 | 98.2 | 98.2 |
| RdRp_LF | RdRp | 100. | 100. | 99.7 | 99.7 | 99.9 | 99.9 | 99.6 | 99.6 |

Table S7: Matched and mismatched coverages for Alves et al. [2] set.

| | | BA.2 | | BA.3 | | BA.4 | | BA.5 | |
|---|---|---|---|---|---|---|---|---|---|
| Primer | Gene Target | $C_{\text{strict}}$ | $C_{\text{part.}}$ | $C_{\text{strict}}$ | $C_{\text{part.}}$ | $C_{\text{strict}}$ | $C_{\text{part.}}$ | $C_{\text{strict}}$ | $C_{\text{part.}}$ |
| E_Set1_B1c | E | 99.6 | 99.6 | 99.7 | 99.7 | 99.2 | 99.2 | 99.1 | 99.1 |
| E_Set1_B2 | E | 0.000 | 0.000 | 0.000 | 0.000 | 0.159 | 0.159 | 0.162 | 0.162 |
| E_Set1_B3 | E | 100. | 100. | 98.9 | 98.9 | 100. | 100. | 100. | 100. |
| E_Set1_F1c | E | 99.8 | 99.9 | 100. | 100. | 99.7 | 99.7 | 99.2 | 99.3 |
| E_Set1_F2 | E | 99.8 | 99.8 | 99.1 | 99.1 | 100. | 100. | 99.8 | 99.8 |
| E_Set1_F3 | E | 99.9 | 99.9 | 99.7 | 99.7 | 99.7 | 99.7 | 98.4 | 98.4 |
| E_Set1_LB | E | 99.9 | 100. | 100. | 100. | 99.8 | 99.8 | 99.8 | 100. |
| E_Set1_LF | E | 99.8 | 99.9 | 100. | 100. | 98.9 | 99.2 | 99.6 | 99.7 |
| N_Set1_B1c | N | 99.7 | 99.7 | 96.3 | 96.3 | 100. | 100. | 99.7 | 99.8 |
| N_Set1_B2 | N | 99.9 | 99.9 | 99.4 | 99.4 | 98.9 | 98.9 | 99.4 | 99.4 |
| N_Set1_B3 | N | 99.2 | 99.2 | 99.7 | 99.7 | 0.954 | 0.954 | 97.6 | 97.6 |
| N_Set1_F1c | N | 100. | 100. | 95.7 | 95.7 | 100. | 100. | 99.6 | 99.7 |
| N_Set1_F2 | N | 100. | 100. | 95.7 | 95.7 | 99.7 | 99.7 | 99.6 | 99.6 |
| N_Set1_F3 | N | 98.4 | 98.4 | 95.7 | 95.7 | 99.2 | 99.2 | 98.6 | 98.6 |
| N_Set1_LB | N | 99.6 | 99.6 | 97.7 | 97.7 | 99.8 | 99.8 | 91.8 | 92.0 |
| N_Set1_LF | N | 99.8 | 100. | 95.7 | 95.7 | 99.8 | 100. | 98.2 | 99.5 |
| N_Set2_B1c | N | 99.5 | 99.9 | 96.0 | 96.0 | 99.5 | 99.8 | 98.6 | 98.9 |
| N_Set2_B2 | N | 100. | 100. | 96.0 | 96.0 | 99.7 | 99.7 | 99.1 | 99.1 |
| N_Set2_B3 | N | 99.6 | 99.6 | 94.3 | 94.3 | 99.0 | 99.0 | 98.7 | 98.7 |
| N_Set2_F1c | N | 1.31 | 1.31 | 0.862 | 0.862 | 0.318 | 0.318 | 0.975 | 1.14 |
| N_Set2_F2 | N | 0.000 | 0.000 | 1.15 | 1.15 | 0.477 | 0.477 | 0.569 | 0.569 |
| N_Set2_F3 | N | 99.5 | 99.5 | 94.3 | 94.3 | 100. | 100. | 99.3 | 99.3 |
| N_Set2_LB | N | 99.8 | 99.9 | 95.1 | 96.0 | 99.0 | 99.2 | 97.4 | 99.2 |
| N_Set2_LF | N | 99.9 | 99.9 | 95.4 | 95.4 | 99.0 | 99.0 | 89.4 | 93.6 |
| RdRp_B1c | RdRp | 99.6 | 99.6 | 99.4 | 99.4 | 99.5 | 99.5 | 95.4 | 95.4 |
| RdRp_B2 | RdRp | 99.9 | 99.9 | 99.4 | 99.4 | 99.8 | 99.8 | 99.4 | 99.4 |
| RdRp_B3 | RdRp | 99.9 | 99.9 | 99.7 | 99.7 | 99.7 | 99.7 | 99.4 | 99.4 |
| RdRp_F1c | RdRp | 99.8 | 99.9 | 99.4 | 99.4 | 99.7 | 99.7 | 99.4 | 99.8 |
| RdRp_F2 | RdRp | 100. | 100. | 99.4 | 99.4 | 99.5 | 99.5 | 99.1 | 99.1 |
| RdRp_F3 | RdRp | 100. | 100. | 99.4 | 99.4 | 100. | 100. | 99.6 | 99.6 |
| RdRp_LB | RdRp | 99.8 | 99.8 | 99.4 | 99.4 | 99.8 | 99.8 | 98.1 | 98.1 |
| RdRp_LF | RdRp | 99.9 | 99.9 | 98.6 | 98.6 | 100. | 100. | 100. | 100. |

Table S8: Matched and mismatched coverages for Diego et al. [3] set.

| | | SARS-CoV-2 | | Alpha | | Beta | | Gamma | |
|---|---|---|---|---|---|---|---|---|---|
| Primer | Gene Target | $C_{\text{strict}}$ | $C_{\text{part.}}$ | $C_{\text{strict}}$ | $C_{\text{part.}}$ | $C_{\text{strict}}$ | $C_{\text{part.}}$ | $C_{\text{strict}}$ | $C_{\text{part.}}$ |
| E-B1c | E | 99.3 | 99.3 | 99.5 | 99.5 | 99.0 | 99.0 | 99.7 | 99.7 |
| E-B2 | E | 99.2 | 99.2 | 99.5 | 99.5 | 99.1 | 99.1 | 99.8 | 99.8 |
| E-B3 | E | 99.1 | 99.1 | 99.5 | 99.6 | 0.293 | 0.293 | 98.9 | 98.9 |
| E-F1c | E | 0.000 | 99.2 | 0.000 | 99.6 | 0.000 | 99.3 | 0.000 | 99.8 |
| E-F2 | E | 99.5 | 99.5 | 99.7 | 99.7 | 99.4 | 99.4 | 99.9 | 99.9 |
| E-F3 | E | 98.6 | 98.6 | 99.1 | 99.1 | 98.5 | 98.5 | 99.9 | 99.9 |
| M-B1c | M | 99.3 | 99.4 | 99.8 | 99.9 | 99.0 | 99.0 | 99.8 | 99.9 |
| M-B2 | M | 99.3 | 99.3 | 99.4 | 99.4 | 98.9 | 98.9 | 99.9 | 99.9 |
| M-B3 | M | 99.4 | 99.4 | 99.8 | 99.8 | 99.1 | 99.1 | 99.9 | 99.9 |
| M-F1c | M | 98.6 | 99.3 | 99.6 | 99.8 | 98.4 | 98.7 | 99.7 | 99.7 |
| M-F2 | M | 99.2 | 99.2 | 99.6 | 99.6 | 98.3 | 98.3 | 99.7 | 99.7 |
| M-F3 | M | 99.3 | 99.3 | 98.8 | 98.8 | 98.1 | 98.1 | 99.8 | 99.8 |
| N5-B1c | N | 97.9 | 98.4 | 99.4 | 99.8 | 97.1 | 97.1 | 99.6 | 99.6 |
| N5-B2 | N | 99.4 | 99.4 | 99.7 | 99.7 | 98.1 | 98.1 | 99.5 | 99.5 |
| N5-B3 | N | 99.4 | 99.4 | 99.8 | 99.8 | 98.3 | 98.3 | 99.6 | 99.6 |
| N5-F1c | N | 99.6 | 99.6 | 99.9 | 100. | 99.4 | 99.5 | 99.8 | 99.9 |
| N5-F2 | N | 99.2 | 99.2 | 99.8 | 99.8 | 98.4 | 98.4 | 98.7 | 98.7 |
| N5-F3 | N | 99.0 | 99.1 | 99.8 | 99.8 | 98.4 | 98.4 | 98.9 | 98.9 |
| N5-LB | N | 99.4 | 99.4 | 99.6 | 99.6 | 97.8 | 97.8 | 0.780 | 0.780 |
| N5-LF | N | 99.4 | 99.5 | 99.5 | 99.5 | 99.1 | 99.1 | 99.2 | 99.2 |
| ORF1b-B1c | Orf1b | 98.9 | 99.3 | 99.7 | 99.9 | 99.4 | 99.8 | 99.9 | 100. |
| ORF1b-B2 | Orf1b | 99.3 | 99.3 | 99.8 | 99.8 | 99.8 | 99.8 | 99.8 | 99.8 |
| ORF1b-B3 | Orf1b | 99.5 | 99.5 | 99.8 | 99.8 | 99.8 | 99.8 | 100. | 100. |
| ORF1b-F1c | Orf1b | 98.8 | 98.8 | 98.4 | 98.4 | 99.6 | 99.6 | 99.8 | 99.8 |
| ORF1b-F2 | Orf1b | 99.4 | 99.4 | 99.9 | 99.9 | 99.8 | 99.8 | 99.9 | 99.9 |
| ORF1b-F3 | Orf1b | 98.9 | 98.9 | 99.7 | 99.7 | 99.6 | 99.6 | 99.9 | 99.9 |
| S447-B1c | S | 0.000 | 99.4 | 0.000 | 99.8 | 0.000 | 99.7 | 0.000 | 99.9 |
| S447-B2 | S | 83.8 | 83.8 | 99.9 | 99.9 | 99.6 | 99.6 | 99.9 | 99.9 |
| S447-B3 | S | 99.3 | 99.3 | 99.7 | 99.7 | 99.1 | 99.1 | 99.7 | 99.7 |
| S447-F1c | S | 99.3 | 99.3 | 99.8 | 99.8 | 99.6 | 99.6 | 98.7 | 98.7 |
| S447-F2 | S | 99.4 | 99.4 | 0.911 | 0.911 | 99.6 | 99.6 | 99.9 | 99.9 |
| S447-F3 | S | 99.4 | 99.4 | 99.7 | 99.7 | 99.5 | 99.5 | 99.9 | 99.9 |
| S447-LB | S | 99.4 | 99.4 | 99.8 | 99.8 | 99.7 | 99.7 | 87.7 | 87.7 |
| S447-LF | S | 99.2 | 99.4 | 99.8 | 99.9 | 99.5 | 99.6 | 99.0 | 100. |
| S555-B1c | S | 98.9 | 98.9 | 99.4 | 99.4 | 98.4 | 98.6 | 98.9 | 98.9 |
| S555-B2 | S | 99.5 | 99.5 | 99.4 | 99.4 | 97.7 | 97.7 | 99.0 | 99.0 |

(continued on next page)

(Table S8, Diego et al. [3] coverages, continued)

| Primer | Gene Target | SARS-CoV-2 | | Alpha | | Beta | | Gamma | |
|---|---|---|---|---|---|---|---|---|---|
| | | $C_{\text{strict}}$ | $C_{\text{part.}}$ | $C_{\text{strict}}$ | $C_{\text{part.}}$ | $C_{\text{strict}}$ | $C_{\text{part.}}$ | $C_{\text{strict}}$ | $C_{\text{part.}}$ |
| S555-B3 | S | 99.4 | 99.4 | 99.4 | 99.4 | 98.1 | 98.1 | 98.7 | 98.7 |
| S555-F1c | S | 99.1 | 99.1 | 99.5 | 99.6 | 98.7 | 98.8 | 99.0 | 99.0 |
| S555-F2 | S | 99.3 | 99.3 | 99.6 | 99.6 | 98.2 | 98.2 | 99.1 | 99.1 |
| S555-F3 | S | 99.2 | 99.2 | 99.6 | 99.6 | 98.8 | 98.8 | 99.1 | 99.1 |

Table S9: Matched and mismatched coverages for Diego et al. [3] set.

| | | Delta | | Lambda | | Mu | | Omicron | |
|---|---|---|---|---|---|---|---|---|---|
| Primer | Gene Target | $C_{\text{strict}}$ | $C_{\text{part.}}$ | $C_{\text{strict}}$ | $C_{\text{part.}}$ | $C_{\text{strict}}$ | $C_{\text{part.}}$ | $C_{\text{strict}}$ | $C_{\text{part.}}$ |
| E-B1c | E | 99.9 | 99.9 | 99.2 | 99.2 | 99.4 | 99.4 | 94.4 | 94.4 |
| E-B2 | E | 100. | 100. | 98.6 | 98.6 | 99.5 | 99.5 | 94.2 | 94.2 |
| E-B3 | E | 99.7 | 99.7 | 98.5 | 98.5 | 99.4 | 99.4 | 94.0 | 94.0 |
| E-F1c | E | 0.000 | 99.9 | 0.000 | 99.1 | 0.000 | 99.5 | 0.000 | 97.0 |
| E-F2 | E | 100. | 100. | 99.9 | 99.9 | 99.9 | 99.9 | 97.5 | 97.5 |
| E-F3 | E | 99.9 | 99.9 | 99.7 | 99.7 | 99.7 | 99.7 | 99.6 | 99.6 |
| M-B1c | M | 99.9 | 100. | 99.3 | 99.6 | 99.9 | 99.9 | 0.569 | 0.569 |
| M-B2 | M | 0.0126 | 0.0126 | 97.2 | 97.2 | 99.9 | 99.9 | 96.1 | 96.1 |
| M-B3 | M | 99.9 | 99.9 | 99.7 | 99.7 | 99.7 | 99.7 | 97.9 | 97.9 |
| M-F1c | M | 99.8 | 99.9 | 98.4 | 98.5 | 99.2 | 99.2 | 96.7 | 96.7 |
| M-F2 | M | 99.9 | 99.9 | 98.9 | 98.9 | 99.8 | 99.8 | 95.9 | 95.9 |
| M-F3 | M | 99.8 | 99.8 | 97.5 | 97.5 | 96.7 | 96.7 | 97.7 | 97.7 |
| N5-B1c | N | 99.5 | 99.8 | 99.6 | 99.7 | 97.1 | 99.3 | 93.8 | 93.8 |
| N5-B2 | N | 99.9 | 99.9 | 99.2 | 99.2 | 99.5 | 99.5 | 94.5 | 94.5 |
| N5-B3 | N | 99.9 | 99.9 | 99.9 | 99.9 | 99.1 | 99.1 | 94.6 | 94.6 |
| N5-F1c | N | 99.8 | 99.9 | 94.8 | 95.0 | 99.5 | 99.8 | 94.9 | 94.9 |
| N5-F2 | N | 99.8 | 99.8 | 99.2 | 99.2 | 88.8 | 88.8 | 94.9 | 94.9 |
| N5-F3 | N | 99.8 | 99.9 | 99.5 | 99.6 | 99.1 | 99.2 | 0.555 | 0.555 |
| N5-LB | N | 99.9 | 99.9 | 99.5 | 99.5 | 97.0 | 97.1 | 93.8 | 93.8 |
| N5-LF | N | 99.7 | 99.8 | 99.6 | 99.6 | 99.7 | 99.7 | 94.8 | 94.8 |
| ORF1b-B1c | Orf1b | 99.7 | 99.9 | 99.6 | 99.9 | 99.7 | 100. | 98.1 | 99.8 |
| ORF1b-B2 | Orf1b | 99.9 | 99.9 | 99.6 | 99.6 | 99.9 | 99.9 | 99.8 | 99.8 |
| ORF1b-B3 | Orf1b | 99.9 | 99.9 | 99.8 | 99.8 | 99.8 | 99.8 | 99.7 | 99.7 |
| ORF1b-F1c | Orf1b | 99.5 | 99.6 | 99.1 | 99.1 | 99.4 | 99.7 | 99.6 | 99.6 |
| ORF1b-F2 | Orf1b | 100. | 100. | 99.9 | 99.9 | 99.8 | 99.8 | 99.9 | 99.9 |
| ORF1b-F3 | Orf1b | 99.9 | 99.9 | 99.5 | 99.5 | 99.4 | 99.4 | 99.9 | 99.9 |
| S447-B1c | S | 0.000 | 100. | 0.000 | 99.7 | 0.000 | 99.9 | 0.000 | 99.8 |
| S447-B2 | S | 99.8 | 99.8 | 99.8 | 99.8 | 99.8 | 99.8 | 99.8 | 99.8 |
| S447-B3 | S | 99.8 | 99.8 | 99.6 | 99.6 | 99.7 | 99.7 | 99.8 | 99.8 |
| S447-F1c | S | 99.7 | 99.7 | 99.7 | 99.7 | 98.7 | 98.7 | 99.8 | 99.8 |
| S447-F2 | S | 100. | 100. | 99.8 | 99.8 | 99.9 | 99.9 | 99.8 | 99.8 |
| S447-F3 | S | 99.4 | 99.4 | 99.6 | 99.6 | 99.8 | 99.8 | 99.7 | 99.7 |
| S447-LB | S | 100. | 100. | 99.6 | 99.8 | 100. | 100. | 99.8 | 99.8 |
| S447-LF | S | 99.8 | 99.9 | 96.7 | 99.4 | 97.1 | 99.4 | 99.8 | 99.8 |
| S555-B1c | S | 99.9 | 99.9 | 99.3 | 99.3 | 98.7 | 98.8 | 98.4 | 98.4 |
| S555-B2 | S | 100. | 100. | 99.3 | 99.3 | 99.0 | 99.0 | 0.341 | 0.341 |

(continued on next page)

(Table S9, Diego et al. [3] set coverages, continued)

| | | Delta | | Lambda | | Mu | | Omicron | |
|---|---|---|---|---|---|---|---|---|---|
| Primer | Gene Target | $C_{\text{strict}}$ | $C_{\text{part.}}$ | $C_{\text{strict}}$ | $C_{\text{part.}}$ | $C_{\text{strict}}$ | $C_{\text{part.}}$ | $C_{\text{strict}}$ | $C_{\text{part.}}$ |
| S555-B3 | S | 99.9 | 99.9 | 99.8 | 99.8 | 99.6 | 99.6 | 98.9 | 98.9 |
| S555-F1c | S | 99.9 | 99.9 | 99.3 | 99.7 | 99.6 | 99.7 | 98.5 | 98.5 |
| S555-F2 | S | 100. | 100. | 99.8 | 99.8 | 99.6 | 99.6 | 98.5 | 98.5 |
| S555-F3 | S | 100. | 100. | 99.7 | 99.7 | 99.8 | 99.8 | 98.6 | 98.6 |

Table S10: Matched and mismatched coverages for Diego et al. [3] set.

| | | BA.2 | | BA.3 | | BA.4 | | BA.5 | |
|---|---|---|---|---|---|---|---|---|---|
| Primer | Gene Target | $C_{\text{strict}}$ | $C_{\text{part.}}$ | $C_{\text{strict}}$ | $C_{\text{part.}}$ | $C_{\text{strict}}$ | $C_{\text{part.}}$ | $C_{\text{strict}}$ | $C_{\text{part.}}$ |
| E-B1c | E | 100. | 100. | 97.7 | 97.7 | 98.6 | 98.6 | 98.1 | 98.1 |
| E-B2 | E | 100. | 100. | 98.0 | 98.0 | 99.2 | 99.2 | 99.8 | 99.8 |
| E-B3 | E | 99.8 | 99.8 | 97.7 | 97.7 | 99.4 | 99.4 | 99.4 | 99.4 |
| E-F1c | E | 0.000 | 99.9 | 0.000 | 97.7 | 0.000 | 100. | 0.000 | 99.5 |
| E-F2 | E | 100. | 100. | 98.9 | 98.9 | 100. | 100. | 100. | 100. |
| E-F3 | E | 99.9 | 99.9 | 100. | 100. | 99.7 | 99.7 | 99.6 | 99.6 |
| M-B1c | M | 0.000 | 0.000 | 0.287 | 0.287 | 0.000 | 0.000 | 0.000 | 0.000 |
| M-B2 | M | 100. | 100. | 99.1 | 99.1 | 99.8 | 99.8 | 99.5 | 99.5 |
| M-B3 | M | 100. | 100. | 98.9 | 98.9 | 99.7 | 99.7 | 99.8 | 99.8 |
| M-F1c | M | 99.5 | 99.5 | 99.7 | 99.7 | 94.9 | 99.5 | 97.1 | 98.8 |
| M-F2 | M | 99.8 | 99.8 | 93.4 | 93.4 | 98.9 | 98.9 | 99.2 | 99.2 |
| M-F3 | M | 99.8 | 99.8 | 93.7 | 93.7 | 98.1 | 98.1 | 99.4 | 99.4 |
| N5-B1c | N | 99.6 | 99.8 | 94.3 | 94.3 | 99.0 | 99.0 | 98.7 | 98.7 |
| N5-B2 | N | 98.4 | 98.4 | 95.7 | 95.7 | 99.2 | 99.2 | 98.6 | 98.6 |
| N5-B3 | N | 100. | 100. | 95.7 | 95.7 | 99.7 | 99.7 | 98.5 | 98.5 |
| N5-F1c | N | 98.1 | 98.1 | 89.4 | 89.9 | 99.2 | 99.4 | 96.7 | 98.5 |
| N5-F2 | N | 99.6 | 99.6 | 96.0 | 96.0 | 99.7 | 99.7 | 98.9 | 98.9 |
| N5-F3 | N | 1.31 | 1.31 | 0.862 | 0.862 | 0.318 | 0.318 | 0.975 | 0.975 |
| N5-LB | N | 99.8 | 99.8 | 94.3 | 94.3 | 98.7 | 99.0 | 98.6 | 98.9 |
| N5-LF | N | 99.9 | 99.9 | 96.0 | 96.0 | 99.5 | 99.5 | 99.3 | 99.4 |
| ORF1b-B1c | Orf1b | 99.9 | 100. | 100. | 100. | 99.4 | 100. | 99.9 | 100. |
| ORF1b-B2 | Orf1b | 100. | 100. | 100. | 100. | 100. | 100. | 99.9 | 99.9 |
| ORF1b-B3 | Orf1b | 100. | 100. | 100. | 100. | 99.7 | 99.7 | 99.7 | 99.7 |
| ORF1b-F1c | Orf1b | 99.6 | 99.6 | 100. | 100. | 100. | 100. | 99.8 | 99.9 |
| ORF1b-F2 | Orf1b | 100. | 100. | 100. | 100. | 100. | 100. | 99.6 | 99.6 |
| ORF1b-F3 | Orf1b | 99.7 | 99.7 | 100. | 100. | 98.9 | 98.9 | 98.8 | 98.8 |
| S447-B1c | S | 0.000 | 100. | 0.000 | 99.7 | 0.000 | 100. | 0.000 | 99.8 |
| S447-B2 | S | 100. | 100. | 99.7 | 99.7 | 100. | 100. | 99.8 | 99.8 |
| S447-B3 | S | 99.9 | 99.9 | 99.7 | 99.7 | 100. | 100. | 100. | 100. |
| S447-F1c | S | 100. | 100. | 99.7 | 99.7 | 99.7 | 99.7 | 99.2 | 99.3 |
| S447-F2 | S | 100. | 100. | 99.7 | 99.7 | 100. | 100. | 99.7 | 99.7 |
| S447-F3 | S | 99.9 | 99.9 | 99.4 | 99.4 | 99.7 | 99.7 | 99.8 | 99.8 |
| S447-LB | S | 99.9 | 99.9 | 99.7 | 99.7 | 100. | 100. | 99.8 | 99.8 |
| S447-LF | S | 99.8 | 99.9 | 99.7 | 99.7 | 99.8 | 99.8 | 99.7 | 99.8 |
| S555-B1c | S | 99.8 | 99.8 | 97.4 | 97.4 | 98.6 | 98.6 | 99.4 | 99.5 |
| S555-B2 | S | 0.000 | 0.000 | 0.000 | 0.000 | 0.000 | 0.000 | 0.244 | 0.244 |

(continued on next page)

(Table S10, Diego et al. [3] set coverages, continued)

| | | BA.2 | | BA.3 | | BA.4 | | BA.5 | |
|---|---|---|---|---|---|---|---|---|---|
| Primer | Gene Target | $C_{\text{strict}}$ | $C_{\text{part.}}$ | $C_{\text{strict}}$ | $C_{\text{part.}}$ | $C_{\text{strict}}$ | $C_{\text{part.}}$ | $C_{\text{strict}}$ | $C_{\text{part.}}$ |
| S555-B3 | S | 100. | 100. | 97.7 | 97.7 | 100. | 100. | 99.8 | 99.8 |
| S555-F1c | S | 99.6 | 99.8 | 97.4 | 97.4 | 99.8 | 99.8 | 99.4 | 99.8 |
| S555-F2 | S | 100. | 100. | 97.1 | 97.1 | 100. | 100. | 99.9 | 99.9 |
| S555-F3 | S | 99.9 | 99.9 | 97.4 | 97.4 | 100. | 100. | 99.6 | 99.6 |

Table S11: Matched and mismatched coverages for Ganguli et al. [4] set.

| | | SARS-CoV-2 | | Alpha | | Beta | | Gamma | |
|---|---|---|---|---|---|---|---|---|---|
| Primer | Gene Target | $C_{\text{strict}}$ | $C_{\text{part.}}$ | $C_{\text{strict}}$ | $C_{\text{part.}}$ | $C_{\text{strict}}$ | $C_{\text{part.}}$ | $C_{\text{strict}}$ | $C_{\text{part.}}$ |
| N-P1-B1c | N | 98.6 | 98.7 | 99.4 | 99.4 | 96.7 | 96.7 | 99.8 | 99.8 |
| N-P1-B2 | N | 97.4 | 97.4 | 0.124 | 0.124 | 96.2 | 96.2 | 99.7 | 99.7 |
| N-P1-B3 | N | 99.5 | 99.5 | 99.3 | 99.3 | 96.3 | 96.3 | 99.6 | 99.6 |
| N-P1-F1c | N | 97.9 | 98.5 | 98.6 | 98.7 | 96.6 | 96.6 | 99.4 | 99.4 |
| N-P1-F2 | N | 0.000 | 0.000 | 0.000 | 0.000 | 0.000 | 0.000 | 0.000 | 0.000 |
| N-P1-F3 | N | 98.3 | 98.3 | 99.6 | 99.6 | 96.9 | 96.9 | 99.8 | 99.8 |
| N-P1-Loop-B | N | 98.9 | 99.1 | 99.3 | 99.4 | 96.5 | 96.5 | 99.5 | 99.5 |
| N-P2-B1c | N | 98.6 | 99.4 | 99.6 | 99.8 | 96.7 | 97.9 | 99.4 | 99.5 |
| N-P2-B2 | N | 99.0 | 99.0 | 98.1 | 98.1 | 97.4 | 97.4 | 99.6 | 99.6 |
| N-P2-B3 | N | 99.2 | 99.2 | 99.0 | 99.0 | 97.8 | 97.8 | 99.7 | 99.7 |
| N-P2-F1c | N | 99.4 | 99.4 | 99.8 | 99.8 | 98.0 | 98.0 | 98.1 | 98.1 |
| N-P2-F2 | N | 99.4 | 99.5 | 99.8 | 99.8 | 98.2 | 98.2 | 99.6 | 99.6 |
| N-P2-F3 | N | 99.4 | 99.4 | 99.8 | 99.8 | 98.3 | 98.3 | 99.5 | 99.5 |
| N-P2-Loop-B | N | 99.4 | 99.4 | 99.4 | 99.8 | 97.5 | 97.7 | 99.4 | 99.6 |
| N-P3-B1c | N | 99.1 | 99.3 | 98.3 | 98.4 | 95.6 | 95.7 | 97.2 | 97.4 |
| N-P3-B2 | N | 99.3 | 99.3 | 98.5 | 98.5 | 95.7 | 95.7 | 97.4 | 97.4 |
| N-P3-B3 | N | 99.4 | 99.4 | 98.2 | 98.2 | 96.2 | 96.2 | 97.6 | 97.6 |
| N-P3-F1c | N | 98.3 | 98.6 | 98.4 | 98.7 | 95.5 | 95.7 | 96.3 | 97.0 |
| N-P3-F2 | N | 99.0 | 99.0 | 99.3 | 99.3 | 89.4 | 89.4 | 99.3 | 99.3 |
| N-P3-F3 | N | 99.3 | 99.3 | 99.7 | 99.7 | 99.2 | 99.2 | 99.8 | 99.8 |
| N-P3-Loop-B | N | 99.0 | 99.4 | 98.4 | 98.6 | 95.4 | 95.8 | 97.2 | 97.4 |
| Orf1a-P1-B1c | Orf1a | 98.9 | 98.9 | 99.4 | 99.5 | 97.9 | 97.9 | 98.6 | 98.6 |
| Orf1a-P1-B2 | Orf1a | 99.3 | 99.3 | 99.5 | 99.5 | 90.9 | 90.9 | 98.9 | 98.9 |
| Orf1a-P1-B3 | Orf1a | 99.2 | 99.2 | 98.9 | 98.9 | 99.0 | 99.0 | 100. | 100. |
| Orf1a-P1-F1c | Orf1a | 99.0 | 99.1 | 99.3 | 99.3 | 98.0 | 98.0 | 98.7 | 98.7 |
| Orf1a-P1-F2 | Orf1a | 99.0 | 99.0 | 99.6 | 99.6 | 97.9 | 97.9 | 98.7 | 98.7 |
| Orf1a-P1-F3 | Orf1a | 99.1 | 99.1 | 99.6 | 99.6 | 98.0 | 98.0 | 98.7 | 98.7 |
| Orf1a-P1-Loop-F | Orf1a | 98.9 | 99.1 | 89.2 | 89.9 | 97.9 | 97.9 | 98.3 | 98.4 |
| Orf1a-P2-B1c | Orf1a | 99.1 | 99.2 | 99.2 | 99.4 | 98.1 | 98.5 | 99.6 | 99.6 |
| Orf1a-P2-B2 | Orf1a | 99.3 | 99.3 | 99.1 | 99.1 | 98.7 | 98.7 | 99.7 | 99.7 |
| Orf1a-P2-B3 | Orf1a | 99.3 | 99.3 | 99.4 | 99.4 | 98.6 | 98.6 | 99.7 | 99.7 |
| Orf1a-P2-F1c | Orf1a | 99.1 | 99.2 | 98.9 | 99.7 | 98.8 | 98.9 | 100. | 100. |
| Orf1a-P2-F2 | Orf1a | 99.2 | 99.2 | 99.5 | 99.5 | 96.3 | 96.3 | 98.6 | 98.6 |
| Orf1a-P2-F3 | Orf1a | 98.9 | 98.9 | 99.5 | 99.5 | 97.9 | 97.9 | 98.6 | 98.6 |
| Orf1a-P3-B1c | Orf1a | 99.3 | 99.3 | 99.4 | 99.5 | 98.6 | 98.7 | 99.7 | 99.7 |
| Orf1a-P3-B2 | Orf1a | 96.0 | 96.0 | 99.5 | 99.5 | 98.8 | 98.8 | 99.8 | 99.8 |

(continued on next page)

(Table S11, Ganguli et al. [4] coverages, continued)

| Primer | Gene Target | SARS-CoV-2 | | Alpha | | Beta | | Gamma | |
|---|---|---|---|---|---|---|---|---|---|
| | | $C_{\text{strict}}$ | $C_{\text{part.}}$ | $C_{\text{strict}}$ | $C_{\text{part.}}$ | $C_{\text{strict}}$ | $C_{\text{part.}}$ | $C_{\text{strict}}$ | $C_{\text{part.}}$ |
| Orf1a-P3-B3 | Orf1a | 99.1 | 99.1 | 99.4 | 99.4 | 97.9 | 97.9 | 99.7 | 99.7 |
| Orf1a-P3-F1c | Orf1a | 99.3 | 99.3 | 99.2 | 99.4 | 98.7 | 98.7 | 99.7 | 99.7 |
| Orf1a-P3-F2 | Orf1a | 99.1 | 99.1 | 98.9 | 98.9 | 98.5 | 98.5 | 99.8 | 99.8 |
| Orf1a-P3-F3 | Orf1a | 99.3 | 99.3 | 99.4 | 99.4 | 98.2 | 98.2 | 99.6 | 99.6 |
| Orf8-P1-B1c | Orf8 | 98.9 | 99.1 | 98.4 | 99.3 | 97.4 | 98.0 | 90.3 | 90.6 |
| Orf8-P1-B2 | Orf8 | 99.1 | 99.1 | 0.179 | 0.179 | 98.1 | 98.1 | 89.7 | 89.7 |
| Orf8-P1-B3 | Orf8 | 1.25 | 1.25 | 0.0690 | 0.0690 | 0.0267 | 0.0267 | 0.000 | 0.000 |
| Orf8-P1-F1c | Orf8 | 93.7 | 93.7 | 0.0552 | 0.0552 | 98.0 | 98.1 | 89.8 | 89.9 |
| Orf8-P1-F2 | Orf8 | 98.8 | 98.8 | 99.2 | 99.2 | 98.3 | 98.3 | 89.9 | 89.9 |
| Orf8-P1-F3 | Orf8 | 99.2 | 99.2 | 99.2 | 99.2 | 98.5 | 98.5 | 93.3 | 93.3 |
| Orf8-P1-Loop-F | Orf8 | 99.0 | 99.3 | 98.4 | 99.3 | 98.3 | 98.5 | 90.1 | 90.3 |
| Orf8-P2-B1c | Orf8 | 0.000 | 0.000 | 0.000 | 0.000 | 0.000 | 0.000 | 0.000 | 0.000 |
| Orf8-P2-B2 | Orf8 | 0.000 | 0.000 | 0.000 | 0.000 | 0.000 | 0.000 | 0.000 | 0.000 |
| Orf8-P2-B3 | Orf8 | 99.3 | 99.3 | 99.8 | 99.8 | 99.1 | 99.1 | 2.21 | 2.21 |
| Orf8-P2-F1c | Orf8 | 99.1 | 99.3 | 0.193 | 98.9 | 98.1 | 98.5 | 89.6 | 90.3 |
| Orf8-P2-F2 | Orf8 | 99.0 | 99.0 | 98.4 | 98.4 | 97.9 | 97.9 | 90.6 | 90.6 |
| Orf8-P2-F3 | Orf8 | 99.2 | 99.2 | 0.0552 | 0.0552 | 97.7 | 97.7 | 89.9 | 89.9 |
| Orf8-P2-Loop-B | Orf8 | 99.2 | 99.3 | 0.0966 | 99.8 | 98.1 | 99.0 | 93.5 | 94.8 |
| Orf8-P3-B1c | Orf8 | 99.1 | 99.2 | 0.193 | 98.9 | 98.1 | 98.4 | 89.7 | 90.3 |
| Orf8-P3-B2 | Orf8 | 98.0 | 98.0 | 68.5 | 68.5 | 97.8 | 97.8 | 90.1 | 90.1 |
| Orf8-P3-B3 | Orf8 | 99.3 | 99.3 | 0.0966 | 0.0966 | 98.3 | 98.3 | 93.6 | 93.6 |
| Orf8-P3-F1c | Orf8 | 99.0 | 99.0 | 98.4 | 98.4 | 97.9 | 97.9 | 90.5 | 90.5 |
| Orf8-P3-F2 | Orf8 | 93.4 | 93.4 | 0.0552 | 0.0552 | 98.1 | 98.1 | 89.9 | 89.9 |
| Orf8-P3-F3 | Orf8 | 99.1 | 99.1 | 98.5 | 98.5 | 98.3 | 98.3 | 90.1 | 90.1 |
| S-P1-B1c | S | 99.1 | 99.2 | 99.0 | 99.0 | 98.2 | 98.3 | 99.4 | 99.4 |
| S-P1-B2 | S | 98.6 | 98.6 | 99.5 | 99.5 | 0.333 | 0.333 | 99.4 | 99.4 |
| S-P1-B3 | S | 99.2 | 99.2 | 98.9 | 98.9 | 99.2 | 99.2 | 98.7 | 98.7 |
| S-P1-F1c | S | 98.4 | 98.5 | 97.8 | 98.0 | 98.9 | 99.1 | 99.1 | 99.2 |
| S-P1-F2 | S | 99.4 | 99.4 | 98.6 | 98.6 | 98.8 | 98.8 | 99.6 | 99.6 |
| S-P1-F3 | S | 98.5 | 98.5 | 98.5 | 98.5 | 97.7 | 97.7 | 99.6 | 99.6 |
| S-P1-Loop-B | S | 98.7 | 98.9 | 0.207 | 0.207 | 98.3 | 99.3 | 99.2 | 99.2 |
| S-P2-B1c | S | 98.7 | 99.2 | 99.2 | 99.5 | 98.8 | 99.4 | 99.3 | 99.5 |
| S-P2-B2 | S | 98.9 | 98.9 | 97.0 | 97.0 | 98.9 | 98.9 | 98.8 | 98.8 |
| S-P2-B3 | S | 99.3 | 99.3 | 99.7 | 99.7 | 99.3 | 99.3 | 99.6 | 99.6 |
| S-P2-F1c | S | 99.1 | 99.2 | 98.9 | 99.0 | 98.9 | 99.0 | 99.3 | 99.4 |
| S-P2-F2 | S | 99.1 | 99.1 | 98.6 | 98.6 | 99.3 | 99.3 | 99.8 | 99.8 |
| S-P2-F3 | S | 99.3 | 99.3 | 98.4 | 98.4 | 98.1 | 98.1 | 99.5 | 99.5 |
| S-P2-Loop-B | S | 99.0 | 99.3 | 99.2 | 99.2 | 99.2 | 99.2 | 98.0 | 98.0 |
| S-P3-B1c | S | 98.7 | 99.2 | 99.2 | 99.5 | 98.8 | 99.4 | 99.3 | 99.5 |
| S-P3-B2 | S | 98.9 | 98.9 | 97.0 | 97.0 | 98.9 | 98.9 | 98.8 | 98.8 |
| S-P3-B3 | S | 99.3 | 99.3 | 99.7 | 99.7 | 99.3 | 99.3 | 99.6 | 99.6 |

(continued on next page)

(Table S11, Ganguli et al. [4] coverages, continued)

| Primer | Gene Target | SARS-CoV-2 | | Alpha | | Beta | | Gamma | |
|---|---|---|---|---|---|---|---|---|---|
| | | $C_{\text{strict}}$ | $C_{\text{part.}}$ | $C_{\text{strict}}$ | $C_{\text{part.}}$ | $C_{\text{strict}}$ | $C_{\text{part.}}$ | $C_{\text{strict}}$ | $C_{\text{part.}}$ |
| S-P3-F1c | S | 99.1 | 99.2 | 98.9 | 99.1 | 98.2 | 99.3 | 99.4 | 99.5 |
| S-P3-F2 | S | 99.0 | 99.0 | 98.4 | 98.4 | 99.2 | 99.2 | 99.7 | 99.7 |
| S-P3-F3 | S | 99.4 | 99.4 | 98.6 | 98.6 | 98.8 | 98.8 | 99.6 | 99.6 |
| S-P3-Loop-B | S | 99.0 | 99.3 | 99.2 | 99.2 | 99.2 | 99.2 | 98.0 | 98.0 |

Table S12: Matched and mismatched coverages for Ganguli et al. [4] set.

| Primer | Gene Target | Delta | | Lambda | | Mu | | Omicron | |
|---|---|---|---|---|---|---|---|---|---|
| | | $C_{\text{strict}}$ | $C_{\text{part.}}$ | $C_{\text{strict}}$ | $C_{\text{part.}}$ | $C_{\text{strict}}$ | $C_{\text{part.}}$ | $C_{\text{strict}}$ | $C_{\text{part.}}$ |
| N-P1-B1c | N | 99.8 | 99.8 | 99.5 | 99.6 | 99.8 | 99.8 | 99.1 | 99.1 |
| N-P1-B2 | N | 99.5 | 99.5 | 99.6 | 99.6 | 99.0 | 99.0 | 99.0 | 99.0 |
| N-P1-B3 | N | 99.7 | 99.7 | 99.4 | 99.4 | 98.6 | 98.6 | 99.0 | 99.0 |
| N-P1-F1c | N | 99.6 | 99.7 | 99.3 | 99.5 | 99.4 | 99.5 | 99.2 | 99.2 |
| N-P1-F2 | N | 0.000 | 0.000 | 0.000 | 0.000 | 0.000 | 0.000 | 0.000 | 0.000 |
| N-P1-F3 | N | 99.8 | 99.8 | 99.5 | 99.5 | 99.5 | 99.5 | 99.2 | 99.2 |
| N-P1-Loop-B | N | 99.9 | 99.9 | 99.4 | 99.5 | 99.6 | 99.7 | 99.0 | 99.0 |
| N-P2-B1c | N | 99.9 | 99.9 | 99.3 | 99.7 | 85.4 | 99.8 | 94.2 | 94.2 |
| N-P2-B2 | N | 99.6 | 99.6 | 99.6 | 99.6 | 99.6 | 99.6 | 95.2 | 95.2 |
| N-P2-B3 | N | 99.6 | 99.6 | 99.4 | 99.4 | 99.7 | 99.7 | 99.2 | 99.2 |
| N-P2-F1c | N | 99.9 | 99.9 | 99.6 | 99.7 | 85.2 | 85.2 | 94.2 | 94.2 |
| N-P2-F2 | N | 99.9 | 99.9 | 99.8 | 99.8 | 99.1 | 99.1 | 94.4 | 94.5 |
| N-P2-F3 | N | 99.9 | 99.9 | 99.9 | 99.9 | 99.5 | 99.5 | 94.6 | 94.6 |
| N-P2-Loop-B | N | 99.8 | 100. | 99.3 | 99.6 | 99.7 | 99.8 | 94.2 | 94.2 |
| N-P3-B1c | N | 92.8 | 99.7 | 80.1 | 80.7 | 98.4 | 98.9 | 99.0 | 99.0 |
| N-P3-B2 | N | 99.9 | 99.9 | 81.9 | 81.9 | 98.8 | 98.8 | 99.1 | 99.1 |
| N-P3-B3 | N | 98.2 | 98.2 | 87.6 | 87.6 | 97.9 | 97.9 | 95.4 | 95.4 |
| N-P3-F1c | N | 0.0881 | 0.0881 | 79.0 | 79.4 | 94.0 | 94.5 | 98.8 | 98.8 |
| N-P3-F2 | N | 99.2 | 99.2 | 93.6 | 93.6 | 99.1 | 99.1 | 99.3 | 99.3 |
| N-P3-F3 | N | 99.9 | 99.9 | 99.1 | 99.1 | 99.8 | 99.8 | 99.7 | 99.7 |
| N-P3-Loop-B | N | 99.8 | 100. | 80.3 | 81.1 | 98.4 | 99.0 | 99.0 | 99.0 |
| Orf1a-P1-B1c | Orf1a | 99.8 | 99.9 | 99.4 | 99.4 | 97.3 | 97.3 | 99.7 | 99.7 |
| Orf1a-P1-B2 | Orf1a | 99.7 | 99.8 | 99.5 | 99.5 | 99.7 | 99.7 | 99.6 | 99.6 |
| Orf1a-P1-B3 | Orf1a | 99.9 | 99.9 | 99.7 | 99.7 | 99.6 | 99.6 | 99.0 | 99.0 |
| Orf1a-P1-F1c | Orf1a | 100. | 100. | 99.6 | 99.8 | 99.7 | 99.7 | 99.7 | 99.7 |
| Orf1a-P1-F2 | Orf1a | 99.7 | 99.7 | 99.7 | 99.7 | 98.9 | 98.9 | 99.7 | 99.7 |
| Orf1a-P1-F3 | Orf1a | 99.9 | 99.9 | 99.6 | 99.6 | 97.4 | 97.4 | 99.7 | 99.7 |
| Orf1a-P1-Loop-F | Orf1a | 99.4 | 99.8 | 98.3 | 98.5 | 98.9 | 99.8 | 99.6 | 99.6 |
| Orf1a-P2-B1c | Orf1a | 99.9 | 99.9 | 99.1 | 99.4 | 99.6 | 99.6 | 98.4 | 98.4 |
| Orf1a-P2-B2 | Orf1a | 99.2 | 99.2 | 99.5 | 99.5 | 98.5 | 98.5 | 98.6 | 98.6 |
| Orf1a-P2-B3 | Orf1a | 100. | 100. | 99.8 | 99.8 | 99.8 | 99.8 | 98.1 | 98.1 |
| Orf1a-P2-F1c | Orf1a | 99.8 | 100. | 99.7 | 99.8 | 99.6 | 99.7 | 99.0 | 99.2 |
| Orf1a-P2-F2 | Orf1a | 99.9 | 99.9 | 99.5 | 99.5 | 99.6 | 99.6 | 99.6 | 99.7 |
| Orf1a-P2-F3 | Orf1a | 99.9 | 99.9 | 99.4 | 99.4 | 97.3 | 97.3 | 99.7 | 99.7 |
| Orf1a-P3-B1c | Orf1a | 99.9 | 100. | 99.8 | 99.9 | 99.8 | 99.9 | 98.1 | 98.5 |
| Orf1a-P3-B2 | Orf1a | 98.7 | 98.7 | 99.7 | 99.7 | 99.9 | 99.9 | 98.6 | 98.6 |

(continued on next page)

(Table S12, Ganguli et al. [4] set coverages, continued)

| Primer | Gene Target | Delta | | Lambda | | Mu | | Omicron | |
|---|---|---|---|---|---|---|---|---|---|
| | | $C_{strict}$ | $C_{part.}$ | $C_{strict}$ | $C_{part.}$ | $C_{strict}$ | $C_{part.}$ | $C_{strict}$ | $C_{part.}$ |
| Orf1a-P3-B3 | Orf1a | 99.9 | 99.9 | 99.5 | 99.5 | 97.7 | 97.7 | 98.7 | 98.7 |
| Orf1a-P3-F1c | Orf1a | 99.2 | 99.2 | 98.9 | 99.1 | 99.7 | 99.7 | 98.6 | 98.6 |
| Orf1a-P3-F2 | Orf1a | 99.9 | 99.9 | 99.2 | 99.2 | 99.8 | 99.8 | 98.4 | 98.4 |
| Orf1a-P3-F3 | Orf1a | 99.9 | 99.9 | 99.5 | 99.5 | 99.7 | 99.7 | 98.4 | 98.4 |
| Orf8-P1-B1c | Orf8 | 99.5 | 99.8 | 96.0 | 97.1 | 0.0908 | 0.0908 | 84.2 | 84.3 |
| Orf8-P1-B2 | Orf8 | 99.7 | 99.7 | 93.9 | 93.9 | 96.5 | 96.5 | 84.2 | 84.2 |
| Orf8-P1-B3 | Orf8 | 0.000 | 0.000 | 0.000 | 0.000 | 0.000 | 0.000 | 0.000 | 0.000 |
| Orf8-P1-F1c | Orf8 | 99.7 | 99.8 | 98.9 | 99.2 | 99.0 | 99.1 | 83.5 | 83.5 |
| Orf8-P1-F2 | Orf8 | 98.9 | 98.9 | 97.9 | 98.0 | 0.333 | 0.333 | 83.5 | 83.5 |
| Orf8-P1-F3 | Orf8 | 99.6 | 99.6 | 99.1 | 99.1 | 99.7 | 99.7 | 82.3 | 82.3 |
| Orf8-P1-Loop-F | Orf8 | 99.6 | 99.9 | 97.9 | 99.6 | 99.3 | 99.8 | 83.4 | 83.5 |
| Orf8-P2-B1c | Orf8 | 0.000 | 0.000 | 0.000 | 0.000 | 0.000 | 0.000 | 0.000 | 0.000 |
| Orf8-P2-B2 | Orf8 | 0.000 | 0.000 | 0.000 | 0.000 | 0.000 | 0.000 | 0.000 | 0.000 |
| Orf8-P2-B3 | Orf8 | 99.9 | 99.9 | 99.0 | 99.0 | 99.2 | 99.2 | 91.9 | 91.9 |
| Orf8-P2-F1c | Orf8 | 99.7 | 99.8 | 92.9 | 99.5 | 96.5 | 99.8 | 84.1 | 84.2 |
| Orf8-P2-F2 | Orf8 | 99.5 | 99.5 | 96.1 | 96.1 | 0.0908 | 0.0908 | 84.3 | 84.3 |
| Orf8-P2-F3 | Orf8 | 99.7 | 99.7 | 99.3 | 99.3 | 99.6 | 99.6 | 84.1 | 84.1 |
| Orf8-P2-Loop-B | Orf8 | 99.4 | 99.8 | 99.0 | 99.5 | 99.4 | 99.8 | 85.8 | 85.8 |
| Orf8-P3-B1c | Orf8 | 99.7 | 99.8 | 92.9 | 98.6 | 96.5 | 99.3 | 84.1 | 84.2 |
| Orf8-P3-B2 | Orf8 | 98.9 | 98.9 | 97.3 | 97.3 | 3.77 | 3.77 | 84.0 | 84.0 |
| Orf8-P3-B3 | Orf8 | 99.4 | 99.4 | 99.2 | 99.2 | 99.5 | 99.5 | 85.8 | 85.8 |
| Orf8-P3-F1c | Orf8 | 99.5 | 99.5 | 96.1 | 96.1 | 0.0908 | 0.0908 | 84.3 | 84.3 |
| Orf8-P3-F2 | Orf8 | 99.8 | 99.8 | 98.9 | 98.9 | 99.0 | 99.0 | 83.6 | 83.6 |
| Orf8-P3-F3 | Orf8 | 99.6 | 99.6 | 98.1 | 98.1 | 99.4 | 99.4 | 83.5 | 83.5 |
| S-P1-B1c | S | 90.4 | 90.4 | 94.3 | 94.7 | 99.1 | 99.2 | 0.498 | 0.498 |
| S-P1-B2 | S | 84.2 | 84.2 | 99.0 | 99.0 | 99.0 | 99.0 | 98.2 | 98.2 |
| S-P1-B3 | S | 38.0 | 38.0 | 99.4 | 99.4 | 0.862 | 0.862 | 0.341 | 0.341 |
| S-P1-F1c | S | 91.4 | 91.6 | 94.2 | 95.0 | 99.0 | 99.3 | 98.3 | 98.4 |
| S-P1-F2 | S | 99.9 | 99.9 | 97.5 | 97.5 | 99.3 | 99.3 | 97.6 | 97.6 |
| S-P1-F3 | S | 99.9 | 99.9 | 98.9 | 98.9 | 99.1 | 99.1 | 97.3 | 97.3 |
| S-P1-Loop-B | S | 88.7 | 88.7 | 0.385 | 91.3 | 98.6 | 98.9 | 0.441 | 0.455 |
| S-P2-B1c | S | 85.1 | 87.0 | 0.396 | 0.418 | 98.8 | 99.3 | 98.2 | 98.3 |
| S-P2-B2 | S | 38.0 | 38.0 | 99.0 | 99.0 | 0.877 | 0.877 | 0.341 | 0.341 |
| S-P2-B3 | S | 82.6 | 82.6 | 99.6 | 99.6 | 99.4 | 99.4 | 98.3 | 98.3 |
| S-P2-F1c | S | 90.5 | 90.5 | 94.4 | 94.9 | 99.2 | 99.4 | 96.7 | 96.7 |
| S-P2-F2 | S | 99.8 | 99.8 | 97.7 | 97.8 | 99.5 | 99.5 | 98.4 | 98.4 |
| S-P2-F3 | S | 99.9 | 99.9 | 98.3 | 98.3 | 99.3 | 99.3 | 97.5 | 97.5 |
| S-P2-Loop-B | S | 84.8 | 84.8 | 98.8 | 98.9 | 98.9 | 99.0 | 97.6 | 97.6 |
| S-P3-B1c | S | 85.1 | 87.0 | 0.396 | 0.418 | 98.8 | 99.3 | 98.2 | 98.3 |
| S-P3-B2 | S | 38.0 | 38.0 | 99.0 | 99.0 | 0.877 | 0.877 | 0.341 | 0.341 |
| S-P3-B3 | S | 82.6 | 82.6 | 99.6 | 99.6 | 99.4 | 99.4 | 98.3 | 98.3 |

(continued on next page)

(Table S12, Ganguli et al. [4] set coverages, continued)

| | | Delta | | Lambda | | Mu | | Omicron | |
|---|---|---|---|---|---|---|---|---|---|
| Primer | Gene Target | $C_{\text{strict}}$ | $C_{\text{part.}}$ | $C_{\text{strict}}$ | $C_{\text{part.}}$ | $C_{\text{strict}}$ | $C_{\text{part.}}$ | $C_{\text{strict}}$ | $C_{\text{part.}}$ |
| S-P3-F1c | S | 90.4 | 90.5 | 94.3 | 95.1 | 99.1 | 99.4 | 0.498 | 96.5 |
| S-P3-F2 | S | 92.6 | 92.6 | 99.0 | 99.0 | 99.3 | 99.3 | 98.5 | 98.5 |
| S-P3-F3 | S | 99.9 | 99.9 | 98.3 | 98.3 | 99.2 | 99.2 | 97.6 | 97.6 |
| S-P3-Loop-B | S | 84.8 | 84.8 | 98.8 | 98.9 | 98.9 | 99.0 | 97.6 | 97.6 |

Table S13: Matched and mismatched coverages for Ganguli et al. [4] set.

| | | BA.2 | | BA.3 | | BA.4 | | BA.5 | |
|---|---|---|---|---|---|---|---|---|---|
| Primer | Gene Target | $C_{\text{strict}}$ | $C_{\text{part.}}$ | $C_{\text{strict}}$ | $C_{\text{part.}}$ | $C_{\text{strict}}$ | $C_{\text{part.}}$ | $C_{\text{strict}}$ | $C_{\text{part.}}$ |
| N-P1-B1c | N | 99.9 | 99.9 | 100. | 100. | 100. | 100. | 99.5 | 99.5 |
| N-P1-B2 | N | 99.6 | 99.6 | 99.7 | 99.7 | 99.2 | 99.2 | 99.7 | 99.7 |
| N-P1-B3 | N | 99.4 | 99.4 | 99.7 | 99.7 | 100. | 100. | 99.6 | 99.6 |
| N-P1-F1c | N | 99.6 | 99.7 | 100. | 100. | 100. | 100. | 99.8 | 99.9 |
| N-P1-F2 | N | 0.000 | 0.000 | 0.000 | 0.000 | 0.000 | 0.000 | 0.000 | 0.000 |
| N-P1-F3 | N | 100. | 100. | 100. | 100. | 100. | 100. | 99.4 | 99.4 |
| N-P1-Loop-B | N | 99.6 | 99.9 | 99.1 | 99.1 | 99.2 | 99.2 | 99.7 | 99.8 |
| N-P2-B1c | N | 99.8 | 99.9 | 96.3 | 96.3 | 100. | 100. | 99.3 | 99.4 |
| N-P2-B2 | N | 99.7 | 99.7 | 97.7 | 97.7 | 100. | 100. | 99.3 | 99.3 |
| N-P2-B3 | N | 99.2 | 99.2 | 99.7 | 99.7 | 0.954 | 0.954 | 97.5 | 97.5 |
| N-P2-F1c | N | 100. | 100. | 95.7 | 95.7 | 100. | 100. | 99.6 | 99.6 |
| N-P2-F2 | N | 99.8 | 99.9 | 95.7 | 95.7 | 99.8 | 99.8 | 98.2 | 98.2 |
| N-P2-F3 | N | 100. | 100. | 95.7 | 95.7 | 99.7 | 99.7 | 99.6 | 99.6 |
| N-P2-Loop-B | N | 99.7 | 99.8 | 96.3 | 96.3 | 99.8 | 100. | 92.3 | 96.1 |
| N-P3-B1c | N | 99.9 | 99.9 | 98.0 | 98.0 | 99.5 | 99.7 | 99.3 | 99.3 |
| N-P3-B2 | N | 99.9 | 99.9 | 97.7 | 97.7 | 99.7 | 99.7 | 99.4 | 99.4 |
| N-P3-B3 | N | 0.0676 | 0.0676 | 1.15 | 1.15 | 0.000 | 0.000 | 0.0812 | 0.0812 |
| N-P3-F1c | N | 99.9 | 99.9 | 96.8 | 97.1 | 99.0 | 99.7 | 98.2 | 98.8 |
| N-P3-F2 | N | 99.5 | 99.5 | 98.9 | 98.9 | 99.2 | 99.2 | 99.4 | 99.4 |
| N-P3-F3 | N | 99.9 | 99.9 | 99.7 | 99.7 | 100. | 100. | 99.7 | 99.7 |
| N-P3-Loop-B | N | 99.8 | 99.9 | 97.4 | 97.4 | 99.8 | 99.8 | 98.7 | 99.3 |
| Orf1a-P1-B1c | Orf1a | 99.8 | 99.9 | 99.1 | 99.1 | 98.1 | 98.1 | 99.5 | 99.5 |
| Orf1a-P1-B2 | Orf1a | 99.9 | 99.9 | 98.6 | 98.6 | 100. | 100. | 98.1 | 98.1 |
| Orf1a-P1-B3 | Orf1a | 99.9 | 99.9 | 98.6 | 98.6 | 99.8 | 99.8 | 96.1 | 96.1 |
| Orf1a-P1-F1c | Orf1a | 99.9 | 99.9 | 99.1 | 99.1 | 98.9 | 98.9 | 99.6 | 99.8 |
| Orf1a-P1-F2 | Orf1a | 100. | 100. | 99.7 | 99.7 | 100. | 100. | 99.7 | 99.7 |
| Orf1a-P1-F3 | Orf1a | 99.8 | 99.8 | 99.7 | 99.7 | 98.7 | 98.7 | 97.0 | 97.0 |
| Orf1a-P1-Loop-F | Orf1a | 99.8 | 99.8 | 99.7 | 99.7 | 99.7 | 99.7 | 99.7 | 99.8 |
| Orf1a-P2-B1c | Orf1a | 100. | 100. | 98.9 | 98.9 | 99.5 | 99.5 | 99.7 | 99.8 |
| Orf1a-P2-B2 | Orf1a | 100. | 100. | 98.6 | 98.6 | 99.4 | 99.4 | 99.5 | 99.5 |
| Orf1a-P2-B3 | Orf1a | 100. | 100. | 99.1 | 99.1 | 99.7 | 99.7 | 99.8 | 99.8 |
| Orf1a-P2-F1c | Orf1a | 99.9 | 100. | 98.6 | 98.6 | 99.8 | 100. | 96.0 | 96.3 |
| Orf1a-P2-F2 | Orf1a | 99.7 | 99.7 | 98.9 | 98.9 | 99.7 | 99.7 | 99.1 | 99.1 |
| Orf1a-P2-F3 | Orf1a | 99.8 | 99.8 | 99.1 | 99.1 | 98.1 | 98.1 | 99.5 | 99.5 |
| Orf1a-P3-B1c | Orf1a | 99.9 | 100. | 99.1 | 99.1 | 99.7 | 99.7 | 99.8 | 99.8 |
| Orf1a-P3-B2 | Orf1a | 100. | 100. | 99.1 | 99.1 | 100. | 100. | 99.5 | 99.5 |

(continued on next page)

(Table S13, Ganguli et al. [4] set coverages, continued)

| Primer | Gene Target | BA.2 | | BA.3 | | BA.4 | | BA.5 | |
|---|---|---|---|---|---|---|---|---|---|
| | | $C_{strict}$ | $C_{part.}$ | $C_{strict}$ | $C_{part.}$ | $C_{strict}$ | $C_{part.}$ | $C_{strict}$ | $C_{part.}$ |
| Orf1a-P3-B3 | Orf1a | 99.8 | 99.8 | 99.1 | 99.1 | 99.8 | 99.8 | 99.6 | 99.6 |
| Orf1a-P3-F1c | Orf1a | 100. | 100. | 98.6 | 98.6 | 99.5 | 99.5 | 99.7 | 99.7 |
| Orf1a-P3-F2 | Orf1a | 100. | 100. | 98.9 | 98.9 | 99.8 | 99.8 | 99.7 | 99.7 |
| Orf1a-P3-F3 | Orf1a | 99.7 | 99.7 | 98.6 | 98.6 | 99.2 | 99.2 | 99.5 | 99.5 |
| Orf8-P1-B1c | Orf8 | 99.7 | 99.8 | 82.8 | 88.5 | 98.3 | 98.6 | 98.7 | 99.0 |
| Orf8-P1-B2 | Orf8 | 99.8 | 99.8 | 88.5 | 88.5 | 98.7 | 98.7 | 99.4 | 99.4 |
| Orf8-P1-B3 | Orf8 | 0.0135 | 0.0135 | 0.000 | 0.000 | 0.000 | 0.000 | 0.000 | 0.000 |
| Orf8-P1-F1c | Orf8 | 99.8 | 99.8 | 88.8 | 88.8 | 98.7 | 98.7 | 98.5 | 99.4 |
| Orf8-P1-F2 | Orf8 | 97.8 | 97.8 | 88.2 | 88.2 | 97.9 | 98.3 | 99.1 | 99.1 |
| Orf8-P1-F3 | Orf8 | 99.7 | 99.7 | 87.6 | 87.6 | 97.5 | 97.5 | 15.6 | 15.6 |
| Orf8-P1-Loop-F | Orf8 | 99.6 | 99.9 | 88.2 | 88.8 | 98.4 | 98.6 | 98.3 | 99.2 |
| Orf8-P2-B1c | Orf8 | 0.000 | 0.000 | 0.000 | 0.000 | 0.000 | 0.000 | 0.000 | 0.000 |
| Orf8-P2-B2 | Orf8 | 0.000 | 0.000 | 0.000 | 0.000 | 0.000 | 0.000 | 0.000 | 0.000 |
| Orf8-P2-B3 | Orf8 | 99.7 | 99.7 | 95.4 | 95.4 | 100. | 100. | 96.5 | 96.5 |
| Orf8-P2-F1c | Orf8 | 99.8 | 99.9 | 88.5 | 88.5 | 98.7 | 98.9 | 99.4 | 99.6 |
| Orf8-P2-F2 | Orf8 | 99.8 | 99.8 | 88.8 | 88.8 | 98.7 | 98.7 | 99.0 | 99.0 |
| Orf8-P2-F3 | Orf8 | 99.7 | 99.7 | 83.0 | 83.0 | 98.6 | 98.6 | 98.6 | 98.6 |
| Orf8-P2-Loop-B | Orf8 | 99.8 | 99.8 | 93.7 | 93.7 | 98.6 | 99.0 | 96.8 | 96.8 |
| Orf8-P3-B1c | Orf8 | 99.8 | 99.8 | 88.5 | 88.5 | 98.7 | 98.7 | 99.4 | 99.6 |
| Orf8-P3-B2 | Orf8 | 99.3 | 99.3 | 89.1 | 89.1 | 98.3 | 98.3 | 96.3 | 96.3 |
| Orf8-P3-B3 | Orf8 | 99.8 | 99.8 | 94.0 | 94.0 | 98.7 | 98.7 | 96.8 | 96.8 |
| Orf8-P3-F1c | Orf8 | 99.8 | 99.8 | 88.5 | 88.5 | 98.7 | 98.7 | 99.0 | 99.0 |
| Orf8-P3-F2 | Orf8 | 99.8 | 99.8 | 88.8 | 88.8 | 98.7 | 98.7 | 98.5 | 98.5 |
| Orf8-P3-F3 | Orf8 | 99.6 | 99.6 | 88.2 | 88.2 | 98.4 | 98.4 | 98.3 | 98.3 |
| S-P1-B1c | S | 92.0 | 92.0 | 0.862 | 0.862 | 98.1 | 98.1 | 94.4 | 94.6 |
| S-P1-B2 | S | 92.4 | 92.4 | 97.4 | 97.4 | 99.2 | 99.2 | 98.5 | 98.5 |
| S-P1-B3 | S | 99.9 | 99.9 | 0.862 | 0.862 | 98.9 | 98.9 | 98.9 | 98.9 |
| S-P1-F1c | S | 91.7 | 91.8 | 100. | 100. | 95.7 | 97.3 | 93.9 | 94.2 |
| S-P1-F2 | S | 90.6 | 90.6 | 99.1 | 99.1 | 97.5 | 97.5 | 94.1 | 94.1 |
| S-P1-F3 | S | 99.9 | 99.9 | 99.4 | 99.4 | 99.8 | 99.8 | 99.3 | 99.3 |
| S-P1-Loop-B | S | 92.1 | 92.2 | 0.287 | 0.287 | 3.02 | 3.18 | 4.55 | 4.87 |
| S-P2-B1c | S | 92.3 | 92.4 | 100. | 100. | 99.2 | 99.7 | 97.6 | 97.9 |
| S-P2-B2 | S | 99.9 | 99.9 | 0.862 | 0.862 | 98.9 | 98.9 | 98.8 | 98.8 |
| S-P2-B3 | S | 99.9 | 99.9 | 98.0 | 98.0 | 99.8 | 99.8 | 99.8 | 99.8 |
| S-P2-F1c | S | 91.9 | 92.0 | 100. | 100. | 97.8 | 97.9 | 93.7 | 93.9 |
| S-P2-F2 | S | 90.0 | 90.0 | 100. | 100. | 96.7 | 96.7 | 92.4 | 92.4 |
| S-P2-F3 | S | 90.6 | 90.6 | 98.9 | 98.9 | 97.6 | 97.6 | 94.4 | 94.4 |
| S-P2-Loop-B | S | 92.9 | 92.9 | 97.1 | 97.1 | 99.8 | 99.8 | 98.9 | 98.9 |
| S-P3-B1c | S | 92.3 | 92.4 | 100. | 100. | 99.2 | 99.7 | 97.6 | 97.9 |
| S-P3-B2 | S | 99.9 | 99.9 | 0.862 | 0.862 | 98.9 | 98.9 | 98.8 | 98.8 |
| S-P3-B3 | S | 99.9 | 99.9 | 98.0 | 98.0 | 99.8 | 99.8 | 99.8 | 99.8 |

(continued on next page)

(Table S13, Ganguli et al. [4] set coverages, continued)

| Primer | Gene Target | BA.2 | | BA.3 | | BA.4 | | BA.5 | |
|---|---|---|---|---|---|---|---|---|---|
| | | $C_{\text{strict}}$ | $C_{\text{part.}}$ | $C_{\text{strict}}$ | $C_{\text{part.}}$ | $C_{\text{strict}}$ | $C_{\text{part.}}$ | $C_{\text{strict}}$ | $C_{\text{part.}}$ |
| S-P3-F1c | S | 92.0 | 92.0 | 0.862 | 100. | 98.1 | 98.3 | 94.4 | 95.1 |
| S-P3-F2 | S | 90.0 | 90.0 | 100. | 100. | 96.8 | 96.8 | 92.5 | 92.5 |
| S-P3-F3 | S | 90.5 | 90.5 | 99.4 | 99.4 | 97.3 | 97.3 | 93.7 | 93.7 |
| S-P3-Loop-B | S | 92.9 | 92.9 | 97.1 | 97.1 | 99.8 | 99.8 | 98.9 | 98.9 |

Table S14: Matched and mismatched coverages for Garcia-Venzor et al. [5] set.

| Primer | Gene Target | SARS-CoV-2 | | Alpha | | Beta | | Gamma | |
|---|---|---|---|---|---|---|---|---|---|
| | | $C_{\text{strict}}$ | $C_{\text{part.}}$ | $C_{\text{strict}}$ | $C_{\text{part.}}$ | $C_{\text{strict}}$ | $C_{\text{part.}}$ | $C_{\text{strict}}$ | $C_{\text{part.}}$ |
| N-geneB1c | N | 99.3 | 99.3 | 99.7 | 99.7 | 97.7 | 97.7 | 99.8 | 99.8 |
| N-geneB2 | N | 99.2 | 99.2 | 98.0 | 98.0 | 97.5 | 97.5 | 99.5 | 99.5 |
| N-geneB3 | N | 99.2 | 99.2 | 99.7 | 99.7 | 97.7 | 97.7 | 99.1 | 99.1 |
| N-geneF1c | N | 99.0 | 99.1 | 99.4 | 99.7 | 98.0 | 98.3 | 99.0 | 99.8 |
| N-geneF2 | N | 99.3 | 99.3 | 99.6 | 99.6 | 98.6 | 98.6 | 99.7 | 99.7 |
| N-geneF3 | N | 99.2 | 99.2 | 99.5 | 99.5 | 98.3 | 98.3 | 99.9 | 99.9 |
| N-geneLB | N | 98.8 | 98.8 | 96.8 | 96.8 | 97.3 | 97.3 | 99.4 | 99.4 |
| N-geneLF | N | 99.0 | 99.0 | 99.6 | 99.6 | 98.4 | 98.4 | 99.6 | 99.6 |

Table S15: Matched and mismatched coverages for Garcia-Venzor et al. [5] set.

| | | Delta | | Lambda | | Mu | | Omicron | |
|---|---|---|---|---|---|---|---|---|---|
| Primer | Gene Target | $C_{\text{strict}}$ | $C_{\text{part.}}$ | $C_{\text{strict}}$ | $C_{\text{part.}}$ | $C_{\text{strict}}$ | $C_{\text{part.}}$ | $C_{\text{strict}}$ | $C_{\text{part.}}$ |
| N-geneB1c | N | 99.9 | 99.9 | 98.9 | 98.9 | 99.8 | 99.8 | 99.3 | 99.3 |
| N-geneB2 | N | 99.7 | 99.7 | 98.8 | 98.8 | 99.2 | 99.2 | 99.5 | 99.5 |
| N-geneB3 | N | 99.7 | 99.7 | 1.04 | 1.04 | 98.9 | 98.9 | 86.5 | 86.5 |
| N-geneF1c | N | 99.7 | 99.9 | 98.6 | 99.5 | 99.4 | 99.9 | 99.3 | 99.3 |
| N-geneF2 | N | 100. | 100. | 99.5 | 99.5 | 99.8 | 99.8 | 99.6 | 99.6 |
| N-geneF3 | N | 99.7 | 99.7 | 99.2 | 99.2 | 99.3 | 99.3 | 99.5 | 99.5 |
| N-geneLB | N | 98.8 | 98.8 | 98.8 | 98.8 | 99.0 | 99.0 | 99.5 | 99.5 |
| N-geneLF | N | 99.9 | 99.9 | 99.3 | 99.3 | 99.7 | 99.7 | 99.4 | 99.4 |

Table S16: Matched and mismatched coverages for Garcia-Venzor et al. [5] set.

| | | BA.2 | | BA.3 | | BA.4 | | BA.5 | |
|---|---|---|---|---|---|---|---|---|---|
| Primer | Gene Target | $C_{\text{strict}}$ | $C_{\text{part.}}$ | $C_{\text{strict}}$ | $C_{\text{part.}}$ | $C_{\text{strict}}$ | $C_{\text{part.}}$ | $C_{\text{strict}}$ | $C_{\text{part.}}$ |
| N-geneB1c | N | 99.9 | 99.9 | 99.7 | 99.7 | 99.4 | 99.4 | 98.1 | 98.1 |
| N-geneB2 | N | 99.9 | 99.9 | 99.4 | 99.4 | 97.6 | 97.6 | 99.3 | 99.3 |
| N-geneB3 | N | 99.2 | 99.2 | 18.7 | 18.7 | 99.8 | 99.8 | 99.4 | 99.4 |
| N-geneF1c | N | 99.7 | 99.8 | 100. | 100. | 99.5 | 99.7 | 99.5 | 99.8 |
| N-geneF2 | N | 99.9 | 99.9 | 99.7 | 99.7 | 99.8 | 99.8 | 100. | 100. |
| N-geneF3 | N | 99.7 | 99.7 | 100. | 100. | 99.8 | 99.8 | 99.6 | 99.6 |
| N-geneLB | N | 99.7 | 99.7 | 99.7 | 99.7 | 99.7 | 99.7 | 99.4 | 99.4 |
| N-geneLF | N | 100. | 100. | 100. | 100. | 99.5 | 99.5 | 99.8 | 99.8 |

Table S17: Matched and mismatched coverages for Huang et al. [6] set.

| Primer | Gene Target | SARS-CoV-2 | | Alpha | | Beta | | Gamma | |
|---|---|---|---|---|---|---|---|---|---|
| | | $C_{\text{strict}}$ | $C_{\text{part.}}$ | $C_{\text{strict}}$ | $C_{\text{part.}}$ | $C_{\text{strict}}$ | $C_{\text{part.}}$ | $C_{\text{strict}}$ | $C_{\text{part.}}$ |
| N1-B1c | N | 99.2 | 99.4 | 99.4 | 99.7 | 98.9 | 99.3 | 98.7 | 99.1 |
| N1-B2 | N | 99.5 | 99.5 | 99.9 | 99.9 | 99.3 | 99.3 | 99.7 | 99.7 |
| N1-B3 | N | 98.4 | 98.4 | 99.4 | 99.5 | 97.1 | 97.1 | 99.6 | 99.6 |
| N1-F1c | N | 99.0 | 99.2 | 99.8 | 99.9 | 98.4 | 99.2 | 98.9 | 99.1 |
| N1-F2 | N | 98.4 | 98.4 | 99.0 | 99.0 | 92.6 | 92.6 | 98.7 | 98.7 |
| N1-F3 | N | 99.3 | 99.3 | 98.7 | 98.7 | 98.3 | 98.3 | 98.8 | 98.8 |
| N1-LB | N | 99.5 | 99.5 | 99.9 | 99.9 | 99.1 | 99.3 | 99.1 | 99.2 |
| N1-LF | N | 99.3 | 99.4 | 99.8 | 99.8 | 99.2 | 99.2 | 99.2 | 99.2 |
| N15-B1c | N | 0.000 | 99.1 | 0.000 | 99.6 | 0.000 | 96.6 | 0.000 | 98.6 |
| N15-B2 | N | 97.3 | 97.3 | 98.0 | 98.0 | 96.7 | 96.7 | 0.563 | 0.563 |
| N15-B3 | N | 98.1 | 98.1 | 98.6 | 98.6 | 96.6 | 96.7 | 99.4 | 99.4 |
| N15-F1c | N | 99.1 | 99.2 | 99.4 | 99.6 | 96.6 | 96.7 | 98.9 | 99.9 |
| N15-F2 | N | 99.1 | 99.1 | 96.9 | 97.2 | 97.6 | 97.6 | 99.7 | 99.7 |
| N15-F3 | N | 99.1 | 99.1 | 99.7 | 99.7 | 97.2 | 97.2 | 99.6 | 99.6 |
| N15-LB | N | 98.0 | 99.0 | 99.5 | 99.6 | 96.7 | 96.8 | 97.6 | 99.9 |
| N15-LF | N | 98.8 | 99.1 | 99.4 | 99.6 | 96.4 | 96.7 | 99.5 | 99.8 |
| O117-B1c | Orf1ab | 98.9 | 98.9 | 99.2 | 99.3 | 98.2 | 98.3 | 99.3 | 99.3 |
| O117-B2 | Orf1ab | 99.2 | 99.2 | 99.3 | 99.3 | 98.4 | 98.4 | 99.4 | 99.4 |
| O117-B3 | Orf1ab | 99.3 | 99.3 | 99.2 | 99.2 | 98.5 | 98.5 | 99.0 | 99.0 |
| O117-F1c | Orf1ab | 98.8 | 99.1 | 99.1 | 99.4 | 98.2 | 98.6 | 98.4 | 99.5 |
| O117-F2 | Orf1ab | 97.9 | 97.9 | 99.2 | 99.2 | 98.1 | 98.1 | 99.3 | 99.3 |
| O117-F3 | Orf1ab | 99.2 | 99.2 | 99.3 | 99.3 | 98.5 | 98.5 | 99.2 | 99.2 |
| O117-LB | Orf1ab | 99.3 | 99.3 | 98.6 | 98.6 | 98.5 | 98.5 | 99.3 | 99.3 |
| O117-LF | Orf1ab | 99.1 | 99.1 | 99.1 | 99.1 | 98.3 | 98.4 | 99.0 | 99.0 |
| S17-B1c | S | 98.7 | 99.2 | 99.2 | 99.5 | 98.8 | 99.4 | 99.3 | 99.5 |
| S17-B2 | S | 98.9 | 98.9 | 97.0 | 97.0 | 98.9 | 98.9 | 98.8 | 98.8 |
| S17-B3 | S | 99.3 | 99.3 | 99.7 | 99.7 | 99.3 | 99.3 | 99.6 | 99.6 |
| S17-F1c | S | 99.2 | 99.2 | 99.0 | 99.0 | 98.9 | 99.0 | 99.4 | 99.4 |
| S17-F2 | S | 99.4 | 99.4 | 98.6 | 98.6 | 98.8 | 98.8 | 99.6 | 99.6 |
| S17-F3 | S | 98.5 | 98.5 | 98.5 | 98.5 | 97.7 | 97.7 | 99.6 | 99.6 |
| S17-LB | S | 99.0 | 99.3 | 99.2 | 99.2 | 99.2 | 99.2 | 98.1 | 98.1 |
| S17-LF | S | 98.4 | 98.5 | 97.8 | 98.0 | 98.9 | 99.1 | 99.1 | 99.2 |

Table S18: Matched and mismatched coverages for Huang et al. [6] set.

| | | Delta | | Lambda | | Mu | | Omicron | |
|---|---|---|---|---|---|---|---|---|---|
| Primer | Gene Target | $C_{\text{strict}}$ | $C_{\text{part.}}$ | $C_{\text{strict}}$ | $C_{\text{part.}}$ | $C_{\text{strict}}$ | $C_{\text{part.}}$ | $C_{\text{strict}}$ | $C_{\text{part.}}$ |
| N1-B1c | N | 99.8 | 99.9 | 99.2 | 99.9 | 88.7 | 99.7 | 94.7 | 94.8 |
| N1-B2 | N | 100. | 100. | 99.8 | 99.8 | 99.8 | 99.8 | 94.7 | 94.7 |
| N1-B3 | N | 99.5 | 99.6 | 99.6 | 99.7 | 97.2 | 97.2 | 93.8 | 93.8 |
| N1-F1c | N | 99.8 | 99.9 | 99.5 | 99.7 | 99.0 | 99.4 | 0.555 | 0.555 |
| N1-F2 | N | 99.7 | 99.7 | 0.664 | 0.664 | 99.1 | 99.1 | 0.640 | 0.640 |
| N1-F3 | N | 99.8 | 99.8 | 99.3 | 99.3 | 99.5 | 99.5 | 94.6 | 94.6 |
| N1-LB | N | 99.8 | 99.8 | 99.5 | 99.7 | 99.7 | 99.8 | 94.8 | 94.8 |
| N1-LF | N | 98.8 | 98.8 | 98.6 | 98.6 | 99.7 | 99.8 | 94.8 | 94.8 |
| N15-B1c | N | 0.000 | 100. | 0.0107 | 99.8 | 0.000 | 99.8 | 0.000 | 99.2 |
| N15-B2 | N | 99.9 | 99.9 | 96.2 | 96.2 | 99.4 | 99.4 | 99.0 | 99.0 |
| N15-B3 | N | 99.5 | 99.6 | 0.107 | 0.107 | 99.4 | 99.4 | 99.2 | 99.2 |
| N15-F1c | N | 99.9 | 100. | 98.5 | 99.9 | 99.6 | 99.8 | 99.3 | 99.3 |
| N15-F2 | N | 99.8 | 99.8 | 98.9 | 99.0 | 99.6 | 99.6 | 99.3 | 99.3 |
| N15-F3 | N | 99.5 | 99.5 | 97.7 | 97.7 | 99.5 | 99.5 | 99.2 | 99.2 |
| N15-LB | N | 99.8 | 100. | 71.1 | 99.5 | 99.3 | 99.8 | 99.1 | 99.2 |
| N15-LF | N | 99.7 | 99.9 | 99.2 | 99.5 | 99.8 | 99.9 | 99.2 | 99.3 |
| O117-B1c | Orf1ab | 99.3 | 99.4 | 99.6 | 99.7 | 99.0 | 99.6 | 99.4 | 99.4 |
| O117-B2 | Orf1ab | 99.9 | 99.9 | 99.6 | 99.6 | 99.7 | 99.7 | 99.5 | 99.5 |
| O117-B3 | Orf1ab | 99.9 | 99.9 | 99.6 | 99.6 | 99.8 | 99.8 | 99.4 | 99.4 |
| O117-F1c | Orf1ab | 99.9 | 100. | 99.5 | 99.9 | 99.5 | 99.9 | 99.4 | 99.4 |
| O117-F2 | Orf1ab | 99.9 | 99.9 | 99.5 | 99.5 | 99.1 | 99.1 | 98.0 | 98.0 |
| O117-F3 | Orf1ab | 99.9 | 99.9 | 99.6 | 99.6 | 99.2 | 99.2 | 99.5 | 99.5 |
| O117-LB | Orf1ab | 99.8 | 99.8 | 99.7 | 99.7 | 99.8 | 99.8 | 99.5 | 99.5 |
| O117-LF | Orf1ab | 99.8 | 99.8 | 99.4 | 99.5 | 99.3 | 99.5 | 99.3 | 99.3 |
| S17-B1c | S | 85.1 | 87.0 | 0.396 | 0.418 | 98.8 | 99.3 | 98.2 | 98.3 |
| S17-B2 | S | 38.0 | 38.0 | 99.0 | 99.0 | 0.877 | 0.877 | 0.341 | 0.341 |
| S17-B3 | S | 82.6 | 82.6 | 99.6 | 99.6 | 99.4 | 99.4 | 98.3 | 98.3 |
| S17-F1c | S | 90.5 | 90.5 | 94.5 | 94.9 | 99.3 | 99.4 | 96.7 | 96.7 |
| S17-F2 | S | 99.9 | 99.9 | 98.3 | 98.3 | 99.2 | 99.2 | 97.6 | 97.6 |
| S17-F3 | S | 99.9 | 99.9 | 98.9 | 98.9 | 99.1 | 99.1 | 97.3 | 97.3 |
| S17-LB | S | 84.7 | 84.7 | 98.8 | 99.1 | 99.0 | 99.0 | 97.6 | 97.6 |
| S17-LF | S | 91.4 | 91.6 | 94.2 | 95.0 | 99.0 | 99.3 | 98.3 | 98.4 |

Table S19: Matched and mismatched coverages for Huang et al. [6] set.

| Primer | Gene Target | BA.2 | | BA.3 | | BA.4 | | BA.5 | |
|---|---|---|---|---|---|---|---|---|---|
| | | $C_{\text{strict}}$ | $C_{\text{part.}}$ | $C_{\text{strict}}$ | $C_{\text{part.}}$ | $C_{\text{strict}}$ | $C_{\text{part.}}$ | $C_{\text{strict}}$ | $C_{\text{part.}}$ |
| N1-B1c | N | 99.5 | 99.9 | 96.0 | 96.0 | 99.5 | 99.8 | 98.6 | 98.9 |
| N1-B2 | N | 100. | 100. | 96.0 | 96.0 | 99.7 | 99.7 | 99.1 | 99.1 |
| N1-B3 | N | 99.6 | 99.6 | 94.3 | 94.3 | 99.0 | 99.0 | 98.7 | 98.7 |
| N1-F1c | N | 1.31 | 1.31 | 0.862 | 0.862 | 0.318 | 0.318 | 0.975 | 1.14 |
| N1-F2 | N | 0.000 | 0.000 | 1.15 | 1.15 | 0.477 | 0.477 | 0.569 | 0.569 |
| N1-F3 | N | 99.5 | 99.5 | 94.3 | 94.3 | 100. | 100. | 99.3 | 99.3 |
| N1-LB | N | 99.8 | 99.9 | 95.1 | 96.0 | 99.0 | 99.2 | 97.4 | 99.2 |
| N1-LF | N | 99.9 | 99.9 | 95.4 | 95.4 | 99.0 | 99.0 | 89.4 | 89.4 |
| N15-B1c | N | 0.000 | 99.9 | 0.000 | 99.7 | 0.000 | 98.9 | 0.000 | 98.9 |
| N15-B2 | N | 99.4 | 99.4 | 99.7 | 99.7 | 96.0 | 96.0 | 99.1 | 99.1 |
| N15-B3 | N | 99.6 | 99.7 | 100. | 100. | 100. | 100. | 99.9 | 99.9 |
| N15-F1c | N | 100. | 100. | 99.7 | 99.7 | 99.7 | 100. | 99.7 | 99.8 |
| N15-F2 | N | 99.9 | 99.9 | 99.7 | 99.7 | 99.7 | 99.7 | 99.5 | 99.6 |
| N15-F3 | N | 99.9 | 99.9 | 99.4 | 99.4 | 98.9 | 98.9 | 99.4 | 99.4 |
| N15-LB | N | 99.8 | 100. | 100. | 100. | 99.7 | 99.8 | 99.4 | 100. |
| N15-LF | N | 99.7 | 99.7 | 99.4 | 99.4 | 100. | 100. | 99.8 | 100. |
| O117-B1c | Orf1ab | 99.2 | 99.2 | 99.7 | 100. | 93.3 | 99.5 | 99.7 | 99.7 |
| O117-B2 | Orf1ab | 99.9 | 99.9 | 100. | 100. | 99.7 | 99.7 | 99.8 | 99.8 |
| O117-B3 | Orf1ab | 100. | 100. | 100. | 100. | 99.7 | 99.7 | 99.2 | 99.2 |
| O117-F1c | Orf1ab | 99.9 | 100. | 99.7 | 100. | 99.8 | 100. | 99.4 | 99.7 |
| O117-F2 | Orf1ab | 99.9 | 99.9 | 100. | 100. | 98.9 | 98.9 | 99.7 | 99.7 |
| O117-F3 | Orf1ab | 99.8 | 99.8 | 99.7 | 99.7 | 99.5 | 99.5 | 99.4 | 99.4 |
| O117-LB | Orf1ab | 99.9 | 99.9 | 99.4 | 99.4 | 100. | 100. | 99.7 | 99.7 |
| O117-LF | Orf1ab | 100. | 100. | 100. | 100. | 100. | 100. | 99.6 | 99.6 |
| S17-B1c | S | 92.3 | 92.4 | 100. | 100. | 99.2 | 99.7 | 97.6 | 97.9 |
| S17-B2 | S | 99.9 | 99.9 | 0.862 | 0.862 | 98.9 | 98.9 | 98.8 | 98.8 |
| S17-B3 | S | 99.9 | 99.9 | 98.0 | 98.0 | 99.8 | 99.8 | 99.8 | 99.8 |
| S17-F1c | S | 92.0 | 92.0 | 100. | 100. | 97.9 | 97.9 | 93.7 | 93.9 |
| S17-F2 | S | 90.5 | 90.5 | 99.4 | 99.4 | 97.3 | 97.3 | 93.7 | 93.7 |
| S17-F3 | S | 99.9 | 99.9 | 99.4 | 99.4 | 99.8 | 99.8 | 99.3 | 99.3 |
| S17-LB | S | 92.8 | 92.9 | 96.8 | 96.8 | 99.8 | 99.8 | 98.9 | 99.0 |
| S17-LF | S | 91.7 | 91.8 | 100. | 100. | 95.7 | 97.3 | 93.9 | 94.2 |

Table S20: Matched and mismatched coverages for Jang et al. [7] set.

| Primer | Gene Target | SARS-CoV-2 | | Alpha | | Beta | | Gamma | |
|---|---|---|---|---|---|---|---|---|---|
| | | $C_{\text{strict}}$ | $C_{\text{part.}}$ | $C_{\text{strict}}$ | $C_{\text{part.}}$ | $C_{\text{strict}}$ | $C_{\text{part.}}$ | $C_{\text{strict}}$ | $C_{\text{part.}}$ |
| E-B1c | E | 99.3 | 99.3 | 99.5 | 99.5 | 99.1 | 99.1 | 99.7 | 99.7 |
| E-B2 | E | 0.000 | 99.0 | 0.000 | 99.5 | 0.000 | 99.1 | 0.000 | 99.7 |
| E-B3 | E | 99.1 | 99.1 | 99.5 | 99.6 | 0.293 | 0.293 | 98.9 | 98.9 |
| E-BLP | E | 99.3 | 99.3 | 99.4 | 99.5 | 99.1 | 99.1 | 99.8 | 99.9 |
| E-F1c | E | 99.1 | 99.2 | 99.5 | 99.6 | 99.2 | 99.3 | 99.8 | 99.8 |
| E-F2 | E | 99.4 | 99.4 | 99.5 | 99.5 | 99.4 | 99.4 | 99.9 | 99.9 |
| E-F3 | E | 98.6 | 98.6 | 99.1 | 99.1 | 98.5 | 98.5 | 99.9 | 99.9 |
| RdRP-B1c | RdRp | 98.9 | 98.9 | 99.3 | 99.3 | 99.6 | 99.6 | 99.8 | 99.8 |
| RdRP-B2 | RdRp | 99.0 | 99.0 | 99.6 | 99.6 | 99.7 | 99.7 | 99.9 | 99.9 |
| RdRP-B3 | RdRp | 99.3 | 99.3 | 99.8 | 99.8 | 99.8 | 99.8 | 99.8 | 99.8 |
| RdRP-BLP | RdRp | 99.2 | 99.4 | 99.0 | 99.8 | 99.3 | 99.8 | 99.9 | 100. |
| RdRP-F1c | RdRp | 99.2 | 99.4 | 99.8 | 99.9 | 99.7 | 99.8 | 100. | 100. |
| RdRP-F2 | RdRp | 99.4 | 99.4 | 99.7 | 99.7 | 99.3 | 99.3 | 99.6 | 99.6 |
| RdRP-F3 | RdRp | 98.6 | 98.6 | 99.8 | 99.8 | 99.2 | 99.2 | 99.2 | 99.2 |
| RdRP-FLP | RdRp | 99.1 | 99.1 | 99.6 | 99.6 | 99.2 | 99.2 | 97.8 | 97.8 |

Table S21: Matched and mismatched coverages for Jang et al. [7] set.

| Primer | Gene Target | Delta | | Lambda | | Mu | | Omicron | |
|---|---|---|---|---|---|---|---|---|---|
| | | $C_{\text{strict}}$ | $C_{\text{part.}}$ | $C_{\text{strict}}$ | $C_{\text{part.}}$ | $C_{\text{strict}}$ | $C_{\text{part.}}$ | $C_{\text{strict}}$ | $C_{\text{part.}}$ |
| E-B1c | E | 99.9 | 99.9 | 99.2 | 99.2 | 99.5 | 99.5 | 94.4 | 94.4 |
| E-B2 | E | 0.000 | 99.6 | 0.000 | 98.4 | 0.000 | 99.4 | 0.000 | 94.2 |
| E-B3 | E | 99.7 | 99.7 | 98.5 | 98.5 | 99.4 | 99.4 | 94.0 | 94.0 |
| E-BLP | E | 99.9 | 100. | 99.1 | 99.2 | 99.5 | 99.5 | 94.3 | 94.3 |
| E-F1c | E | 99.8 | 99.9 | 99.1 | 99.2 | 99.4 | 99.5 | 97.0 | 97.0 |
| E-F2 | E | 99.9 | 99.9 | 99.7 | 99.7 | 99.9 | 99.9 | 0.185 | 0.185 |
| E-F3 | E | 99.9 | 99.9 | 99.7 | 99.7 | 99.7 | 99.7 | 99.6 | 99.6 |
| RdRP-B1c | RdRp | 99.9 | 99.9 | 99.3 | 99.3 | 99.8 | 99.8 | 99.2 | 99.2 |
| RdRP-B2 | RdRp | 99.9 | 99.9 | 99.7 | 99.7 | 99.9 | 99.9 | 99.8 | 99.8 |
| RdRP-B3 | RdRp | 99.9 | 99.9 | 99.6 | 99.6 | 99.9 | 99.9 | 99.8 | 99.8 |
| RdRP-BLP | RdRp | 99.6 | 100. | 99.4 | 99.9 | 99.5 | 99.9 | 99.7 | 99.7 |
| RdRP-F1c | RdRp | 99.9 | 99.9 | 99.8 | 99.9 | 99.7 | 99.8 | 99.9 | 99.9 |
| RdRP-F2 | RdRp | 100. | 100. | 99.7 | 99.7 | 99.9 | 99.9 | 99.9 | 99.9 |
| RdRP-F3 | RdRp | 98.8 | 98.8 | 99.9 | 99.9 | 99.8 | 99.8 | 99.8 | 99.8 |
| RdRP-FLP | RdRp | 100. | 100. | 98.6 | 98.6 | 99.5 | 99.5 | 99.9 | 99.9 |

Table S22: Matched and mismatched coverages for Jang et al. [7] set.

| Primer | Gene Target | BA.2 | | BA.3 | | BA.4 | | BA.5 | |
|---|---|---|---|---|---|---|---|---|---|
| | | $C_{\text{strict}}$ | $C_{\text{part.}}$ | $C_{\text{strict}}$ | $C_{\text{part.}}$ | $C_{\text{strict}}$ | $C_{\text{part.}}$ | $C_{\text{strict}}$ | $C_{\text{part.}}$ |
| E-B1c | E | 100. | 100. | 97.7 | 97.7 | 98.6 | 98.6 | 98.1 | 98.1 |
| E-B2 | E | 0.000 | 99.9 | 0.000 | 98.0 | 0.000 | 99.2 | 0.000 | 99.8 |
| E-B3 | E | 99.8 | 99.8 | 97.7 | 97.7 | 99.4 | 99.4 | 99.4 | 99.4 |
| E-BLP | E | 100. | 100. | 97.7 | 97.7 | 99.2 | 99.4 | 97.9 | 99.9 |
| E-F1c | E | 99.9 | 99.9 | 97.7 | 97.7 | 100. | 100. | 99.4 | 99.5 |
| E-F2 | E | 0.000 | 0.000 | 0.000 | 0.000 | 0.159 | 0.159 | 0.162 | 0.162 |
| E-F3 | E | 99.9 | 99.9 | 100. | 100. | 99.7 | 99.7 | 99.6 | 99.6 |
| RdRP-B1c | RdRp | 0.0406 | 0.0406 | 0.575 | 0.575 | 0.000 | 0.000 | 0.0812 | 0.0812 |
| RdRP-B2 | RdRp | 100. | 100. | 100. | 100. | 99.7 | 99.7 | 99.9 | 99.9 |
| RdRP-B3 | RdRp | 100. | 100. | 100. | 100. | 100. | 100. | 99.9 | 99.9 |
| RdRP-BLP | RdRp | 99.7 | 100. | 100. | 100. | 100. | 100. | 99.5 | 100. |
| RdRP-F1c | RdRp | 100. | 100. | 100. | 100. | 98.9 | 100. | 98.9 | 99.6 |
| RdRP-F2 | RdRp | 100. | 100. | 100. | 100. | 99.7 | 99.7 | 98.9 | 98.9 |
| RdRP-F3 | RdRp | 99.9 | 99.9 | 100. | 100. | 99.8 | 99.8 | 99.8 | 99.8 |
| RdRP-FLP | RdRp | 99.7 | 99.7 | 100. | 100. | 100. | 100. | 99.5 | 99.5 |

Table S23: Matched and mismatched coverages for Ji et al. [8] set.

| Primer | Gene Target | SARS-CoV-2 | | Alpha | | Beta | | Gamma | |
|---|---|---|---|---|---|---|---|---|---|
| | | $C_{\text{strict}}$ | $C_{\text{part.}}$ | $C_{\text{strict}}$ | $C_{\text{part.}}$ | $C_{\text{strict}}$ | $C_{\text{part.}}$ | $C_{\text{strict}}$ | $C_{\text{part.}}$ |
| N1-B1c | N | 0.000 | 99.4 | 0.000 | 99.9 | 0.000 | 97.8 | 0.000 | 99.7 |
| N1-B2 | N | 99.4 | 99.4 | 99.7 | 99.7 | 98.1 | 98.1 | 99.5 | 99.5 |
| N1-B3 | N | 99.4 | 99.4 | 99.8 | 99.8 | 98.3 | 98.3 | 99.5 | 99.5 |
| N1-F1c | N | 0.000 | 99.0 | 0.000 | 99.8 | 0.000 | 98.4 | 0.000 | 98.7 |
| N1-F2 | N | 99.3 | 99.3 | 99.8 | 99.8 | 99.1 | 99.1 | 99.0 | 99.0 |
| N1-F3 | N | 99.2 | 99.3 | 99.1 | 99.2 | 98.3 | 98.5 | 98.8 | 98.8 |
| N1-LB | N | 99.4 | 99.4 | 99.6 | 99.6 | 97.8 | 97.8 | 0.780 | 0.780 |
| N1-LF | N | 99.3 | 99.3 | 99.9 | 99.9 | 98.6 | 98.6 | 99.2 | 99.2 |
| N2-B1c | N | 0.000 | 65.2 | 0.000 | 0.0690 | 0.000 | 96.4 | 0.000 | 0.477 |
| N2-B2 | N | 98.7 | 98.7 | 99.4 | 99.4 | 96.6 | 96.6 | 99.5 | 99.5 |
| N2-B3 | N | 98.1 | 98.1 | 98.4 | 98.4 | 96.5 | 96.5 | 99.7 | 99.7 |
| N2-F1c | N | 0.000 | 99.1 | 0.000 | 99.6 | 0.000 | 96.7 | 0.000 | 99.9 |
| N2-F2 | N | 99.2 | 99.2 | 97.0 | 97.2 | 97.5 | 97.5 | 99.7 | 99.7 |
| N2-F3 | N | 99.1 | 99.1 | 99.0 | 99.0 | 97.8 | 97.8 | 99.7 | 99.7 |
| N2-LB | N | 97.9 | 98.3 | 98.6 | 98.7 | 96.6 | 96.7 | 99.4 | 99.5 |
| N2-LF | N | 99.1 | 99.1 | 99.4 | 99.5 | 96.4 | 96.6 | 99.5 | 99.8 |
| ORFlab-1-B1c | Orf1ab | 99.3 | 99.4 | 99.7 | 99.8 | 98.8 | 98.9 | 99.0 | 99.2 |
| ORFlab-1-B2 | Orf1ab | 99.5 | 99.5 | 99.8 | 99.8 | 97.8 | 97.8 | 99.3 | 99.3 |
| ORFlab-1-B3 | Orf1ab | 99.0 | 99.0 | 99.5 | 99.5 | 95.8 | 95.8 | 99.1 | 99.1 |
| ORFlab-1-F1c | Orf1ab | 0.000 | 99.2 | 0.000 | 99.5 | 0.000 | 96.8 | 0.000 | 99.1 |
| ORFlab-1-F2 | Orf1ab | 99.3 | 99.3 | 0.152 | 0.152 | 98.8 | 98.8 | 99.2 | 99.2 |
| ORFlab-1-F3 | Orf1ab | 99.3 | 99.3 | 98.3 | 98.3 | 98.8 | 98.8 | 99.4 | 99.4 |
| ORFlab-1-LB | Orf1ab | 0.000 | 0.000 | 0.000 | 0.000 | 0.000 | 0.000 | 0.000 | 0.000 |
| ORFlab-1-LF | Orf1ab | 99.4 | 99.4 | 99.8 | 99.8 | 98.8 | 98.8 | 99.4 | 99.4 |
| ORFlab-2-B1c | Orf1ab | 99.3 | 99.3 | 99.8 | 99.8 | 99.5 | 99.6 | 99.9 | 99.9 |
| ORFlab-2-B2 | Orf1ab | 99.4 | 99.4 | 99.7 | 99.7 | 99.5 | 99.5 | 98.0 | 98.0 |
| ORFlab-2-B3 | Orf1ab | 99.4 | 99.4 | 99.6 | 99.6 | 99.8 | 99.8 | 99.8 | 99.8 |
| ORFlab-2-F1c | Orf1ab | 98.3 | 99.3 | 98.5 | 99.8 | 99.0 | 99.7 | 99.6 | 99.9 |
| ORFlab-2-F2 | Orf1ab | 99.2 | 99.2 | 99.7 | 99.7 | 99.6 | 99.6 | 99.4 | 99.4 |
| ORFlab-2-F3 | Orf1ab | 99.3 | 99.3 | 99.7 | 99.7 | 99.5 | 99.5 | 99.8 | 99.8 |
| ORFlab-2-LB | Orf1ab | 98.3 | 98.9 | 99.8 | 99.8 | 99.7 | 99.7 | 100. | 100. |
| ORFlab-2-LF | Orf1ab | 96.3 | 96.3 | 99.7 | 99.7 | 99.6 | 99.6 | 99.9 | 99.9 |

Table S24: Matched and mismatched coverages for Ji et al. [8] set.

| Primer | Gene Target | Delta | | Lambda | | Mu | | Omicron | |
|---|---|---|---|---|---|---|---|---|---|
| | | $C_{\text{strict}}$ | $C_{\text{part.}}$ | $C_{\text{strict}}$ | $C_{\text{part.}}$ | $C_{\text{strict}}$ | $C_{\text{part.}}$ | $C_{\text{strict}}$ | $C_{\text{part.}}$ |
| N1-B1c | N | 0.000 | 100. | 0.000 | 99.9 | 0.000 | 99.9 | 0.000 | 94.0 |
| N1-B2 | N | 99.9 | 99.9 | 99.2 | 99.2 | 99.5 | 99.5 | 94.6 | 94.6 |
| N1-B3 | N | 99.9 | 99.9 | 99.9 | 99.9 | 99.5 | 99.5 | 94.6 | 94.6 |
| N1-F1c | N | 0.000 | 99.8 | 0.000 | 99.5 | 0.000 | 89.0 | 0.000 | 93.4 |
| N1-F2 | N | 98.8 | 98.8 | 98.5 | 98.5 | 99.7 | 99.7 | 94.4 | 94.4 |
| N1-F3 | N | 99.8 | 99.8 | 99.3 | 99.6 | 99.5 | 99.5 | 94.6 | 94.6 |
| N1-LB | N | 99.9 | 99.9 | 99.5 | 99.5 | 97.0 | 97.0 | 93.8 | 93.8 |
| N1-LF | N | 99.8 | 99.8 | 99.4 | 99.4 | 99.7 | 99.7 | 94.9 | 94.9 |
| N2-B1c | N | 0.000 | 0.0126 | 0.000 | 0.161 | 0.000 | 99.3 | 0.000 | 0.0854 |
| N2-B2 | N | 99.9 | 99.9 | 99.7 | 99.7 | 99.8 | 99.8 | 99.1 | 99.1 |
| N2-B3 | N | 99.8 | 99.8 | 99.4 | 99.4 | 99.5 | 99.5 | 99.1 | 99.1 |
| N2-F1c | N | 0.000 | 100. | 0.000 | 99.9 | 0.000 | 99.9 | 0.000 | 99.3 |
| N2-F2 | N | 99.8 | 99.9 | 99.0 | 99.0 | 99.6 | 99.6 | 99.3 | 99.3 |
| N2-F3 | N | 99.4 | 99.4 | 99.4 | 99.4 | 99.7 | 99.7 | 99.2 | 99.2 |
| N2-LB | N | 99.6 | 99.7 | 99.3 | 99.5 | 99.4 | 99.7 | 99.2 | 99.2 |
| N2-LF | N | 99.8 | 99.9 | 99.4 | 99.5 | 99.9 | 99.9 | 99.1 | 99.2 |
| ORFlab-1-B1c | Orf1ab | 99.8 | 100. | 99.8 | 99.9 | 99.7 | 99.8 | 99.8 | 99.8 |
| ORFlab-1-B2 | Orf1ab | 100. | 100. | 99.8 | 99.8 | 97.5 | 97.5 | 98.6 | 98.6 |
| ORFlab-1-B3 | Orf1ab | 100. | 100. | 99.2 | 99.2 | 99.2 | 99.2 | 98.5 | 98.5 |
| ORFlab-1-F1c | Orf1ab | 0.000 | 99.8 | 0.000 | 99.4 | 0.000 | 99.7 | 0.000 | 99.7 |
| ORFlab-1-F2 | Orf1ab | 100. | 100. | 99.7 | 99.7 | 99.4 | 99.4 | 99.8 | 99.8 |
| ORFlab-1-F3 | Orf1ab | 100. | 100. | 99.6 | 99.6 | 98.9 | 98.9 | 99.7 | 99.7 |
| ORFlab-1-LB | Orf1ab | 0.000 | 0.000 | 0.000 | 0.000 | 0.000 | 0.000 | 0.000 | 0.000 |
| ORFlab-1-LF | Orf1ab | 100. | 100. | 99.8 | 99.8 | 99.8 | 99.8 | 99.7 | 99.7 |
| ORFlab-2-B1c | Orf1ab | 100. | 100. | 99.6 | 99.9 | 99.9 | 99.9 | 99.6 | 99.6 |
| ORFlab-2-B2 | Orf1ab | 99.4 | 99.4 | 99.7 | 99.7 | 99.5 | 99.5 | 99.7 | 99.7 |
| ORFlab-2-B3 | Orf1ab | 99.5 | 99.5 | 99.7 | 99.7 | 99.7 | 99.7 | 99.7 | 99.7 |
| ORFlab-2-F1c | Orf1ab | 99.8 | 100. | 99.0 | 99.9 | 82.1 | 99.9 | 99.0 | 99.1 |
| ORFlab-2-F2 | Orf1ab | 99.8 | 99.8 | 99.8 | 99.8 | 99.8 | 99.8 | 99.0 | 99.0 |
| ORFlab-2-F3 | Orf1ab | 100. | 100. | 99.9 | 99.9 | 99.5 | 99.5 | 98.9 | 98.9 |
| ORFlab-2-LB | Orf1ab | 100. | 100. | 99.9 | 99.9 | 99.4 | 99.4 | 99.8 | 99.8 |
| ORFlab-2-LF | Orf1ab | 99.8 | 99.9 | 99.8 | 99.8 | 99.9 | 99.9 | 99.1 | 99.1 |

Table S25: Matched and mismatched coverages for Ji et al. [8] set.

| Primer | Gene Target | BA.2 | | BA.3 | | BA.4 | | BA.5 | |
|---|---|---|---|---|---|---|---|---|---|
| | | $C_{\text{strict}}$ | $C_{\text{part.}}$ | $C_{\text{strict}}$ | $C_{\text{part.}}$ | $C_{\text{strict}}$ | $C_{\text{part.}}$ | $C_{\text{strict}}$ | $C_{\text{part.}}$ |
| N1-B1c | N | 0.000 | 99.8 | 0.000 | 94.3 | 0.000 | 99.0 | 0.000 | 99.0 |
| N1-B2 | N | 98.4 | 98.4 | 95.7 | 95.7 | 99.2 | 99.2 | 98.6 | 98.6 |
| N1-B3 | N | 100. | 100. | 95.7 | 95.7 | 99.7 | 99.7 | 99.6 | 99.6 |
| N1-F1c | N | 0.000 | 98.9 | 0.000 | 95.1 | 0.000 | 67.7 | 0.000 | 80.4 |
| N1-F2 | N | 99.8 | 99.8 | 95.4 | 95.4 | 98.7 | 98.7 | 89.5 | 89.5 |
| N1-F3 | N | 99.5 | 99.5 | 95.1 | 95.1 | 100. | 100. | 99.5 | 99.6 |
| N1-LB | N | 99.8 | 99.8 | 94.3 | 94.3 | 98.7 | 99.0 | 98.6 | 98.7 |
| N1-LF | N | 99.9 | 99.9 | 95.7 | 95.7 | 99.5 | 99.5 | 99.1 | 99.1 |
| N2-B1c | N | 0.000 | 0.0947 | 0.000 | 0.000 | 0.000 | 0.000 | 0.000 | 0.000 |
| N2-B2 | N | 99.9 | 99.9 | 100. | 100. | 99.5 | 99.5 | 99.8 | 99.8 |
| N2-B3 | N | 99.1 | 99.1 | 98.9 | 98.9 | 99.0 | 99.0 | 99.5 | 99.5 |
| N2-F1c | N | 0.000 | 100. | 0.000 | 99.7 | 0.000 | 99.4 | 0.000 | 99.4 |
| N2-F2 | N | 99.9 | 99.9 | 99.7 | 99.7 | 99.7 | 99.7 | 99.6 | 99.7 |
| N2-F3 | N | 99.2 | 99.2 | 99.7 | 99.7 | 0.954 | 0.954 | 97.4 | 97.4 |
| N2-LB | N | 99.6 | 99.7 | 100. | 100. | 100. | 100. | 99.8 | 99.9 |
| N2-LF | N | 99.8 | 99.8 | 98.9 | 99.4 | 100. | 100. | 99.9 | 100. |
| ORFlab-1-B1c | Orf1ab | 100. | 100. | 99.7 | 99.7 | 100. | 100. | 99.8 | 99.9 |
| ORFlab-1-B2 | Orf1ab | 99.9 | 99.9 | 99.1 | 99.1 | 100. | 100. | 99.9 | 99.9 |
| ORFlab-1-B3 | Orf1ab | 99.9 | 99.9 | 99.7 | 99.7 | 100. | 100. | 99.5 | 99.5 |
| ORFlab-1-F1c | Orf1ab | 0.000 | 99.9 | 0.000 | 99.4 | 0.000 | 100. | 0.000 | 99.9 |
| ORFlab-1-F2 | Orf1ab | 99.9 | 99.9 | 99.4 | 99.4 | 98.6 | 98.6 | 99.9 | 99.9 |
| ORFlab-1-F3 | Orf1ab | 99.9 | 99.9 | 99.4 | 99.4 | 100. | 100. | 100. | 100. |
| ORFlab-1-LB | Orf1ab | 0.000 | 0.000 | 0.000 | 0.000 | 0.000 | 0.000 | 0.000 | 0.000 |
| ORFlab-1-LF | Orf1ab | 99.3 | 99.3 | 99.1 | 99.1 | 100. | 100. | 99.5 | 99.5 |
| ORFlab-2-B1c | Orf1ab | 100. | 100. | 99.7 | 99.7 | 100. | 100. | 99.8 | 99.8 |
| ORFlab-2-B2 | Orf1ab | 99.8 | 99.8 | 100. | 100. | 99.7 | 99.7 | 99.6 | 99.6 |
| ORFlab-2-B3 | Orf1ab | 99.9 | 99.9 | 100. | 100. | 99.8 | 99.8 | 100. | 100. |
| ORFlab-2-F1c | Orf1ab | 99.4 | 99.9 | 99.4 | 99.7 | 99.5 | 100. | 99.8 | 99.9 |
| ORFlab-2-F2 | Orf1ab | 99.9 | 99.9 | 99.7 | 99.7 | 99.8 | 99.8 | 100. | 100. |
| ORFlab-2-F3 | Orf1ab | 100. | 100. | 99.7 | 99.7 | 99.4 | 99.4 | 92.1 | 92.1 |
| ORFlab-2-LB | Orf1ab | 100. | 100. | 100. | 100. | 100. | 100. | 99.5 | 99.5 |
| ORFlab-2-LF | Orf1ab | 99.9 | 99.9 | 99.7 | 99.7 | 100. | 100. | 99.8 | 99.8 |

Table S26: Matched and mismatched coverages for Jiang et al. [9] set.

| Primer | Gene Target | SARS-CoV-2 | | Alpha | | Beta | | Gamma | |
|---|---|---|---|---|---|---|---|---|---|
| | | $C_{\text{strict}}$ | $C_{\text{part.}}$ | $C_{\text{strict}}$ | $C_{\text{part.}}$ | $C_{\text{strict}}$ | $C_{\text{part.}}$ | $C_{\text{strict}}$ | $C_{\text{part.}}$ |
| nCoV-N-B1c | N | 97.9 | 98.4 | 99.4 | 99.8 | 97.1 | 97.1 | 99.6 | 99.6 |
| nCoV-N-B2 | N | 99.4 | 99.4 | 99.7 | 99.7 | 98.1 | 98.1 | 99.5 | 99.5 |
| nCoV-N-B3 | N | 99.4 | 99.4 | 99.8 | 99.8 | 98.3 | 98.3 | 99.6 | 99.6 |
| nCoV-N-F1c | N | 99.6 | 99.6 | 99.9 | 100. | 99.4 | 99.5 | 99.8 | 99.9 |
| nCoV-N-F2 | N | 99.2 | 99.2 | 99.8 | 99.8 | 98.4 | 98.4 | 98.7 | 98.7 |
| nCoV-N-F3 | N | 99.0 | 99.1 | 99.8 | 99.8 | 98.4 | 98.4 | 98.9 | 98.9 |
| nCoV-N-LB | N | 99.5 | 99.5 | 99.6 | 99.8 | 97.8 | 98.0 | 0.780 | 99.6 |
| nCoV-N-LF | N | 99.4 | 99.4 | 99.5 | 99.5 | 99.1 | 99.1 | 99.2 | 99.2 |

Table S27: Matched and mismatched coverages for Jiang et al. [9] set.

| Primer | Gene Target | Delta | | Lambda | | Mu | | Omicron | |
|---|---|---|---|---|---|---|---|---|---|
| | | $C_{\text{strict}}$ | $C_{\text{part.}}$ | $C_{\text{strict}}$ | $C_{\text{part.}}$ | $C_{\text{strict}}$ | $C_{\text{part.}}$ | $C_{\text{strict}}$ | $C_{\text{part.}}$ |
| nCoV-N-B1c | N | 99.5 | 99.8 | 99.6 | 99.7 | 97.1 | 99.3 | 93.8 | 93.8 |
| nCoV-N-B2 | N | 99.9 | 99.9 | 99.2 | 99.2 | 99.5 | 99.5 | 94.5 | 94.5 |
| nCoV-N-B3 | N | 99.9 | 99.9 | 99.9 | 99.9 | 99.1 | 99.1 | 94.6 | 94.6 |
| nCoV-N-F1c | N | 99.8 | 99.9 | 94.8 | 95.0 | 99.5 | 99.8 | 94.9 | 94.9 |
| nCoV-N-F2 | N | 99.8 | 99.8 | 99.2 | 99.2 | 88.8 | 88.8 | 94.9 | 94.9 |
| nCoV-N-F3 | N | 99.8 | 99.9 | 99.5 | 99.6 | 99.1 | 99.2 | 0.555 | 0.555 |
| nCoV-N-LB | N | 99.9 | 100. | 99.5 | 99.6 | 97.1 | 97.2 | 93.8 | 93.8 |
| nCoV-N-LF | N | 99.7 | 99.7 | 99.6 | 99.6 | 99.7 | 99.7 | 94.8 | 94.8 |

Table S28: Matched and mismatched coverages for Jiang et al. [9] set.

| | | BA.2 | | BA.3 | | BA.4 | | BA.5 | |
|---|---|---|---|---|---|---|---|---|---|
| Primer | Gene Target | $C_{\text{strict}}$ | $C_{\text{part.}}$ | $C_{\text{strict}}$ | $C_{\text{part.}}$ | $C_{\text{strict}}$ | $C_{\text{part.}}$ | $C_{\text{strict}}$ | $C_{\text{part.}}$ |
| nCoV-N-B1c | N | 99.6 | 99.8 | 94.3 | 94.3 | 99.0 | 99.0 | 98.7 | 98.7 |
| nCoV-N-B2 | N | 98.4 | 98.4 | 95.7 | 95.7 | 99.2 | 99.2 | 98.6 | 98.6 |
| nCoV-N-B3 | N | 100. | 100. | 95.7 | 95.7 | 99.7 | 99.7 | 98.5 | 98.5 |
| nCoV-N-F1c | N | 98.1 | 98.1 | 89.4 | 89.9 | 99.2 | 99.4 | 96.7 | 98.5 |
| nCoV-N-F2 | N | 99.6 | 99.6 | 96.0 | 96.0 | 99.7 | 99.7 | 98.9 | 98.9 |
| nCoV-N-F3 | N | 1.31 | 1.31 | 0.862 | 0.862 | 0.318 | 0.318 | 0.975 | 0.975 |
| nCoV-N-LB | N | 99.8 | 99.8 | 94.3 | 94.3 | 98.7 | 98.7 | 98.8 | 99.0 |
| nCoV-N-LF | N | 99.9 | 99.9 | 96.0 | 96.0 | 99.5 | 99.5 | 99.4 | 99.4 |

Table S29: Matched and mismatched coverages for Lalli et al. [10] set.

| | | SARS-CoV-2 | | Alpha | | Beta | | Gamma | |
|---|---|---|---|---|---|---|---|---|---|
| Primer | Gene Target | $C_{\text{strict}}$ | $C_{\text{part.}}$ | $C_{\text{strict}}$ | $C_{\text{part.}}$ | $C_{\text{strict}}$ | $C_{\text{part.}}$ | $C_{\text{strict}}$ | $C_{\text{part.}}$ |
| NEB_E1-B1c | E | 99.2 | 99.3 | 99.6 | 99.7 | 99.3 | 99.3 | 99.8 | 99.8 |
| NEB_E1-B2 | E | 99.3 | 99.3 | 99.4 | 99.4 | 99.2 | 99.2 | 99.9 | 99.9 |
| NEB_E1-B3 | E | 99.0 | 99.0 | 99.6 | 99.6 | 98.9 | 98.9 | 99.8 | 99.8 |
| NEB_E1-F1c | E | 99.2 | 99.3 | 99.6 | 99.6 | 99.4 | 99.4 | 99.8 | 99.8 |
| NEB_E1-F2 | E | 98.5 | 98.5 | 98.9 | 98.9 | 98.5 | 98.5 | 99.9 | 99.9 |
| NEB_E1-F3 | E | 98.5 | 98.5 | 99.8 | 99.8 | 99.7 | 99.7 | 100. | 100. |
| NEB_E1-LB | E | 99.3 | 99.3 | 99.5 | 99.6 | 99.1 | 99.2 | 99.7 | 99.8 |
| NEB_E1-LF | E | 99.5 | 99.5 | 99.7 | 99.7 | 99.4 | 99.4 | 99.9 | 99.9 |
| NEB_N2-B1c | N | 99.3 | 99.3 | 99.7 | 99.7 | 97.7 | 97.7 | 99.8 | 99.8 |
| NEB_N2-B2 | N | 99.2 | 99.2 | 98.0 | 98.0 | 97.5 | 97.5 | 99.6 | 99.6 |
| NEB_N2-B3 | N | 99.2 | 99.3 | 99.6 | 99.6 | 97.8 | 97.8 | 98.7 | 98.7 |
| NEB_N2-F1c | N | 99.2 | 99.2 | 99.6 | 99.7 | 98.4 | 98.4 | 99.5 | 99.9 |
| NEB_N2-F2 | N | 99.2 | 99.2 | 99.5 | 99.5 | 98.2 | 98.2 | 99.0 | 99.0 |
| NEB_N2-F3 | N | 99.0 | 99.0 | 99.5 | 99.5 | 98.2 | 98.2 | 99.6 | 99.6 |
| NEB_N2-LB | N | 98.8 | 98.8 | 99.3 | 99.4 | 97.4 | 97.4 | 99.4 | 99.5 |
| NEB_N2-LF | N | 99.0 | 99.0 | 98.5 | 98.7 | 98.1 | 98.1 | 99.8 | 99.8 |
| NEB_geneN-A-B1c | N | 98.7 | 98.8 | 99.6 | 99.7 | 96.9 | 97.6 | 99.5 | 99.6 |
| NEB_geneN-A-B2 | N | 99.1 | 99.1 | 99.7 | 99.7 | 97.2 | 97.2 | 99.6 | 99.6 |
| NEB_geneN-A-B3 | N | 99.1 | 99.1 | 96.1 | 96.1 | 97.5 | 97.5 | 99.7 | 99.7 |
| NEB_geneN-A-F1c | N | 99.4 | 99.5 | 99.8 | 99.9 | 98.0 | 98.1 | 98.1 | 98.2 |
| NEB_geneN-A-F2 | N | 99.4 | 99.4 | 99.8 | 99.8 | 98.3 | 98.3 | 99.5 | 99.5 |
| NEB_geneN-A-F3 | N | 99.4 | 99.4 | 99.7 | 99.7 | 98.1 | 98.1 | 99.5 | 99.5 |
| NEB_geneN-A-LB | N | 98.9 | 99.2 | 97.9 | 97.9 | 97.3 | 97.3 | 99.5 | 99.5 |
| NEB_geneN-A-LF | N | 99.4 | 99.5 | 99.8 | 99.9 | 98.2 | 98.3 | 99.6 | 99.6 |
| NEB_orf1a-A-B1c | Orf1a | 98.7 | 98.7 | 99.6 | 99.7 | 99.4 | 99.5 | 99.7 | 99.7 |
| NEB_orf1a-A-B2 | Orf1a | 98.7 | 98.7 | 99.7 | 99.7 | 99.3 | 99.3 | 99.0 | 99.0 |
| NEB_orf1a-A-B3 | Orf1a | 97.7 | 97.7 | 99.8 | 99.8 | 98.9 | 98.9 | 99.8 | 99.8 |
| NEB_orf1a-A-F1c | Orf1a | 0.000 | 98.9 | 0.000 | 99.7 | 0.000 | 99.3 | 0.000 | 99.8 |
| NEB_orf1a-A-F2 | Orf1a | 98.6 | 98.6 | 99.3 | 99.3 | 98.9 | 98.9 | 99.6 | 99.6 |
| NEB_orf1a-A-F3 | Orf1a | 98.2 | 98.2 | 99.5 | 99.5 | 98.7 | 98.7 | 99.1 | 99.1 |
| NEB_orf1a-A-LB | Orf1a | 98.6 | 98.7 | 99.6 | 99.6 | 99.4 | 99.5 | 99.7 | 99.7 |
| NEB_orf1a-A-LF | Orf1a | 98.9 | 98.9 | 99.6 | 99.7 | 98.2 | 98.2 | 98.9 | 99.7 |

Table S30: Matched and mismatched coverages for Lalli et al. [10] set.

| | | Delta | | Lambda | | Mu | | Omicron | |
|---|---|---|---|---|---|---|---|---|---|
| Primer | Gene Target | $C_{\text{strict}}$ | $C_{\text{part.}}$ | $C_{\text{strict}}$ | $C_{\text{part.}}$ | $C_{\text{strict}}$ | $C_{\text{part.}}$ | $C_{\text{strict}}$ | $C_{\text{part.}}$ |
| NEB_E1-B1c | E | 99.9 | 99.9 | 99.2 | 99.2 | 99.5 | 99.5 | 94.6 | 94.6 |
| NEB_E1-B2 | E | 100. | 100. | 99.0 | 99.0 | 99.5 | 99.5 | 94.3 | 94.3 |
| NEB_E1-B3 | E | 99.6 | 99.6 | 98.7 | 98.7 | 99.6 | 99.6 | 94.2 | 94.2 |
| NEB_E1-F1c | E | 99.9 | 99.9 | 99.0 | 99.0 | 99.5 | 99.5 | 97.1 | 97.1 |
| NEB_E1-F2 | E | 99.8 | 99.8 | 99.6 | 99.6 | 99.7 | 99.7 | 0.199 | 0.199 |
| NEB_E1-F3 | E | 100. | 100. | 100. | 100. | 99.9 | 99.9 | 99.7 | 99.7 |
| NEB_E1-LB | E | 99.9 | 99.9 | 99.2 | 99.2 | 99.4 | 99.4 | 94.3 | 94.3 |
| NEB_E1-LF | E | 100. | 100. | 99.9 | 99.9 | 99.9 | 99.9 | 97.5 | 97.5 |
| NEB_N2-B1c | N | 99.9 | 99.9 | 98.9 | 98.9 | 99.8 | 99.8 | 99.3 | 99.3 |
| NEB_N2-B2 | N | 99.8 | 99.8 | 98.8 | 98.8 | 99.3 | 99.3 | 99.5 | 99.5 |
| NEB_N2-B3 | N | 99.8 | 99.9 | 1.03 | 1.03 | 98.9 | 98.9 | 86.5 | 99.1 |
| NEB_N2-F1c | N | 99.9 | 100. | 98.9 | 98.9 | 99.5 | 99.6 | 99.4 | 99.4 |
| NEB_N2-F2 | N | 99.8 | 99.8 | 98.9 | 99.0 | 99.5 | 99.5 | 99.3 | 99.3 |
| NEB_N2-F3 | N | 99.8 | 99.8 | 98.9 | 98.9 | 99.6 | 99.6 | 99.4 | 99.4 |
| NEB_N2-LB | N | 98.7 | 98.7 | 98.8 | 99.0 | 98.9 | 99.0 | 99.5 | 99.5 |
| NEB_N2-LF | N | 100. | 100. | 99.3 | 99.3 | 99.0 | 99.0 | 99.4 | 99.4 |
| NEB_geneN-A-B1c | N | 99.9 | 99.9 | 99.3 | 99.4 | 85.5 | 85.7 | 94.2 | 94.3 |
| NEB_geneN-A-B2 | N | 99.5 | 99.5 | 97.7 | 97.7 | 99.5 | 99.5 | 99.2 | 99.2 |
| NEB_geneN-A-B3 | N | 99.5 | 99.5 | 99.3 | 99.3 | 99.6 | 99.6 | 99.2 | 99.2 |
| NEB_geneN-A-F1c | N | 99.9 | 99.9 | 99.6 | 100. | 85.2 | 99.7 | 94.2 | 94.4 |
| NEB_geneN-A-F2 | N | 99.9 | 99.9 | 99.9 | 99.9 | 99.5 | 99.5 | 94.6 | 94.6 |
| NEB_geneN-A-F3 | N | 99.9 | 99.9 | 99.2 | 99.2 | 99.6 | 99.6 | 94.6 | 94.6 |
| NEB_geneN-A-LB | N | 99.4 | 99.5 | 99.4 | 99.5 | 99.6 | 99.6 | 94.8 | 94.8 |
| NEB_geneN-A-LF | N | 99.9 | 99.9 | 99.8 | 99.9 | 99.1 | 99.6 | 94.4 | 94.5 |
| NEB_orf1a-A-B1c | Orf1a | 99.8 | 99.9 | 99.7 | 99.8 | 99.5 | 99.6 | 99.4 | 99.4 |
| NEB_orf1a-A-B2 | Orf1a | 98.0 | 98.0 | 99.4 | 99.4 | 97.4 | 97.4 | 99.0 | 99.0 |
| NEB_orf1a-A-B3 | Orf1a | 98.0 | 98.0 | 99.6 | 99.6 | 98.8 | 98.8 | 99.4 | 99.4 |
| NEB_orf1a-A-F1c | Orf1a | 0.000 | 99.8 | 0.000 | 99.9 | 0.000 | 99.6 | 0.0142 | 99.5 |
| NEB_orf1a-A-F2 | Orf1a | 97.9 | 97.9 | 99.4 | 99.4 | 94.4 | 94.4 | 98.6 | 98.6 |
| NEB_orf1a-A-F3 | Orf1a | 98.4 | 98.4 | 98.3 | 98.3 | 93.5 | 93.5 | 98.9 | 98.9 |
| NEB_orf1a-A-LB | Orf1a | 99.9 | 99.9 | 99.6 | 99.7 | 99.7 | 99.8 | 99.4 | 99.5 |
| NEB_orf1a-A-LF | Orf1a | 99.8 | 99.9 | 99.6 | 99.9 | 99.5 | 99.8 | 99.4 | 99.4 |

Table S31: Matched and mismatched coverages for Lalli et al. [10] set.

| Primer | Gene Target | BA.2 | | BA.3 | | BA.4 | | BA.5 | |
|---|---|---|---|---|---|---|---|---|---|
| | | $C_{\text{strict}}$ | $C_{\text{part.}}$ | $C_{\text{strict}}$ | $C_{\text{part.}}$ | $C_{\text{strict}}$ | $C_{\text{part.}}$ | $C_{\text{strict}}$ | $C_{\text{part.}}$ |
| NEB_E1-B1c | E | 100. | 100. | 97.7 | 97.7 | 99.8 | 99.8 | 99.8 | 100. |
| NEB_E1-B2 | E | 100. | 100. | 97.7 | 97.7 | 99.5 | 99.5 | 97.8 | 97.8 |
| NEB_E1-B3 | E | 99.9 | 99.9 | 97.7 | 97.7 | 99.5 | 99.5 | 99.5 | 99.5 |
| NEB_E1-F1c | E | 99.9 | 99.9 | 98.3 | 98.3 | 100. | 100. | 99.7 | 99.7 |
| NEB_E1-F2 | E | 0.000 | 0.000 | 0.000 | 0.000 | 0.159 | 0.159 | 0.162 | 0.162 |
| NEB_E1-F3 | E | 100. | 100. | 100. | 100. | 99.8 | 99.8 | 99.4 | 99.4 |
| NEB_E1-LB | E | 100. | 100. | 97.7 | 97.7 | 98.6 | 98.6 | 98.1 | 98.8 |
| NEB_E1-LF | E | 100. | 100. | 98.9 | 98.9 | 100. | 100. | 100. | 100. |
| NEB_N2-B1c | N | 99.9 | 99.9 | 99.7 | 99.7 | 99.4 | 99.4 | 98.1 | 98.1 |
| NEB_N2-B2 | N | 99.9 | 99.9 | 99.4 | 99.4 | 97.6 | 97.6 | 99.2 | 99.2 |
| NEB_N2-B3 | N | 99.6 | 99.8 | 18.7 | 18.7 | 99.8 | 99.8 | 99.3 | 99.4 |
| NEB_N2-F1c | N | 99.9 | 99.9 | 100. | 100. | 99.8 | 99.8 | 99.8 | 99.8 |
| NEB_N2-F2 | N | 99.8 | 99.8 | 100. | 100. | 99.7 | 99.7 | 99.6 | 99.6 |
| NEB_N2-F3 | N | 99.9 | 99.9 | 100. | 100. | 99.0 | 99.0 | 99.8 | 99.8 |
| NEB_N2-LB | N | 99.7 | 99.8 | 99.7 | 99.7 | 98.7 | 98.7 | 95.1 | 95.1 |
| NEB_N2-LF | N | 99.6 | 99.6 | 99.7 | 99.7 | 100. | 100. | 99.8 | 99.8 |
| NEB_geneN-A-B1c | N | 99.7 | 99.7 | 96.3 | 96.3 | 100. | 100. | 99.7 | 99.8 |
| NEB_geneN-A-B2 | N | 99.9 | 99.9 | 99.4 | 99.4 | 98.9 | 98.9 | 99.4 | 99.4 |
| NEB_geneN-A-B3 | N | 99.2 | 99.2 | 99.7 | 99.7 | 0.954 | 0.954 | 97.6 | 97.6 |
| NEB_geneN-A-F1c | N | 100. | 100. | 95.7 | 95.7 | 100. | 100. | 99.6 | 99.7 |
| NEB_geneN-A-F2 | N | 100. | 100. | 95.7 | 95.7 | 99.7 | 99.7 | 99.6 | 99.6 |
| NEB_geneN-A-F3 | N | 98.4 | 98.4 | 95.7 | 95.7 | 99.2 | 99.2 | 98.6 | 98.6 |
| NEB_geneN-A-LB | N | 99.6 | 99.6 | 97.7 | 97.7 | 99.8 | 99.8 | 91.8 | 92.0 |
| NEB_geneN-A-LF | N | 99.8 | 100. | 95.7 | 95.7 | 99.8 | 100. | 98.2 | 99.5 |
| NEB_orf1a-A-B1c | Orf1a | 99.9 | 99.9 | 99.4 | 99.4 | 99.4 | 99.7 | 97.9 | 98.1 |
| NEB_orf1a-A-B2 | Orf1a | 0.189 | 0.189 | 0.575 | 0.575 | 0.159 | 0.159 | 0.325 | 0.325 |
| NEB_orf1a-A-B3 | Orf1a | 99.5 | 99.5 | 79.6 | 79.6 | 4.45 | 4.45 | 96.9 | 96.9 |
| NEB_orf1a-A-F1c | Orf1a | 0.000 | 99.8 | 0.000 | 99.1 | 0.000 | 99.4 | 0.000 | 97.1 |
| NEB_orf1a-A-F2 | Orf1a | 98.7 | 98.7 | 96.0 | 96.0 | 94.1 | 94.1 | 87.8 | 87.8 |
| NEB_orf1a-A-F3 | Orf1a | 99.1 | 99.1 | 95.7 | 95.7 | 96.7 | 96.7 | 87.4 | 87.4 |
| NEB_orf1a-A-LB | Orf1a | 98.8 | 99.1 | 97.1 | 97.1 | 99.5 | 99.5 | 97.8 | 97.8 |
| NEB_orf1a-A-LF | Orf1a | 99.7 | 99.8 | 99.1 | 99.1 | 99.2 | 99.2 | 97.3 | 97.3 |

Table S32: Matched and mismatched coverages for Lau et al. [11] set.

| | | SARS-CoV-2 | | Alpha | | Beta | | Gamma | |
|---|---|---|---|---|---|---|---|---|---|
| Primer | Gene Target | $C_{\text{strict}}$ | $C_{\text{part.}}$ | $C_{\text{strict}}$ | $C_{\text{part.}}$ | $C_{\text{strict}}$ | $C_{\text{part.}}$ | $C_{\text{strict}}$ | $C_{\text{part.}}$ |
| B1c | N | 98.5 | 98.6 | 99.1 | 99.1 | 92.7 | 92.7 | 98.7 | 98.7 |
| B2 | N | 99.0 | 99.0 | 99.8 | 99.8 | 99.1 | 99.1 | 98.9 | 98.9 |
| BLP | N | 99.3 | 99.3 | 99.8 | 99.8 | 99.2 | 99.2 | 99.2 | 99.2 |
| BP | N | 99.3 | 99.3 | 99.7 | 99.7 | 98.9 | 98.9 | 98.8 | 98.8 |
| BSP | N | 98.6 | 98.6 | 99.7 | 99.7 | 92.3 | 92.3 | 99.1 | 99.1 |
| F1c | N | 99.1 | 99.4 | 0.221 | 0.235 | 98.8 | 99.2 | 98.2 | 98.4 |
| F2 | N | 99.4 | 99.4 | 98.9 | 98.9 | 99.3 | 99.3 | 97.0 | 97.0 |
| FLP | N | 98.5 | 98.5 | 98.0 | 98.0 | 0.814 | 0.814 | 91.9 | 91.9 |
| FP | N | 99.1 | 99.1 | 99.6 | 99.6 | 98.7 | 98.7 | 96.7 | 96.7 |
| FSP | N | 99.1 | 99.1 | 0.235 | 0.235 | 98.9 | 98.9 | 98.4 | 98.4 |

Table S33: Matched and mismatched coverages for Lau et al. [11] set.

| Primer | Gene Target | Delta | | Lambda | | Mu | | Omicron | |
|---|---|---|---|---|---|---|---|---|---|
| | | $C_{\text{strict}}$ | $C_{\text{part.}}$ | $C_{\text{strict}}$ | $C_{\text{part.}}$ | $C_{\text{strict}}$ | $C_{\text{part.}}$ | $C_{\text{strict}}$ | $C_{\text{part.}}$ |
| B1c | N | 99.7 | 99.7 | 0.664 | 0.664 | 99.1 | 99.1 | 0.640 | 0.640 |
| B2 | N | 99.8 | 99.8 | 99.5 | 99.5 | 99.1 | 99.1 | 0.555 | 0.555 |
| BLP | N | 98.8 | 98.8 | 98.6 | 98.6 | 99.7 | 99.7 | 94.8 | 94.8 |
| BP | N | 99.8 | 99.8 | 99.3 | 99.3 | 88.8 | 88.8 | 94.9 | 94.9 |
| BSP | N | 99.7 | 99.7 | 0.664 | 0.664 | 99.2 | 99.2 | 0.640 | 0.640 |
| F1c | N | 3.78 | 3.85 | 0.0535 | 99.1 | 0.197 | 99.4 | 0.142 | 93.9 |
| F2 | N | 100. | 100. | 99.2 | 99.2 | 99.2 | 99.2 | 92.3 | 92.3 |
| FLP | N | 0.768 | 0.768 | 0.739 | 0.739 | 95.0 | 95.0 | 91.8 | 91.8 |
| FP | N | 99.7 | 99.7 | 98.5 | 98.5 | 97.6 | 97.6 | 91.8 | 91.8 |
| FSP | N | 99.8 | 99.8 | 99.3 | 99.3 | 99.5 | 99.5 | 94.6 | 94.6 |

Table S34: Matched and mismatched coverages for Lau et al. [11] set.

| Primer | Gene Target | BA.2 | | BA.3 | | BA.4 | | BA.5 | |
|---|---|---|---|---|---|---|---|---|---|
| | | $C_{\text{strict}}$ | $C_{\text{part.}}$ | $C_{\text{strict}}$ | $C_{\text{part.}}$ | $C_{\text{strict}}$ | $C_{\text{part.}}$ | $C_{\text{strict}}$ | $C_{\text{part.}}$ |
| B1c | N | 0.000 | 0.000 | 1.15 | 1.15 | 0.477 | 0.477 | 0.569 | 0.569 |
| B2 | N | 1.31 | 1.31 | 0.862 | 0.862 | 0.318 | 0.318 | 0.975 | 0.975 |
| BLP | N | 99.9 | 99.9 | 95.4 | 95.4 | 99.0 | 99.0 | 89.4 | 89.4 |
| BP | N | 99.6 | 99.6 | 96.0 | 96.0 | 99.5 | 99.5 | 98.7 | 98.7 |
| BSP | N | 0.000 | 0.000 | 1.15 | 1.15 | 0.477 | 0.477 | 0.569 | 0.569 |
| F1c | N | 0.000 | 99.5 | 0.000 | 93.4 | 1.11 | 99.5 | 1.38 | 99.5 |
| F2 | N | 99.8 | 99.8 | 95.7 | 95.7 | 99.7 | 99.7 | 96.6 | 96.6 |
| FLP | N | 99.6 | 99.6 | 94.3 | 94.3 | 95.5 | 95.5 | 89.4 | 89.4 |
| FP | N | 99.8 | 99.8 | 94.8 | 94.8 | 99.8 | 99.8 | 96.7 | 96.7 |
| FSP | N | 99.5 | 99.5 | 94.0 | 94.0 | 99.8 | 99.8 | 99.4 | 99.4 |

Table S35: Matched and mismatched coverages for Luo et al. [12] set.

| | | SARS-CoV-2 | | Alpha | | Beta | | Gamma | |
|---|---|---|---|---|---|---|---|---|---|
| Primer | Gene Target | $C_{\text{strict}}$ | $C_{\text{part.}}$ | $C_{\text{strict}}$ | $C_{\text{part.}}$ | $C_{\text{strict}}$ | $C_{\text{part.}}$ | $C_{\text{strict}}$ | $C_{\text{part.}}$ |
| PS1-B1c | N | 97.9 | 98.0 | 98.8 | 99.1 | 96.5 | 96.6 | 98.5 | 98.6 |
| PS1-B2 | N | 65.2 | 65.2 | 0.0690 | 0.0690 | 96.4 | 96.5 | 0.477 | 0.520 |
| PS1-B3 | N | 98.0 | 98.0 | 98.7 | 98.7 | 96.6 | 96.6 | 99.4 | 99.4 |
| PS1-F1c | N | 99.1 | 99.2 | 99.4 | 99.6 | 96.6 | 96.7 | 98.9 | 99.9 |
| PS1-F2 | N | 99.2 | 99.2 | 97.0 | 97.2 | 97.5 | 97.5 | 99.7 | 99.7 |
| PS1-F3 | N | 99.1 | 99.1 | 99.0 | 99.0 | 97.8 | 97.8 | 99.7 | 99.7 |
| PS1-LB | N | 97.7 | 99.2 | 99.5 | 99.6 | 96.6 | 96.8 | 97.5 | 99.8 |
| PS1-LF | N | 98.8 | 98.9 | 99.4 | 99.5 | 96.4 | 96.6 | 99.5 | 99.8 |
| PS2-B1c | N | 97.9 | 98.4 | 99.4 | 99.8 | 97.1 | 97.1 | 99.6 | 99.6 |
| PS2-B2 | N | 99.4 | 99.4 | 99.7 | 99.7 | 98.1 | 98.1 | 99.5 | 99.5 |
| PS2-B3 | N | 99.4 | 99.4 | 99.8 | 99.8 | 98.3 | 98.3 | 99.6 | 99.6 |
| PS2-F1c | N | 99.6 | 99.6 | 99.9 | 100. | 99.4 | 99.5 | 99.8 | 99.9 |
| PS2-F2 | N | 99.2 | 99.2 | 99.8 | 99.8 | 98.4 | 98.4 | 98.7 | 98.7 |
| PS2-F3 | N | 99.0 | 99.1 | 99.8 | 99.8 | 98.4 | 98.4 | 98.9 | 98.9 |
| PS2-LB | N | 99.4 | 99.5 | 99.6 | 99.8 | 97.8 | 97.9 | 0.780 | 0.780 |
| PS2-LF | N | 99.4 | 99.4 | 99.5 | 99.5 | 99.1 | 99.1 | 99.2 | 99.2 |
| PS3-B1c | N | 97.9 | 98.4 | 99.4 | 99.8 | 97.1 | 97.1 | 99.6 | 99.6 |
| PS3-B2 | N | 99.4 | 99.4 | 99.7 | 99.7 | 98.1 | 98.1 | 99.5 | 99.5 |
| PS3-B3 | N | 99.4 | 99.4 | 99.8 | 99.8 | 98.3 | 98.3 | 99.6 | 99.6 |
| PS3-F1c | N | 99.6 | 99.6 | 99.9 | 100. | 99.4 | 99.5 | 99.8 | 99.9 |
| PS3-F2 | N | 99.2 | 99.2 | 99.8 | 99.8 | 98.4 | 98.4 | 98.7 | 98.7 |
| PS3-F3 | N | 99.0 | 99.1 | 99.8 | 99.8 | 98.4 | 98.4 | 98.9 | 98.9 |
| PS3-LB | N | 99.4 | 99.4 | 99.6 | 99.6 | 97.8 | 97.8 | 0.780 | 0.780 |
| PS3-LF | N | 99.4 | 99.5 | 99.5 | 99.5 | 99.1 | 99.1 | 99.2 | 99.2 |
| PS4-B1c | N | 97.9 | 98.0 | 98.8 | 99.1 | 96.5 | 96.6 | 98.5 | 98.6 |
| PS4-B2 | N | 65.2 | 65.2 | 0.0690 | 0.0690 | 96.4 | 96.5 | 0.477 | 0.520 |
| PS4-B3 | N | 98.1 | 98.1 | 98.6 | 98.6 | 96.6 | 96.7 | 99.4 | 99.4 |
| PS4-F1c | N | 99.1 | 99.2 | 99.4 | 99.6 | 96.6 | 96.7 | 98.9 | 99.9 |
| PS4-F2 | N | 99.1 | 99.1 | 96.9 | 97.2 | 97.6 | 97.6 | 99.7 | 99.7 |
| PS4-F3 | N | 99.1 | 99.1 | 99.7 | 99.7 | 97.2 | 97.2 | 99.6 | 99.6 |
| PS4-LB | N | 98.0 | 99.0 | 99.5 | 99.6 | 96.7 | 96.8 | 97.6 | 99.9 |
| PS4-LF | N | 98.8 | 99.1 | 99.4 | 99.6 | 96.4 | 96.7 | 99.5 | 99.8 |
| PS5-B1c | N | 97.9 | 98.0 | 98.8 | 99.1 | 96.5 | 96.6 | 98.5 | 98.6 |
| PS5-B2 | N | 65.2 | 65.2 | 0.0690 | 0.0690 | 96.4 | 96.5 | 0.477 | 0.520 |
| PS5-B3 | N | 98.1 | 98.1 | 98.6 | 98.6 | 96.6 | 96.7 | 99.4 | 99.4 |
| PS5-F1c | N | 99.1 | 99.2 | 99.4 | 99.6 | 96.6 | 96.7 | 98.9 | 99.9 |

(continued on next page)

(Table S35, Luo et al. [12] coverages, continued)

| | | SARS-CoV-2 | | Alpha | | Beta | | Gamma | |
|---|---|---|---|---|---|---|---|---|---|
| Primer | Gene Target | $C_{strict}$ | $C_{part.}$ | $C_{strict}$ | $C_{part.}$ | $C_{strict}$ | $C_{part.}$ | $C_{strict}$ | $C_{part.}$ |
| PS5-F2 | N | 99.1 | 99.1 | 96.9 | 97.2 | 97.6 | 97.6 | 99.7 | 99.7 |
| PS5-F3 | N | 99.1 | 99.1 | 99.7 | 99.7 | 97.2 | 97.2 | 99.6 | 99.6 |
| PS5-LB | N | 98.0 | 99.0 | 99.5 | 99.6 | 96.7 | 96.8 | 97.6 | 99.9 |
| PS5-LF | N | 98.8 | 99.1 | 99.4 | 99.6 | 96.4 | 96.7 | 99.5 | 99.8 |
| PS6-B1c | N | 99.2 | 99.4 | 99.4 | 99.7 | 98.9 | 99.3 | 98.7 | 99.1 |
| PS6-B2 | N | 99.5 | 99.5 | 99.9 | 99.9 | 99.3 | 99.3 | 99.7 | 99.7 |
| PS6-B3 | N | 98.4 | 98.4 | 99.4 | 99.5 | 97.1 | 97.1 | 99.6 | 99.6 |
| PS6-F1c | N | 99.0 | 99.2 | 99.8 | 99.9 | 98.4 | 99.2 | 98.9 | 99.1 |
| PS6-F2 | N | 98.4 | 98.4 | 99.0 | 99.0 | 92.6 | 92.6 | 98.7 | 98.7 |
| PS6-F3 | N | 99.3 | 99.3 | 98.7 | 98.7 | 98.3 | 98.3 | 98.8 | 98.8 |
| PS6-LB | N | 99.6 | 99.6 | 99.9 | 99.9 | 99.3 | 99.3 | 99.1 | 99.2 |
| PS6-LF | N | 99.3 | 99.4 | 99.8 | 99.8 | 99.2 | 99.2 | 99.2 | 99.2 |

Table S36: Matched and mismatched coverages for Luo et al. [12] set.

| | | Delta | | Lambda | | Mu | | Omicron | |
|---|---|---|---|---|---|---|---|---|---|
| Primer | Gene Target | $C_{\text{strict}}$ | $C_{\text{part.}}$ | $C_{\text{strict}}$ | $C_{\text{part.}}$ | $C_{\text{strict}}$ | $C_{\text{part.}}$ | $C_{\text{strict}}$ | $C_{\text{part.}}$ |
| PS1-B1c | N | 99.9 | 99.9 | 99.5 | 99.7 | 99.7 | 99.8 | 99.3 | 99.3 |
| PS1-B2 | N | 0.000 | 0.000 | 0.161 | 0.161 | 99.3 | 99.4 | 0.0854 | 0.0854 |
| PS1-B3 | N | 99.6 | 99.6 | 99.3 | 99.3 | 99.4 | 99.4 | 99.2 | 99.2 |
| PS1-F1c | N | 99.9 | 100. | 98.5 | 99.9 | 99.6 | 99.8 | 99.3 | 99.3 |
| PS1-F2 | N | 99.8 | 99.9 | 99.0 | 99.0 | 99.6 | 99.6 | 99.3 | 99.3 |
| PS1-F3 | N | 99.4 | 99.4 | 99.4 | 99.4 | 99.7 | 99.7 | 99.2 | 99.2 |
| PS1-LB | N | 99.8 | 100. | 70.9 | 99.5 | 98.5 | 99.8 | 99.1 | 99.2 |
| PS1-LF | N | 99.7 | 99.8 | 99.2 | 99.5 | 99.8 | 99.8 | 99.2 | 99.2 |
| PS2-B1c | N | 99.5 | 99.8 | 99.6 | 99.7 | 97.1 | 99.3 | 93.8 | 93.8 |
| PS2-B2 | N | 99.9 | 99.9 | 99.2 | 99.2 | 99.5 | 99.5 | 94.5 | 94.5 |
| PS2-B3 | N | 99.9 | 99.9 | 99.9 | 99.9 | 99.1 | 99.1 | 94.6 | 94.6 |
| PS2-F1c | N | 99.8 | 99.9 | 94.8 | 95.0 | 99.5 | 99.8 | 94.9 | 94.9 |
| PS2-F2 | N | 99.8 | 99.8 | 99.2 | 99.2 | 88.8 | 88.8 | 94.9 | 94.9 |
| PS2-F3 | N | 99.8 | 99.9 | 99.5 | 99.6 | 99.1 | 99.2 | 0.555 | 0.555 |
| PS2-LB | N | 99.9 | 100. | 99.5 | 99.6 | 97.1 | 97.2 | 93.8 | 93.8 |
| PS2-LF | N | 99.7 | 99.7 | 99.6 | 99.6 | 99.7 | 99.7 | 94.8 | 94.8 |
| PS3-B1c | N | 99.5 | 99.8 | 99.6 | 99.7 | 97.1 | 99.3 | 93.8 | 93.8 |
| PS3-B2 | N | 99.9 | 99.9 | 99.2 | 99.2 | 99.5 | 99.5 | 94.5 | 94.5 |
| PS3-B3 | N | 99.9 | 99.9 | 99.9 | 99.9 | 99.1 | 99.1 | 94.6 | 94.6 |
| PS3-F1c | N | 99.8 | 99.9 | 94.8 | 95.0 | 99.5 | 99.8 | 94.9 | 94.9 |
| PS3-F2 | N | 99.8 | 99.8 | 99.2 | 99.2 | 88.8 | 88.8 | 94.9 | 94.9 |
| PS3-F3 | N | 99.8 | 99.9 | 99.5 | 99.6 | 99.1 | 99.2 | 0.555 | 0.555 |
| PS3-LB | N | 99.9 | 99.9 | 99.5 | 99.5 | 97.1 | 97.1 | 93.8 | 93.8 |
| PS3-LF | N | 99.7 | 99.8 | 99.6 | 99.6 | 99.7 | 99.7 | 94.8 | 94.8 |
| PS4-B1c | N | 99.9 | 99.9 | 99.5 | 99.7 | 99.7 | 99.8 | 99.3 | 99.3 |
| PS4-B2 | N | 0.000 | 0.000 | 0.161 | 0.161 | 99.3 | 99.4 | 0.0854 | 0.0854 |
| PS4-B3 | N | 99.5 | 99.6 | 0.107 | 0.107 | 99.4 | 99.4 | 99.2 | 99.2 |
| PS4-F1c | N | 99.9 | 100. | 98.5 | 99.9 | 99.6 | 99.8 | 99.3 | 99.3 |
| PS4-F2 | N | 99.8 | 99.8 | 98.9 | 99.0 | 99.6 | 99.6 | 99.3 | 99.3 |
| PS4-F3 | N | 99.5 | 99.5 | 97.7 | 97.7 | 99.5 | 99.5 | 99.2 | 99.2 |
| PS4-LB | N | 99.8 | 100. | 71.1 | 99.5 | 99.3 | 99.8 | 99.1 | 99.2 |
| PS4-LF | N | 99.7 | 99.9 | 99.2 | 99.5 | 99.8 | 99.9 | 99.2 | 99.3 |
| PS5-B1c | N | 99.9 | 99.9 | 99.5 | 99.7 | 99.7 | 99.8 | 99.3 | 99.3 |
| PS5-B2 | N | 0.000 | 0.000 | 0.161 | 0.161 | 99.3 | 99.4 | 0.0854 | 0.0854 |
| PS5-B3 | N | 99.5 | 99.6 | 0.107 | 0.107 | 99.4 | 99.4 | 99.2 | 99.2 |
| PS5-F1c | N | 99.9 | 100. | 98.5 | 99.9 | 99.6 | 99.8 | 99.3 | 99.3 |

(continued on next page)

(Table S36, Luo et al. [12] set coverages, continued)

| | | Delta | | Lambda | | Mu | | Omicron | |
|---|---|---|---|---|---|---|---|---|---|
| Primer | Gene Target | $C_{strict}$ | $C_{part.}$ | $C_{strict}$ | $C_{part.}$ | $C_{strict}$ | $C_{part.}$ | $C_{strict}$ | $C_{part.}$ |
| PS5-F2 | N | 99.8 | 99.8 | 98.9 | 99.0 | 99.6 | 99.6 | 99.3 | 99.3 |
| PS5-F3 | N | 99.5 | 99.5 | 97.7 | 97.7 | 99.5 | 99.5 | 99.2 | 99.2 |
| PS5-LB | N | 99.8 | 100. | 71.1 | 99.5 | 99.3 | 99.8 | 99.1 | 99.2 |
| PS5-LF | N | 99.7 | 99.9 | 99.2 | 99.6 | 99.8 | 99.9 | 99.2 | 99.3 |
| PS6-B1c | N | 99.8 | 99.9 | 99.2 | 99.9 | 88.7 | 99.7 | 94.7 | 94.8 |
| PS6-B2 | N | 100. | 100. | 99.8 | 99.8 | 99.8 | 99.8 | 94.7 | 94.7 |
| PS6-B3 | N | 99.5 | 99.6 | 99.6 | 99.7 | 97.2 | 97.2 | 93.8 | 93.8 |
| PS6-F1c | N | 99.8 | 99.9 | 99.5 | 99.7 | 99.0 | 99.4 | 0.555 | 0.555 |
| PS6-F2 | N | 99.7 | 99.7 | 0.664 | 0.664 | 99.1 | 99.1 | 0.640 | 0.640 |
| PS6-F3 | N | 99.8 | 99.8 | 99.3 | 99.3 | 99.5 | 99.5 | 94.6 | 94.6 |
| PS6-LB | N | 99.8 | 99.8 | 99.6 | 99.7 | 99.6 | 99.6 | 94.8 | 94.8 |
| PS6-LF | N | 98.8 | 98.8 | 98.6 | 98.6 | 99.7 | 99.8 | 94.8 | 94.8 |

Table S37: Matched and mismatched coverages for Luo et al. [12] set.

| | | BA.2 | | BA.3 | | BA.4 | | BA.5 | |
|---|---|---|---|---|---|---|---|---|---|
| Primer | Gene Target | $C_{\text{strict}}$ | $C_{\text{part.}}$ | $C_{\text{strict}}$ | $C_{\text{part.}}$ | $C_{\text{strict}}$ | $C_{\text{part.}}$ | $C_{\text{strict}}$ | $C_{\text{part.}}$ |
| PS1-B1c | N | 99.9 | 99.9 | 99.7 | 99.7 | 98.9 | 98.9 | 98.9 | 98.9 |
| PS1-B2 | N | 0.0947 | 0.0947 | 0.000 | 0.000 | 0.000 | 0.000 | 0.000 | 0.000 |
| PS1-B3 | N | 99.7 | 99.7 | 100. | 100. | 100. | 100. | 99.8 | 99.8 |
| PS1-F1c | N | 100. | 100. | 99.7 | 99.7 | 99.7 | 100. | 99.7 | 99.8 |
| PS1-F2 | N | 99.9 | 99.9 | 99.7 | 99.7 | 99.7 | 99.7 | 99.6 | 99.7 |
| PS1-F3 | N | 99.2 | 99.2 | 99.7 | 99.7 | 0.954 | 0.954 | 97.4 | 97.4 |
| PS1-LB | N | 99.7 | 100. | 100. | 100. | 99.5 | 99.8 | 99.4 | 100. |
| PS1-LF | N | 99.7 | 99.7 | 99.4 | 99.4 | 100. | 100. | 99.8 | 99.9 |
| PS2-B1c | N | 99.6 | 99.8 | 94.3 | 94.3 | 99.0 | 99.0 | 98.7 | 98.7 |
| PS2-B2 | N | 98.4 | 98.4 | 95.7 | 95.7 | 99.2 | 99.2 | 98.6 | 98.6 |
| PS2-B3 | N | 100. | 100. | 95.7 | 95.7 | 99.7 | 99.7 | 98.5 | 98.5 |
| PS2-F1c | N | 98.1 | 98.1 | 89.4 | 89.9 | 99.2 | 99.4 | 96.7 | 98.5 |
| PS2-F2 | N | 99.6 | 99.6 | 96.0 | 96.0 | 99.7 | 99.7 | 98.9 | 98.9 |
| PS2-F3 | N | 1.31 | 1.31 | 0.862 | 0.862 | 0.318 | 0.318 | 0.975 | 0.975 |
| PS2-LB | N | 99.8 | 99.8 | 94.3 | 94.3 | 98.7 | 98.7 | 98.7 | 98.9 |
| PS2-LF | N | 99.9 | 99.9 | 96.0 | 96.0 | 99.5 | 99.5 | 99.4 | 99.4 |
| PS3-B1c | N | 99.6 | 99.8 | 94.3 | 94.3 | 99.0 | 99.0 | 98.7 | 98.7 |
| PS3-B2 | N | 98.4 | 98.4 | 95.7 | 95.7 | 99.2 | 99.2 | 98.6 | 98.6 |
| PS3-B3 | N | 100. | 100. | 95.7 | 95.7 | 99.7 | 99.7 | 98.5 | 98.5 |
| PS3-F1c | N | 98.1 | 98.1 | 89.4 | 89.9 | 99.2 | 99.4 | 96.7 | 98.5 |
| PS3-F2 | N | 99.6 | 99.6 | 96.0 | 96.0 | 99.7 | 99.7 | 98.9 | 98.9 |
| PS3-F3 | N | 1.31 | 1.31 | 0.862 | 0.862 | 0.318 | 0.318 | 0.975 | 0.975 |
| PS3-LB | N | 99.8 | 99.8 | 94.3 | 94.3 | 98.7 | 98.7 | 98.7 | 98.9 |
| PS3-LF | N | 99.9 | 99.9 | 96.0 | 96.0 | 99.5 | 99.5 | 99.3 | 99.4 |
| PS4-B1c | N | 99.9 | 99.9 | 99.7 | 99.7 | 98.9 | 98.9 | 98.9 | 98.9 |
| PS4-B2 | N | 0.0947 | 0.0947 | 0.000 | 0.000 | 0.000 | 0.000 | 0.000 | 0.000 |
| PS4-B3 | N | 99.6 | 99.7 | 100. | 100. | 100. | 100. | 99.9 | 99.9 |
| PS4-F1c | N | 100. | 100. | 99.7 | 99.7 | 99.7 | 100. | 99.7 | 99.8 |
| PS4-F2 | N | 99.9 | 99.9 | 99.7 | 99.7 | 99.7 | 99.7 | 99.5 | 99.6 |
| PS4-F3 | N | 99.9 | 99.9 | 99.4 | 99.4 | 98.9 | 98.9 | 99.4 | 99.4 |
| PS4-LB | N | 99.8 | 100. | 100. | 100. | 99.7 | 99.8 | 99.4 | 100. |
| PS4-LF | N | 99.7 | 99.7 | 99.4 | 99.4 | 100. | 100. | 99.8 | 100. |
| PS5-B1c | N | 99.9 | 99.9 | 99.7 | 99.7 | 98.9 | 98.9 | 98.9 | 98.9 |
| PS5-B2 | N | 0.0947 | 0.0947 | 0.000 | 0.000 | 0.000 | 0.000 | 0.000 | 0.000 |
| PS5-B3 | N | 99.6 | 99.7 | 100. | 100. | 100. | 100. | 99.9 | 99.9 |
| PS5-F1c | N | 100. | 100. | 99.7 | 99.7 | 99.7 | 100. | 99.7 | 99.8 |

(continued on next page)

(Table S37, Luo et al. [12] set coverages, continued)

| Primer | Gene Target | BA.2 | | BA.3 | | BA.4 | | BA.5 | |
|---|---|---|---|---|---|---|---|---|---|
| | | $C_{\text{strict}}$ | $C_{\text{part.}}$ | $C_{\text{strict}}$ | $C_{\text{part.}}$ | $C_{\text{strict}}$ | $C_{\text{part.}}$ | $C_{\text{strict}}$ | $C_{\text{part.}}$ |
| PS5-F2 | N | 99.9 | 99.9 | 99.7 | 99.7 | 99.7 | 99.7 | 99.5 | 99.6 |
| PS5-F3 | N | 99.9 | 99.9 | 99.4 | 99.4 | 98.9 | 98.9 | 99.4 | 99.4 |
| PS5-LB | N | 99.8 | 100. | 100. | 100. | 99.7 | 99.8 | 99.4 | 100. |
| PS5-LF | N | 99.7 | 99.9 | 99.4 | 99.4 | 100. | 100. | 99.8 | 100. |
| PS6-B1c | N | 99.5 | 99.9 | 96.0 | 96.0 | 99.5 | 99.8 | 98.6 | 98.9 |
| PS6-B2 | N | 100. | 100. | 96.0 | 96.0 | 99.7 | 99.7 | 99.1 | 99.1 |
| PS6-B3 | N | 99.6 | 99.6 | 94.3 | 94.3 | 99.0 | 99.0 | 98.7 | 98.7 |
| PS6-F1c | N | 1.31 | 1.31 | 0.862 | 0.862 | 0.318 | 0.318 | 0.975 | 1.14 |
| PS6-F2 | N | 0.000 | 0.000 | 1.15 | 1.15 | 0.477 | 0.477 | 0.569 | 0.569 |
| PS6-F3 | N | 99.5 | 99.5 | 94.3 | 94.3 | 100. | 100. | 99.3 | 99.3 |
| PS6-LB | N | 99.9 | 99.9 | 95.1 | 95.1 | 99.2 | 99.2 | 97.4 | 97.6 |
| PS6-LF | N | 99.9 | 99.9 | 95.4 | 95.4 | 99.0 | 99.0 | 89.4 | 89.4 |

Table S38: Matched and mismatched coverages for Mautner et al. [13] set.

| | | SARS-CoV-2 | | Alpha | | Beta | | Gamma | |
|---|---|---|---|---|---|---|---|---|---|
| Primer | Gene Target | $C_{\text{strict}}$ | $C_{\text{part.}}$ | $C_{\text{strict}}$ | $C_{\text{part.}}$ | $C_{\text{strict}}$ | $C_{\text{part.}}$ | $C_{\text{strict}}$ | $C_{\text{part.}}$ |
| OFR8-B2 | Orf8 | 98.3 | 98.3 | 68.6 | 68.6 | 97.9 | 97.9 | 90.9 | 90.9 |
| OFR8-F1c | Orf8 | 98.9 | 99.0 | 98.4 | 98.4 | 97.5 | 97.5 | 90.3 | 90.5 |
| OFR8-F2 | Orf8 | 99.1 | 99.1 | 98.5 | 98.5 | 98.3 | 98.3 | 90.1 | 90.1 |
| ORF8-B1c | Orf8 | 99.1 | 99.1 | 0.193 | 0.193 | 98.1 | 98.3 | 89.6 | 89.8 |
| ORF8-B3 | Orf8 | 99.3 | 99.3 | 99.8 | 99.8 | 99.1 | 99.1 | 2.21 | 2.21 |
| ORF8-F3 | Orf8 | 99.0 | 99.0 | 98.5 | 98.5 | 98.3 | 98.3 | 93.1 | 93.1 |
| ORF8-LB | Orf8 | 97.4 | 97.5 | 98.6 | 98.6 | 97.4 | 97.4 | 90.1 | 90.1 |
| ORF8-LF | Orf8 | 93.7 | 93.7 | 0.0552 | 0.0552 | 98.1 | 98.1 | 89.9 | 89.9 |

Table S39: Matched and mismatched coverages for Mautner et al. [13] set.

| Primer | Gene Target | Delta | | Lambda | | Mu | | Omicron | |
|---|---|---|---|---|---|---|---|---|---|
| | | $C_{\text{strict}}$ | $C_{\text{part.}}$ | $C_{\text{strict}}$ | $C_{\text{part.}}$ | $C_{\text{strict}}$ | $C_{\text{part.}}$ | $C_{\text{strict}}$ | $C_{\text{part.}}$ |
| OFR8-B2 | Orf8 | 99.0 | 99.0 | 98.0 | 98.0 | 3.77 | 3.77 | 84.2 | 84.2 |
| OFR8-F1c | Orf8 | 99.4 | 99.5 | 96.1 | 96.2 | 0.0908 | 0.0908 | 84.2 | 84.2 |
| OFR8-F2 | Orf8 | 99.6 | 99.6 | 98.1 | 98.1 | 99.4 | 99.4 | 83.5 | 83.5 |
| ORF8-B1c | Orf8 | 99.7 | 99.7 | 92.9 | 93.9 | 96.5 | 96.5 | 84.1 | 84.2 |
| ORF8-B3 | Orf8 | 99.9 | 99.9 | 98.8 | 98.8 | 99.2 | 99.2 | 91.9 | 91.9 |
| ORF8-F3 | Orf8 | 28.8 | 28.8 | 98.2 | 98.2 | 99.0 | 99.0 | 81.7 | 81.7 |
| ORF8-LB | Orf8 | 99.7 | 99.7 | 98.2 | 98.8 | 98.5 | 98.5 | 83.9 | 83.9 |
| ORF8-LF | Orf8 | 99.7 | 99.7 | 99.0 | 99.0 | 99.0 | 99.0 | 84.0 | 84.0 |

Table S40: Matched and mismatched coverages for Mautner et al. [13] set.

| | | BA.2 | | BA.3 | | BA.4 | | BA.5 | |
|---|---|---|---|---|---|---|---|---|---|
| Primer | Gene Target | $C_{\text{strict}}$ | $C_{\text{part.}}$ | $C_{\text{strict}}$ | $C_{\text{part.}}$ | $C_{\text{strict}}$ | $C_{\text{part.}}$ | $C_{\text{strict}}$ | $C_{\text{part.}}$ |
| OFR8-B2 | Orf8 | 99.4 | 99.4 | 89.7 | 89.7 | 98.3 | 98.3 | 96.3 | 96.3 |
| OFR8-F1c | Orf8 | 99.7 | 99.8 | 82.8 | 88.5 | 98.3 | 98.4 | 98.9 | 98.9 |
| OFR8-F2 | Orf8 | 99.6 | 99.6 | 88.2 | 88.2 | 98.4 | 98.4 | 98.3 | 98.3 |
| ORF8-B1c | Orf8 | 99.8 | 99.8 | 88.5 | 88.5 | 98.7 | 98.7 | 99.4 | 99.6 |
| ORF8-B3 | Orf8 | 99.7 | 99.7 | 95.4 | 95.4 | 99.8 | 99.8 | 96.3 | 96.3 |
| ORF8-F3 | Orf8 | 99.6 | 99.6 | 87.1 | 87.1 | 97.1 | 97.1 | 15.5 | 15.5 |
| ORF8-LB | Orf8 | 99.6 | 99.6 | 88.5 | 88.5 | 98.7 | 98.7 | 96.9 | 96.9 |
| ORF8-LF | Orf8 | 99.8 | 99.8 | 88.8 | 88.8 | 99.0 | 99.0 | 98.7 | 98.7 |

Table S41: Matched and mismatched coverages for Mohon et al. [14] set.

| Primer | Gene Target | SARS-CoV-2 | | Alpha | | Beta | | Gamma | |
|---|---|---|---|---|---|---|---|---|---|
| | | $C_{\text{strict}}$ | $C_{\text{part.}}$ | $C_{\text{strict}}$ | $C_{\text{part.}}$ | $C_{\text{strict}}$ | $C_{\text{part.}}$ | $C_{\text{strict}}$ | $C_{\text{part.}}$ |
| S2-B1c | S | 99.0 | 99.0 | 99.6 | 99.6 | 1.99 | 1.99 | 96.1 | 96.2 |
| S2-B2 | S | 97.6 | 97.6 | 96.7 | 96.7 | 55.1 | 55.1 | 93.4 | 93.4 |
| S2-B3 | S | 96.0 | 96.0 | 96.7 | 96.7 | 39.5 | 39.5 | 94.2 | 94.2 |
| S2-F1c | S | 99.1 | 99.2 | 99.4 | 99.7 | 97.8 | 97.8 | 96.4 | 96.4 |
| S2-F2 | S | 98.8 | 98.8 | 98.9 | 98.9 | 2.89 | 2.89 | 96.5 | 96.5 |
| S2-F3 | S | 99.1 | 99.1 | 99.7 | 99.7 | 98.2 | 98.2 | 96.7 | 96.7 |
| S2-LPB | S | 99.2 | 99.2 | 99.2 | 99.3 | 94.5 | 94.6 | 98.1 | 98.1 |
| S2-LPF | S | 98.2 | 98.2 | 99.4 | 99.4 | 97.3 | 97.3 | 96.2 | 96.2 |
| S3-B1c | RdRp | 99.2 | 99.3 | 99.7 | 99.7 | 98.9 | 99.0 | 99.4 | 99.4 |
| S3-B2 | RdRp | 99.3 | 99.3 | 99.8 | 99.8 | 99.1 | 99.1 | 99.5 | 99.5 |
| S3-B3 | RdRp | 99.4 | 99.4 | 99.3 | 99.3 | 99.0 | 99.0 | 99.5 | 99.5 |
| S3-F1c | RdRp | 98.8 | 98.9 | 99.6 | 99.6 | 98.9 | 99.0 | 99.4 | 99.4 |
| S3-F2 | RdRp | 99.5 | 99.5 | 99.8 | 99.8 | 99.1 | 99.1 | 99.5 | 99.5 |
| S3-F3 | RdRp | 99.4 | 99.4 | 0.110 | 0.110 | 99.0 | 99.0 | 99.5 | 99.5 |
| S3-LPB | RdRp | 99.1 | 99.2 | 99.8 | 99.9 | 99.1 | 99.1 | 99.5 | 99.5 |
| S3-LPF | RdRp | 97.1 | 97.1 | 99.6 | 99.6 | 99.0 | 99.0 | 99.5 | 99.5 |

Table S42: Matched and mismatched coverages for Mohon et al. [14] set.

| | | Delta | | Lambda | | Mu | | Omicron | |
|---|---|---|---|---|---|---|---|---|---|
| Primer | Gene Target | $C_{\text{strict}}$ | $C_{\text{part.}}$ | $C_{\text{strict}}$ | $C_{\text{part.}}$ | $C_{\text{strict}}$ | $C_{\text{part.}}$ | $C_{\text{strict}}$ | $C_{\text{part.}}$ |
| S2-B1c | S | 99.9 | 99.9 | 96.9 | 97.0 | 99.4 | 99.4 | 89.2 | 89.2 |
| S2-B2 | S | 99.2 | 99.2 | 91.3 | 91.3 | 91.7 | 91.7 | 94.8 | 94.8 |
| S2-B3 | S | 99.1 | 99.1 | 90.8 | 90.8 | 91.5 | 91.5 | 94.9 | 94.9 |
| S2-F1c | S | 99.9 | 99.9 | 98.2 | 98.2 | 99.5 | 99.6 | 89.4 | 89.4 |
| S2-F2 | S | 94.6 | 94.6 | 98.6 | 98.6 | 99.4 | 99.4 | 0.512 | 0.512 |
| S2-F3 | S | 99.6 | 99.6 | 99.2 | 99.2 | 99.3 | 99.3 | 90.4 | 90.4 |
| S2-LPB | S | 99.6 | 99.6 | 2.74 | 2.87 | 99.2 | 99.2 | 95.0 | 95.0 |
| S2-LPF | S | 85.3 | 85.3 | 97.5 | 97.5 | 99.4 | 99.4 | 89.8 | 89.8 |
| S3-B1c | RdRp | 99.9 | 99.9 | 99.7 | 99.7 | 99.9 | 99.9 | 90.3 | 90.3 |
| S3-B2 | RdRp | 99.9 | 99.9 | 100. | 100. | 100. | 100. | 91.0 | 91.0 |
| S3-B3 | RdRp | 2.19 | 2.19 | 99.7 | 99.7 | 99.5 | 99.5 | 90.9 | 90.9 |
| S3-F1c | RdRp | 100. | 100. | 99.9 | 99.9 | 99.8 | 99.8 | 90.3 | 90.3 |
| S3-F2 | RdRp | 100. | 100. | 99.9 | 99.9 | 100. | 100. | 99.6 | 99.6 |
| S3-F3 | RdRp | 99.5 | 99.5 | 99.9 | 99.9 | 99.8 | 99.8 | 99.6 | 99.6 |
| S3-LPB | RdRp | 100. | 100. | 100. | 100. | 98.1 | 98.2 | 90.0 | 90.0 |
| S3-LPF | RdRp | 99.9 | 99.9 | 99.6 | 99.6 | 90.0 | 90.0 | 90.6 | 90.6 |

Table S43: Matched and mismatched coverages for Mohon et al. [14] set.

| Primer | Gene Target | BA.2 | | BA.3 | | BA.4 | | BA.5 | |
|---|---|---|---|---|---|---|---|---|---|
| | | $C_{\text{strict}}$ | $C_{\text{part.}}$ | $C_{\text{strict}}$ | $C_{\text{part.}}$ | $C_{\text{strict}}$ | $C_{\text{part.}}$ | $C_{\text{strict}}$ | $C_{\text{part.}}$ |
| S2-B1c | S | 99.9 | 99.9 | 93.1 | 93.4 | 99.5 | 99.5 | 99.8 | 99.9 |
| S2-B2 | S | 99.9 | 99.9 | 96.8 | 96.8 | 99.0 | 99.0 | 96.7 | 96.7 |
| S2-B3 | S | 99.9 | 99.9 | 97.4 | 97.4 | 99.7 | 99.7 | 99.8 | 99.8 |
| S2-F1c | S | 100. | 100. | 92.8 | 92.8 | 99.8 | 99.8 | 99.9 | 99.9 |
| S2-F2 | S | 0.000 | 0.000 | 0.000 | 0.000 | 0.318 | 0.318 | 0.325 | 0.325 |
| S2-F3 | S | 99.9 | 99.9 | 93.4 | 93.4 | 99.8 | 99.8 | 99.8 | 99.8 |
| S2-LPB | S | 99.9 | 99.9 | 97.4 | 97.4 | 99.4 | 99.4 | 98.9 | 99.0 |
| S2-LPF | S | 99.9 | 99.9 | 92.8 | 92.8 | 99.4 | 99.4 | 99.6 | 99.6 |
| S3-B1c | RdRp | 99.8 | 99.8 | 98.6 | 98.6 | 99.7 | 99.7 | 95.0 | 95.0 |
| S3-B2 | RdRp | 99.9 | 99.9 | 99.1 | 99.1 | 99.7 | 99.7 | 99.5 | 99.5 |
| S3-B3 | RdRp | 99.9 | 99.9 | 99.1 | 99.1 | 99.0 | 99.0 | 99.4 | 99.4 |
| S3-F1c | RdRp | 100. | 100. | 98.9 | 98.9 | 99.4 | 99.4 | 99.8 | 99.8 |
| S3-F2 | RdRp | 100. | 100. | 100. | 100. | 100. | 100. | 100. | 100. |
| S3-F3 | RdRp | 99.8 | 99.8 | 99.1 | 99.1 | 100. | 100. | 100. | 100. |
| S3-LPB | RdRp | 100. | 100. | 98.9 | 98.9 | 99.0 | 99.2 | 99.0 | 99.0 |
| S3-LPF | RdRp | 100. | 100. | 99.1 | 99.1 | 99.4 | 99.4 | 94.9 | 94.9 |

Table S44: Matched and mismatched coverages for Rodriguez-Manzano et al. [15] set.

| Primer | Gene Target | SARS-CoV-2 | | Alpha | | Beta | | Gamma | |
|---|---|---|---|---|---|---|---|---|---|
| | | $C_{\text{strict}}$ | $C_{\text{part.}}$ | $C_{\text{strict}}$ | $C_{\text{part.}}$ | $C_{\text{strict}}$ | $C_{\text{part.}}$ | $C_{\text{strict}}$ | $C_{\text{part.}}$ |
| B1c-N | N | 99.3 | 99.4 | 99.8 | 99.9 | 97.5 | 98.1 | 99.5 | 99.6 |
| B2-N | N | 99.1 | 99.1 | 99.7 | 99.7 | 97.2 | 97.2 | 99.6 | 99.6 |
| B3-N | N | 99.0 | 99.0 | 96.7 | 96.8 | 97.5 | 97.5 | 99.7 | 99.7 |
| F1c-N | N | 99.4 | 99.5 | 99.8 | 99.9 | 98.0 | 98.1 | 98.1 | 98.2 |
| F2-N | N | 0.000 | 0.000 | 0.000 | 0.000 | 0.000 | 0.000 | 0.000 | 0.000 |
| F3-N | N | 99.4 | 99.4 | 99.8 | 99.8 | 98.0 | 98.0 | 0.780 | 0.780 |
| LB-N | N | 98.9 | 99.2 | 97.9 | 97.9 | 97.3 | 97.3 | 99.5 | 99.5 |
| LF-N | N | 99.4 | 99.5 | 99.8 | 99.9 | 98.2 | 98.3 | 99.6 | 99.6 |

Table S45: Matched and mismatched coverages for Rodriguez-Manzano et al. [15] set.

| | | Delta | | Lambda | | Mu | | Omicron | |
|---|---|---|---|---|---|---|---|---|---|
| Primer | Gene Target | $C_{\text{strict}}$ | $C_{\text{part.}}$ | $C_{\text{strict}}$ | $C_{\text{part.}}$ | $C_{\text{strict}}$ | $C_{\text{part.}}$ | $C_{\text{strict}}$ | $C_{\text{part.}}$ |
| B1c-N | N | 99.9 | 100. | 99.6 | 99.8 | 99.1 | 99.8 | 94.2 | 94.3 |
| B2-N | N | 99.5 | 99.5 | 97.7 | 97.7 | 99.5 | 99.5 | 99.2 | 99.2 |
| B3-N | N | 99.8 | 99.8 | 99.1 | 99.2 | 99.5 | 99.6 | 99.2 | 99.2 |
| F1c-N | N | 99.9 | 99.9 | 99.6 | 100. | 85.2 | 99.7 | 94.2 | 94.4 |
| F2-N | N | 0.000 | 0.000 | 0.000 | 0.000 | 0.000 | 0.000 | 0.000 | 0.000 |
| F3-N | N | 99.9 | 99.9 | 99.6 | 99.6 | 97.0 | 97.0 | 94.1 | 94.1 |
| LB-N | N | 99.4 | 99.5 | 99.4 | 99.5 | 99.6 | 99.6 | 94.8 | 94.8 |
| LF-N | N | 99.9 | 99.9 | 99.8 | 99.9 | 99.1 | 99.6 | 94.4 | 94.5 |

Table S46: Matched and mismatched coverages for Rodriguez-Manzano et al. [15] set.

| Primer | Gene Target | BA.2 | | BA.3 | | BA.4 | | BA.5 | |
|---|---|---|---|---|---|---|---|---|---|
| | | $C_{\text{strict}}$ | $C_{\text{part.}}$ | $C_{\text{strict}}$ | $C_{\text{part.}}$ | $C_{\text{strict}}$ | $C_{\text{part.}}$ | $C_{\text{strict}}$ | $C_{\text{part.}}$ |
| B1c-N | N | 99.9 | 100. | 96.3 | 96.3 | 100. | 100. | 99.3 | 99.5 |
| B2-N | N | 99.9 | 99.9 | 99.4 | 99.4 | 98.9 | 98.9 | 99.4 | 99.4 |
| B3-N | N | 99.9 | 99.9 | 99.7 | 99.7 | 100. | 100. | 99.5 | 99.5 |
| F1c-N | N | 100. | 100. | 95.7 | 95.7 | 100. | 100. | 99.6 | 99.7 |
| F2-N | N | 0.000 | 0.000 | 0.000 | 0.000 | 0.000 | 0.000 | 0.000 | 0.000 |
| F3-N | N | 99.8 | 99.8 | 94.3 | 94.3 | 99.5 | 99.5 | 98.9 | 98.9 |
| LB-N | N | 99.6 | 99.6 | 97.7 | 97.7 | 99.8 | 99.8 | 91.8 | 92.0 |
| LF-N | N | 99.8 | 100. | 95.7 | 95.7 | 99.8 | 100. | 98.2 | 99.5 |

Table S47: Matched and mismatched coverages for Yan et al. [16] set.

| Primer | Gene Target | SARS-CoV-2 | | Alpha | | Beta | | Gamma | |
|---|---|---|---|---|---|---|---|---|---|
| | | $C_{\text{strict}}$ | $C_{\text{part.}}$ | $C_{\text{strict}}$ | $C_{\text{part.}}$ | $C_{\text{strict}}$ | $C_{\text{part.}}$ | $C_{\text{strict}}$ | $C_{\text{part.}}$ |
| S-123B1c | S | 99.3 | 99.4 | 99.4 | 99.4 | 97.5 | 97.5 | 99.0 | 99.0 |
| S-123B2 | S | 99.0 | 99.0 | 99.4 | 99.4 | 96.6 | 96.6 | 99.0 | 99.0 |
| S-123B3 | S | 98.9 | 98.9 | 99.3 | 99.3 | 94.0 | 94.0 | 99.0 | 99.0 |
| S-123F1c | S | 99.3 | 99.3 | 99.5 | 99.5 | 98.9 | 98.9 | 99.3 | 99.3 |
| S-123F2 | S | 98.6 | 98.6 | 99.5 | 99.5 | 97.8 | 97.8 | 99.1 | 99.1 |
| S-123F3 | S | 98.7 | 98.7 | 0.138 | 0.138 | 98.8 | 98.8 | 99.1 | 99.1 |
| S-123LB | S | 99.2 | 99.3 | 99.3 | 99.4 | 96.2 | 96.5 | 99.0 | 99.0 |
| S-123LF | S | 99.1 | 99.2 | 99.1 | 99.1 | 97.3 | 97.8 | 99.2 | 99.2 |
| orf1ab-4B1c | Orf1ab | 99.4 | 99.5 | 99.9 | 99.9 | 99.0 | 99.5 | 99.8 | 99.9 |
| orf1ab-4B2 | Orf1ab | 99.5 | 99.5 | 99.8 | 99.8 | 99.5 | 99.5 | 99.9 | 99.9 |
| orf1ab-4B3 | Orf1ab | 99.4 | 99.4 | 99.8 | 99.8 | 99.6 | 99.6 | 100. | 100. |
| orf1ab-4F1c | Orf1ab | 99.4 | 99.4 | 99.9 | 99.9 | 99.5 | 99.6 | 99.9 | 99.9 |
| orf1ab-4F2 | Orf1ab | 99.5 | 99.5 | 99.8 | 99.8 | 99.1 | 99.1 | 100. | 100. |
| orf1ab-4F3 | Orf1ab | 99.3 | 99.3 | 99.7 | 99.7 | 99.6 | 99.6 | 99.8 | 99.8 |
| orf1ab-4LF | Orf1ab | 99.2 | 99.2 | 99.4 | 99.4 | 99.5 | 99.5 | 99.8 | 100. |

Table S48: Matched and mismatched coverages for Yan et al. [16] set.

| | | Delta | | Lambda | | Mu | | Omicron | |
|---|---|---|---|---|---|---|---|---|---|
| Primer | Gene Target | $C_{\text{strict}}$ | $C_{\text{part.}}$ | $C_{\text{strict}}$ | $C_{\text{part.}}$ | $C_{\text{strict}}$ | $C_{\text{part.}}$ | $C_{\text{strict}}$ | $C_{\text{part.}}$ |
| S-123B1c | S | 100. | 100. | 99.0 | 99.2 | 99.5 | 99.6 | 75.9 | 75.9 |
| S-123B2 | S | 100. | 100. | 99.2 | 99.2 | 98.8 | 98.8 | 98.5 | 98.5 |
| S-123B3 | S | 99.8 | 99.8 | 98.5 | 98.5 | 99.5 | 99.5 | 98.4 | 98.4 |
| S-123F1c | S | 99.9 | 100. | 98.6 | 98.6 | 98.6 | 99.8 | 74.5 | 74.5 |
| S-123F2 | S | 97.9 | 97.9 | 0.0642 | 0.0642 | 100. | 100. | 87.2 | 87.2 |
| S-123F3 | S | 100. | 100. | 95.1 | 95.1 | 99.8 | 99.8 | 88.0 | 88.0 |
| S-123LB | S | 99.9 | 100. | 99.1 | 99.3 | 99.0 | 99.2 | 0.726 | 0.726 |
| S-123LF | S | 99.9 | 99.9 | 99.5 | 99.6 | 99.7 | 99.8 | 86.4 | 86.4 |
| orf1ab-4B1c | Orf1ab | 99.9 | 99.9 | 99.9 | 100. | 99.8 | 100. | 99.8 | 99.9 |
| orf1ab-4B2 | Orf1ab | 100. | 100. | 99.8 | 99.8 | 99.9 | 99.9 | 99.5 | 99.5 |
| orf1ab-4B3 | Orf1ab | 99.8 | 99.8 | 99.6 | 99.6 | 99.5 | 99.5 | 99.9 | 99.9 |
| orf1ab-4F1c | Orf1ab | 99.3 | 100. | 99.7 | 99.7 | 99.9 | 100. | 99.9 | 99.9 |
| orf1ab-4F2 | Orf1ab | 99.8 | 99.8 | 99.6 | 99.6 | 99.6 | 99.6 | 99.9 | 99.9 |
| orf1ab-4F3 | Orf1ab | 100. | 100. | 99.9 | 99.9 | 99.7 | 99.7 | 99.7 | 99.7 |
| orf1ab-4LF | Orf1ab | 99.8 | 100. | 99.7 | 100. | 99.8 | 100. | 99.9 | 99.9 |

Table S49: Matched and mismatched coverages for Yan et al. [16] set.

| Primer | Gene Target | BA.2 | | BA.3 | | BA.4 | | BA.5 | |
|---|---|---|---|---|---|---|---|---|---|
| | | $C_{\text{strict}}$ | $C_{\text{part.}}$ | $C_{\text{strict}}$ | $C_{\text{part.}}$ | $C_{\text{strict}}$ | $C_{\text{part.}}$ | $C_{\text{strict}}$ | $C_{\text{part.}}$ |
| S-123B1c | S | 100. | 100. | 94.8 | 94.8 | 100. | 100. | 99.9 | 99.9 |
| S-123B2 | S | 99.8 | 99.8 | 96.3 | 96.3 | 99.5 | 99.5 | 99.6 | 99.6 |
| S-123B3 | S | 100. | 100. | 96.6 | 96.6 | 99.4 | 99.4 | 99.6 | 99.6 |
| S-123F1c | S | 99.8 | 100. | 93.7 | 93.7 | 99.8 | 99.8 | 99.8 | 99.9 |
| S-123F2 | S | 99.9 | 99.9 | 95.1 | 95.1 | 100. | 100. | 99.8 | 99.8 |
| S-123F3 | S | 99.9 | 99.9 | 93.7 | 93.7 | 99.8 | 99.8 | 99.8 | 99.8 |
| S-123LB | S | 0.0135 | 0.0135 | 0.000 | 0.000 | 0.636 | 0.795 | 0.569 | 0.569 |
| S-123LF | S | 99.2 | 99.9 | 94.0 | 95.1 | 99.8 | 99.8 | 99.8 | 99.8 |
| orf1ab-4B1c | Orf1ab | 100. | 100. | 100. | 100. | 100. | 100. | 99.9 | 100. |
| orf1ab-4B2 | Orf1ab | 100. | 100. | 100. | 100. | 99.4 | 99.4 | 99.7 | 99.7 |
| orf1ab-4B3 | Orf1ab | 100. | 100. | 100. | 100. | 99.4 | 99.4 | 100. | 100. |
| orf1ab-4F1c | Orf1ab | 99.9 | 99.9 | 100. | 100. | 100. | 100. | 100. | 100. |
| orf1ab-4F2 | Orf1ab | 99.9 | 99.9 | 99.7 | 99.7 | 99.7 | 99.7 | 99.8 | 99.8 |
| orf1ab-4F3 | Orf1ab | 100. | 100. | 99.4 | 99.4 | 100. | 100. | 99.4 | 99.4 |
| orf1ab-4LF | Orf1ab | 99.9 | 100. | 99.7 | 99.7 | 99.8 | 99.8 | 99.8 | 100. |

Table S50: Matched and mismatched coverages for Yang et al. [17] set.

| Primer | Gene Target | SARS-CoV-2 | | Alpha | | Beta | | Gamma | |
|---|---|---|---|---|---|---|---|---|---|
| | | $C_{\text{strict}}$ | $C_{\text{part.}}$ | $C_{\text{strict}}$ | $C_{\text{part.}}$ | $C_{\text{strict}}$ | $C_{\text{part.}}$ | $C_{\text{strict}}$ | $C_{\text{part.}}$ |
| CU-N2-B1c | N | 97.9 | 98.4 | 98.6 | 98.9 | 96.6 | 96.8 | 99.4 | 99.7 |
| CU-N2-B2 | N | 98.7 | 99.1 | 99.4 | 99.5 | 96.5 | 96.6 | 99.5 | 99.5 |
| CU-N2-B3 | N | 98.1 | 98.1 | 98.4 | 98.4 | 96.5 | 96.5 | 99.7 | 99.7 |
| CU-N2-F1c | N | 65.2 | 66.6 | 0.0690 | 0.0690 | 96.3 | 96.4 | 0.477 | 0.563 |
| CU-N2-F2 | N | 98.2 | 98.3 | 99.5 | 99.5 | 96.8 | 96.8 | 99.8 | 99.8 |
| CU-N2-F3 | N | 97.9 | 97.9 | 99.1 | 99.1 | 96.6 | 96.6 | 98.5 | 98.5 |
| CU-N2-Loop-B | N | 98.9 | 99.1 | 99.4 | 99.5 | 96.7 | 96.7 | 99.8 | 99.8 |
| ORF1e-B1c | Orf1ab | 0.000 | 99.2 | 0.000 | 98.9 | 0.000 | 98.9 | 0.000 | 100. |
| ORF1e-B2 | Orf1ab | 98.8 | 98.8 | 99.1 | 99.1 | 98.7 | 98.7 | 99.0 | 99.0 |
| ORF1e-B3 | Orf1ab | 99.3 | 99.3 | 99.1 | 99.1 | 98.8 | 98.8 | 99.7 | 99.7 |
| ORF1e-F1c | Orf1ab | 99.2 | 99.3 | 99.6 | 99.6 | 96.4 | 97.3 | 98.6 | 98.7 |
| ORF1e-F2 | Orf1ab | 99.2 | 99.2 | 99.7 | 99.7 | 98.0 | 98.0 | 98.8 | 98.8 |
| ORF1e-F3 | Orf1ab | 98.7 | 98.7 | 88.9 | 88.9 | 97.8 | 97.8 | 98.4 | 98.4 |
| ORF1e-Loop-B | Orf1ab | 99.0 | 99.0 | 99.4 | 99.4 | 98.2 | 98.2 | 99.8 | 99.8 |
| ORF1e-Loop-F | Orf1ab | 98.7 | 98.7 | 99.5 | 99.5 | 97.6 | 97.6 | 98.6 | 98.6 |

Table S51: Matched and mismatched coverages for Yang et al. [17] set.

| Primer | Gene Target | Delta | | Lambda | | Mu | | Omicron | |
|---|---|---|---|---|---|---|---|---|---|
| | | $C_{\text{strict}}$ | $C_{\text{part.}}$ | $C_{\text{strict}}$ | $C_{\text{part.}}$ | $C_{\text{strict}}$ | $C_{\text{part.}}$ | $C_{\text{strict}}$ | $C_{\text{part.}}$ |
| CU-N2-B1c | N | 99.6 | 99.9 | 99.3 | 99.5 | 99.4 | 99.8 | 99.2 | 99.2 |
| CU-N2-B2 | N | 99.9 | 99.9 | 99.6 | 99.7 | 99.8 | 99.8 | 99.1 | 99.1 |
| CU-N2-B3 | N | 99.8 | 99.8 | 99.4 | 99.4 | 99.5 | 99.5 | 99.0 | 99.0 |
| CU-N2-F1c | N | 0.000 | 0.000 | 0.161 | 0.161 | 99.3 | 99.6 | 0.0854 | 0.0854 |
| CU-N2-F2 | N | 99.8 | 99.8 | 99.4 | 99.4 | 99.4 | 99.4 | 99.2 | 99.2 |
| CU-N2-F3 | N | 99.9 | 99.9 | 99.6 | 99.6 | 99.7 | 99.7 | 99.3 | 99.3 |
| CU-N2-Loop-B | N | 28.9 | 28.9 | 0.107 | 0.107 | 99.6 | 99.6 | 98.8 | 98.8 |
| ORF1e-B1c | Orf1ab | 0.000 | 100. | 0.000 | 99.7 | 0.000 | 99.6 | 0.000 | 99.0 |
| ORF1e-B2 | Orf1ab | 99.8 | 99.8 | 99.6 | 99.6 | 99.8 | 99.8 | 98.5 | 98.5 |
| ORF1e-B3 | Orf1ab | 99.2 | 99.2 | 99.0 | 99.0 | 99.7 | 99.7 | 98.6 | 98.6 |
| ORF1e-F1c | Orf1ab | 99.9 | 99.9 | 99.5 | 99.8 | 99.6 | 99.8 | 99.6 | 99.7 |
| ORF1e-F2 | Orf1ab | 99.9 | 99.9 | 99.7 | 99.7 | 97.9 | 97.9 | 99.7 | 99.7 |
| ORF1e-F3 | Orf1ab | 99.8 | 99.8 | 98.2 | 98.2 | 99.6 | 99.6 | 99.7 | 99.7 |
| ORF1e-Loop-B | Orf1ab | 99.9 | 99.9 | 99.5 | 99.5 | 99.8 | 99.8 | 98.4 | 98.4 |
| ORF1e-Loop-F | Orf1ab | 99.9 | 99.9 | 99.5 | 99.5 | 99.0 | 99.0 | 99.6 | 99.6 |

Table S52: Matched and mismatched coverages for Yang et al. [17] set.

| Primer | Gene Target | BA.2 | | BA.3 | | BA.4 | | BA.5 | |
|---|---|---|---|---|---|---|---|---|---|
| | | $C_{\text{strict}}$ | $C_{\text{part.}}$ | $C_{\text{strict}}$ | $C_{\text{part.}}$ | | | | |
| CU-N2-B1c | N | 99.6 | 99.7 | 100. | 100. | 100. | 100. | 99.8 | 99.9 |
| CU-N2-B2 | N | 99.9 | 99.9 | 100. | 100. | 99.5 | 99.5 | 99.8 | 99.8 |
| CU-N2-B3 | N | 99.1 | 99.1 | 98.9 | 98.9 | 99.0 | 99.0 | 99.5 | 99.5 |
| CU-N2-F1c | N | 0.0947 | 0.0947 | 0.000 | 0.000 | 0.000 | 0.000 | 0.000 | 0.000 |
| CU-N2-F2 | N | 100. | 100. | 100. | 100. | 100. | 100. | 99.4 | 99.4 |
| CU-N2-F3 | N | 99.9 | 99.9 | 99.7 | 99.7 | 99.0 | 99.0 | 98.9 | 98.9 |
| CU-N2-Loop-B | N | 99.8 | 99.9 | 98.9 | 98.9 | 100. | 100. | 99.6 | 99.6 |
| ORF1e-B1c | Orf1ab | 0.000 | 100. | 0.000 | 98.6 | 0.000 | 99.8 | 0.000 | 96.0 |
| ORF1e-B2 | Orf1ab | 100. | 100. | 98.9 | 98.9 | 99.7 | 99.7 | 99.5 | 99.5 |
| ORF1e-B3 | Orf1ab | 100. | 100. | 98.6 | 98.6 | 99.5 | 99.5 | 99.5 | 99.5 |
| ORF1e-F1c | Orf1ab | 99.7 | 100. | 98.9 | 99.1 | 99.7 | 100. | 99.2 | 99.8 |
| ORF1e-F2 | Orf1ab | 99.9 | 99.9 | 99.1 | 99.1 | 99.0 | 99.0 | 99.7 | 99.7 |
| ORF1e-F3 | Orf1ab | 99.8 | 99.8 | 99.7 | 99.7 | 99.0 | 99.0 | 99.7 | 99.7 |
| ORF1e-Loop-B | Orf1ab | 99.7 | 99.7 | 98.6 | 98.6 | 99.5 | 99.5 | 99.4 | 99.4 |
| ORF1e-Loop-F | Orf1ab | 99.8 | 99.8 | 99.1 | 99.1 | 99.0 | 99.0 | 99.5 | 99.5 |

Table S53: Matched and mismatched coverages for Yoshikawa et al. [18] set.

| Primer | Gene Target | SARS-CoV-2 | | Alpha | | Beta | | Gamma | |
|---|---|---|---|---|---|---|---|---|---|
| | | $C_{\text{strict}}$ | $C_{\text{part.}}$ | $C_{\text{strict}}$ | $C_{\text{part.}}$ | $C_{\text{strict}}$ | $C_{\text{part.}}$ | $C_{\text{strict}}$ | $C_{\text{part.}}$ |
| LAMP_ORF1b-1_B1c | Orf1b | 98.9 | 98.9 | 99.4 | 99.4 | 98.7 | 98.7 | 98.7 | 98.7 |
| LAMP_ORF1b-1_B2 | Orf1b | 98.9 | 98.9 | 98.5 | 98.5 | 98.5 | 98.5 | 99.0 | 99.0 |
| LAMP_ORF1b-1_B3 | Orf1b | 99.0 | 99.0 | 99.5 | 99.5 | 98.7 | 98.7 | 98.9 | 98.9 |
| LAMP_ORF1b-1_F1c | Orf1b | 98.6 | 99.1 | 99.2 | 99.5 | 98.8 | 98.9 | 98.9 | 98.9 |
| LAMP_ORF1b-1_F2 | Orf1b | 99.0 | 99.0 | 99.5 | 99.5 | 98.7 | 98.7 | 98.9 | 98.9 |
| LAMP_ORF1b-1_F3 | Orf1b | 99.5 | 99.5 | 99.4 | 99.4 | 98.7 | 98.7 | 99.1 | 99.1 |
| LAMP_ORF1b-1_LB | Orf1b | 98.2 | 98.5 | 99.3 | 99.5 | 98.4 | 98.4 | 98.3 | 98.8 |
| LAMP_ORF1b-1_LF | Orf1b | 98.8 | 98.8 | 99.0 | 99.0 | 98.3 | 98.3 | 97.5 | 97.5 |
| LAMP_ORF1b-2_B1c | Orf1b | 97.9 | 97.9 | 96.8 | 96.8 | 91.3 | 91.6 | 94.1 | 94.1 |
| LAMP_ORF1b-2_B2 | Orf1b | 0.000 | 0.000 | 0.000 | 0.000 | 0.000 | 0.000 | 0.000 | 0.000 |
| LAMP_ORF1b-2_B3 | Orf1b | 98.7 | 98.7 | 97.2 | 97.2 | 91.4 | 91.4 | 93.8 | 93.8 |
| LAMP_ORF1b-2_F1c | Orf1b | 98.5 | 98.5 | 97.3 | 97.3 | 91.6 | 91.6 | 94.0 | 94.0 |
| LAMP_ORF1b-2_F2 | Orf1b | 0.000 | 98.8 | 0.000 | 98.2 | 0.000 | 94.1 | 0.000 | 95.8 |
| LAMP_ORF1b-2_F3 | Orf1b | 99.2 | 99.2 | 99.5 | 99.5 | 96.7 | 96.7 | 98.6 | 98.6 |
| LAMP_ORF1b-2_LB | Orf1b | 98.7 | 98.7 | 96.8 | 96.9 | 91.3 | 91.4 | 93.9 | 93.9 |
| LAMP_ORF1b-2_LF | Orf1b | 98.5 | 98.6 | 96.2 | 97.3 | 87.7 | 87.7 | 94.0 | 94.0 |

Table S54: Matched and mismatched coverages for Yoshikawa et al. [18] set.

| | | Delta | | Lambda | | Mu | | Omicron | |
|---|---|---|---|---|---|---|---|---|---|
| Primer | Gene Target | $C_{\text{strict}}$ | $C_{\text{part.}}$ | $C_{\text{strict}}$ | $C_{\text{part.}}$ | $C_{\text{strict}}$ | $C_{\text{part.}}$ | $C_{\text{strict}}$ | $C_{\text{part.}}$ |
| LAMP_ORF1b-1_B1c | Orf1b | 99.8 | 99.8 | 99.3 | 99.3 | 99.1 | 99.3 | 98.9 | 98.9 |
| LAMP_ORF1b-1_B2 | Orf1b | 99.7 | 99.7 | 99.7 | 99.7 | 99.3 | 99.3 | 98.8 | 98.8 |
| LAMP_ORF1b-1_B3 | Orf1b | 99.9 | 99.9 | 98.1 | 98.1 | 99.8 | 99.8 | 98.7 | 98.7 |
| LAMP_ORF1b-1_F1c | Orf1b | 99.7 | 99.7 | 99.3 | 99.8 | 99.5 | 99.8 | 98.6 | 98.8 |
| LAMP_ORF1b-1_F2 | Orf1b | 99.9 | 99.9 | 99.5 | 99.6 | 99.7 | 99.7 | 98.8 | 98.8 |
| LAMP_ORF1b-1_F3 | Orf1b | 99.8 | 99.8 | 99.8 | 99.8 | 99.8 | 99.8 | 98.9 | 98.9 |
| LAMP_ORF1b-1_LB | Orf1b | 99.7 | 99.8 | 99.6 | 99.6 | 99.3 | 99.4 | 98.7 | 98.8 |
| LAMP_ORF1b-1_LF | Orf1b | 99.6 | 99.6 | 99.8 | 99.8 | 99.5 | 99.5 | 98.4 | 98.4 |
| LAMP_ORF1b-2_B1c | Orf1b | 99.5 | 99.7 | 95.3 | 95.4 | 96.4 | 96.4 | 98.7 | 98.7 |
| LAMP_ORF1b-2_B2 | Orf1b | 0.000 | 0.000 | 0.000 | 0.000 | 0.000 | 0.000 | 0.000 | 0.000 |
| LAMP_ORF1b-2_B3 | Orf1b | 99.9 | 99.9 | 93.8 | 93.8 | 99.2 | 99.2 | 98.7 | 98.7 |
| LAMP_ORF1b-2_F1c | Orf1b | 99.8 | 99.8 | 95.1 | 95.1 | 98.9 | 98.9 | 98.5 | 98.5 |
| LAMP_ORF1b-2_F2 | Orf1b | 0.000 | 99.1 | 0.000 | 96.1 | 0.000 | 94.7 | 0.000 | 98.8 |
| LAMP_ORF1b-2_F3 | Orf1b | 99.9 | 99.9 | 99.8 | 99.8 | 99.0 | 99.0 | 99.0 | 99.0 |
| LAMP_ORF1b-2_LB | Orf1b | 99.9 | 99.9 | 95.1 | 95.1 | 98.6 | 99.0 | 98.5 | 98.6 |
| LAMP_ORF1b-2_LF | Orf1b | 99.9 | 99.9 | 95.6 | 95.7 | 99.2 | 99.2 | 98.4 | 98.5 |

Table S55: Matched and mismatched coverages for Yoshikawa et al. [18] set.

| Primer | Gene Target | BA.2 | | BA.3 | | BA.4 | | BA.5 | |
|---|---|---|---|---|---|---|---|---|---|
| | | $C_{\text{strict}}$ | $C_{\text{part.}}$ | $C_{\text{strict}}$ | $C_{\text{part.}}$ | $C_{\text{strict}}$ | $C_{\text{part.}}$ | $C_{\text{strict}}$ | $C_{\text{part.}}$ |
| LAMP_ORF1b-1_B1c | Orf1b | 97.9 | 99.7 | 99.1 | 99.1 | 99.7 | 99.7 | 99.6 | 99.6 |
| LAMP_ORF1b-1_B2 | Orf1b | 99.8 | 99.8 | 98.9 | 98.9 | 99.4 | 99.4 | 99.9 | 99.9 |
| LAMP_ORF1b-1_B3 | Orf1b | 100. | 100. | 98.9 | 98.9 | 99.7 | 99.7 | 99.4 | 99.4 |
| LAMP_ORF1b-1_F1c | Orf1b | 100. | 100. | 98.9 | 98.9 | 99.7 | 99.7 | 98.8 | 99.4 |
| LAMP_ORF1b-1_F2 | Orf1b | 100. | 100. | 99.1 | 99.1 | 99.7 | 99.7 | 99.9 | 99.9 |
| LAMP_ORF1b-1_F3 | Orf1b | 100. | 100. | 99.1 | 99.1 | 99.7 | 99.7 | 100. | 100. |
| LAMP_ORF1b-1_LB | Orf1b | 99.9 | 100. | 98.9 | 98.9 | 99.4 | 99.5 | 99.6 | 99.8 |
| LAMP_ORF1b-1_LF | Orf1b | 100. | 100. | 99.1 | 99.1 | 99.7 | 99.7 | 100. | 100. |
| LAMP_ORF1b-2_B1c | Orf1b | 100. | 100. | 98.6 | 98.6 | 99.5 | 99.5 | 99.8 | 99.8 |
| LAMP_ORF1b-2_B2 | Orf1b | 0.000 | 0.000 | 0.000 | 0.000 | 0.000 | 0.000 | 0.000 | 0.000 |
| LAMP_ORF1b-2_B3 | Orf1b | 100. | 100. | 97.1 | 97.1 | 99.7 | 99.7 | 100. | 100. |
| LAMP_ORF1b-2_F1c | Orf1b | 100. | 100. | 98.6 | 98.6 | 100. | 100. | 98.0 | 98.0 |
| LAMP_ORF1b-2_F2 | Orf1b | 0.000 | 99.5 | 0.000 | 98.6 | 0.000 | 99.8 | 0.000 | 98.8 |
| LAMP_ORF1b-2_F3 | Orf1b | 100. | 100. | 99.1 | 99.1 | 99.8 | 99.8 | 99.4 | 99.4 |
| LAMP_ORF1b-2_LB | Orf1b | 100. | 100. | 98.0 | 98.0 | 100. | 100. | 99.8 | 99.9 |
| LAMP_ORF1b-2_LF | Orf1b | 99.9 | 100. | 98.6 | 98.6 | 99.8 | 99.8 | 99.5 | 99.6 |

Table S56: Potential drop-out primers for SARS-CoV-2 with partial coverage ($\Delta T_{\text{lim.}}$ = 5°C) below 5%.

| Set | Primer | SARS-CoV-2 | |
|---|---|---|---|
| | | $C_{\text{strict}}$ | $C_{\text{part.}}$ |
| Ganguli et al. [4] | N-P1-F2 | 0.000 | 0.000 |
| Ganguli et al. [4] | Orf8-P1-B3 | 1.25 | 1.25 |
| Ganguli et al. [4] | Orf8-P2-B1c | 0.000 | 0.000 |
| Ganguli et al. [4] | Orf8-P2-B2 | 0.000 | 0.000 |
| Ji et al. [8] | ORFlab-1-LB | 0.000 | 0.000 |
| Rodriguez-Manzano et al. [15] | F2-N | 0.000 | 0.000 |
| Yoshikawa et al. [18] | LAMP_ORF1b-2_B2 | 0.000 | 0.000 |

Table S57: Potential drop-out primers for Alpha variant with partial coverage ($\Delta T_{\mathrm{lim.}}$ = 5°C) below 5%.

| Set | Primer | Alpha | |
|---|---|---|---|
| | | $C_{\mathrm{strict}}$ | $C_{\mathrm{part.}}$ |
| Diego et al. [3] | S447-F2 | 0.911 | 0.911 |
| Ganguli et al. [4] | N-P1-B2 | 0.124 | 0.124 |
| Ganguli et al. [4] | N-P1-F2 | 0.000 | 0.000 |
| Ganguli et al. [4] | Orf8-P1-B2 | 0.179 | 0.179 |
| Ganguli et al. [4] | Orf8-P1-B3 | 0.0690 | 0.0690 |
| Ganguli et al. [4] | Orf8-P1-F1c | 0.0552 | 0.0552 |
| Ganguli et al. [4] | Orf8-P2-B1c | 0.000 | 0.000 |
| Ganguli et al. [4] | Orf8-P2-B2 | 0.000 | 0.000 |
| Ganguli et al. [4] | Orf8-P2-F3 | 0.0552 | 0.0552 |
| Ganguli et al. [4] | Orf8-P3-B3 | 0.0966 | 0.0966 |
| Ganguli et al. [4] | Orf8-P3-F2 | 0.0552 | 0.0552 |
| Ganguli et al. [4] | S-P1-Loop-B | 0.207 | 0.207 |
| Ji et al. [8] | N2-B1c | 0.000 | 0.0690 |
| Ji et al. [8] | ORFlab-1-F2 | 0.152 | 0.152 |
| Ji et al. [8] | ORFlab-1-LB | 0.000 | 0.000 |
| Lau et al. [11] | F1c | 0.221 | 0.235 |
| Lau et al. [11] | FSP | 0.235 | 0.235 |
| Luo et al. [12] | PS1-B2 | 0.0690 | 0.0690 |
| Luo et al. [12] | PS4-B2 | 0.0690 | 0.0690 |
| Luo et al. [12] | PS5-B2 | 0.0690 | 0.0690 |
| Mautner et al. [13] | ORF8-B1c | 0.193 | 0.193 |
| Mautner et al. [13] | ORF8-LF | 0.0552 | 0.0552 |
| Mohon et al. [14] | S3-F3 | 0.110 | 0.110 |
| Rodriguez-Manzano et al. [15] | F2-N | 0.000 | 0.000 |
| Yan et al. [16] | S-123F3 | 0.138 | 0.138 |
| Yang et al. [17] | CU-N2-F1c | 0.0690 | 0.0690 |
| Yoshikawa et al. [18] | LAMP_ORF1b-2_B2 | 0.000 | 0.000 |

Table S58: Potential drop-out primers for Beta variant with partial coverage ($\Delta T_{\text{lim.}}$ = 5°C) below 5%.

| Set | Primer | Beta | |
|---|---|---|---|
| | | $C_{\text{strict}}$ | $C_{\text{part.}}$ |
| Diego et al. [3] | E-B3 | 0.293 | 0.293 |
| Ganguli et al. [4] | N-P1-F2 | 0.000 | 0.000 |
| Ganguli et al. [4] | Orf8-P1-B3 | 0.0267 | 0.0267 |
| Ganguli et al. [4] | Orf8-P2-B1c | 0.000 | 0.000 |
| Ganguli et al. [4] | Orf8-P2-B2 | 0.000 | 0.000 |
| Ganguli et al. [4] | S-P1-B2 | 0.333 | 0.333 |
| Jang et al. [7] | E-B3 | 0.293 | 0.293 |
| Ji et al. [8] | ORFlab-1-LB | 0.000 | 0.000 |
| Lau et al. [11] | FLP | 0.814 | 0.814 |
| Mohon et al. [14] | S2-B1c | 1.99 | 1.99 |
| Mohon et al. [14] | S2-F2 | 2.89 | 2.89 |
| Rodriguez-Manzano et al. [15] | F2-N | 0.000 | 0.000 |
| Yoshikawa et al. [18] | LAMP_ORF1b-2_B2 | 0.000 | 0.000 |

Table S59: Potential drop-out primers for Gamma variant with partial coverage ($\Delta T_{\text{lim.}}$ = 5°C) below 5%.

| | | Gamma | |
|---|---|---|---|
| Set | Primer | $C_{\text{strict}}$ | $C_{\text{part.}}$ |
| Alves et al. [2] | RdRp_B1c | 1.08 | 1.08 |
| Diego et al. [3] | N5-LB | 0.780 | 0.780 |
| Ganguli et al. [4] | N-P1-F2 | 0.000 | 0.000 |
| Ganguli et al. [4] | Orf8-P1-B3 | 0.000 | 0.000 |
| Ganguli et al. [4] | Orf8-P2-B1c | 0.000 | 0.000 |
| Ganguli et al. [4] | Orf8-P2-B2 | 0.000 | 0.000 |
| Ganguli et al. [4] | Orf8-P2-B3 | 2.21 | 2.21 |
| Huang et al. [6] | N15-B2 | 0.563 | 0.563 |
| Ji et al. [8] | N1-LB | 0.780 | 0.780 |
| Ji et al. [8] | N2-B1c | 0.000 | 0.477 |
| Ji et al. [8] | ORFlab-1-LB | 0.000 | 0.000 |
| Luo et al. [12] | PS1-B2 | 0.477 | 0.520 |
| Luo et al. [12] | PS2-LB | 0.780 | 0.780 |
| Luo et al. [12] | PS3-LB | 0.780 | 0.780 |
| Luo et al. [12] | PS4-B2 | 0.477 | 0.520 |
| Luo et al. [12] | PS5-B2 | 0.477 | 0.520 |
| Mautner et al. [13] | ORF8-B3 | 2.21 | 2.21 |
| Rodriguez-Manzano et al. [15] | F2-N | 0.000 | 0.000 |
| Rodriguez-Manzano et al. [15] | F3-N | 0.780 | 0.780 |
| Yang et al. [17] | CU-N2-F1c | 0.477 | 0.563 |
| Yoshikawa et al. [18] | LAMP_ORF1b-2_B2 | 0.000 | 0.000 |

Table S60: Potential drop-out primers for Delta variant with partial coverage ($\Delta T_{\text{lim.}}$ = 5°C) below 5%.

| | | Delta | |
|---|---|---|---|
| Set | Primer | $C_{\text{strict}}$ | $C_{\text{part.}}$ |
| Diego et al. [3] | M-B2 | 0.0126 | 0.0126 |
| Ganguli et al. [4] | N-P1-F2 | 0.000 | 0.000 |
| Ganguli et al. [4] | N-P3-F1c | 0.0881 | 0.0881 |
| Ganguli et al. [4] | Orf8-P1-B3 | 0.000 | 0.000 |
| Ganguli et al. [4] | Orf8-P2-B1c | 0.000 | 0.000 |
| Ganguli et al. [4] | Orf8-P2-B2 | 0.000 | 0.000 |
| Ji et al. [8] | N2-B1c | 0.000 | 0.0126 |
| Ji et al. [8] | ORFlab-1-LB | 0.000 | 0.000 |
| Lau et al. [11] | F1c | 3.78 | 3.85 |
| Lau et al. [11] | FLP | 0.768 | 0.768 |
| Luo et al. [12] | PS1-B2 | 0.000 | 0.000 |
| Luo et al. [12] | PS4-B2 | 0.000 | 0.000 |
| Luo et al. [12] | PS5-B2 | 0.000 | 0.000 |
| Mohon et al. [14] | S3-B3 | 2.19 | 2.19 |
| Rodriguez-Manzano et al. [15] | F2-N | 0.000 | 0.000 |
| Yang et al. [17] | CU-N2-F1c | 0.000 | 0.000 |
| Yoshikawa et al. [18] | LAMP_ORF1b-2_B2 | 0.000 | 0.000 |

Table S61: Potential drop-out primers for Lambda variant with partial coverage ($\Delta T_{\mathrm{lim.}}$ = 5°C) below 5%.

| | | Lambda | |
|---|---|---|---|
| Set | Primer | $C_{\mathrm{strict}}$ | $C_{\mathrm{part.}}$ |
| Alves et al. [2] | N_Set2_F2 | 0.664 | 0.664 |
| Ganguli et al. [4] | N-P1-F2 | 0.000 | 0.000 |
| Ganguli et al. [4] | Orf8-P1-B3 | 0.000 | 0.000 |
| Ganguli et al. [4] | Orf8-P2-B1c | 0.000 | 0.000 |
| Ganguli et al. [4] | Orf8-P2-B2 | 0.000 | 0.000 |
| Ganguli et al. [4] | S-P2-B1c | 0.396 | 0.418 |
| Ganguli et al. [4] | S-P3-B1c | 0.396 | 0.418 |
| Garcia-Venzor et al. [5] | N-geneB3 | 1.04 | 1.04 |
| Huang et al. [6] | N1-F2 | 0.664 | 0.664 |
| Huang et al. [6] | N15-B3 | 0.107 | 0.107 |
| Huang et al. [6] | S17-B1c | 0.396 | 0.418 |
| Ji et al. [8] | N2-B1c | 0.000 | 0.161 |
| Ji et al. [8] | ORFlab-1-LB | 0.000 | 0.000 |
| Lalli et al. [10] | NEB_N2-B3 | 1.03 | 1.03 |
| Lau et al. [11] | B1c | 0.664 | 0.664 |
| Lau et al. [11] | BSP | 0.664 | 0.664 |
| Lau et al. [11] | FLP | 0.739 | 0.739 |
| Luo et al. [12] | PS1-B2 | 0.161 | 0.161 |
| Luo et al. [12] | PS4-B2 | 0.161 | 0.161 |
| Luo et al. [12] | PS4-B3 | 0.107 | 0.107 |
| Luo et al. [12] | PS5-B2 | 0.161 | 0.161 |
| Luo et al. [12] | PS5-B3 | 0.107 | 0.107 |
| Luo et al. [12] | PS6-F2 | 0.664 | 0.664 |
| Mohon et al. [14] | S2-LPB | 2.74 | 2.87 |
| Rodriguez-Manzano et al. [15] | F2-N | 0.000 | 0.000 |
| Yan et al. [16] | S-123F2 | 0.0642 | 0.0642 |
| Yang et al. [17] | CU-N2-F1c | 0.161 | 0.161 |
| Yang et al. [17] | CU-N2-Loop-B | 0.107 | 0.107 |
| Yoshikawa et al. [18] | LAMP_ORF1b-2_B2 | 0.000 | 0.000 |

Table S62: Potential drop-out primers for Mu variant with partial coverage ($\Delta T_{\text{lim.}}$ = 5°C) below 5%.

| Set | Primer | Mu $C_{\text{strict}}$ | Mu $C_{\text{part.}}$ |
|---|---|---|---|
| Alves et al. [2] | E_Set1_LF | 0.756 | 0.953 |
| Ganguli et al. [4] | N-P1-F2 | 0.000 | 0.000 |
| Ganguli et al. [4] | Orf8-P1-B1c | 0.0908 | 0.0908 |
| Ganguli et al. [4] | Orf8-P1-B3 | 0.000 | 0.000 |
| Ganguli et al. [4] | Orf8-P1-F2 | 0.333 | 0.333 |
| Ganguli et al. [4] | Orf8-P2-B1c | 0.000 | 0.000 |
| Ganguli et al. [4] | Orf8-P2-B2 | 0.000 | 0.000 |
| Ganguli et al. [4] | Orf8-P2-F2 | 0.0908 | 0.0908 |
| Ganguli et al. [4] | Orf8-P3-B2 | 3.77 | 3.77 |
| Ganguli et al. [4] | Orf8-P3-F1c | 0.0908 | 0.0908 |
| Ganguli et al. [4] | S-P1-B3 | 0.862 | 0.862 |
| Ganguli et al. [4] | S-P2-B2 | 0.877 | 0.877 |
| Ganguli et al. [4] | S-P3-B2 | 0.877 | 0.877 |
| Huang et al. [6] | S17-B2 | 0.877 | 0.877 |
| Ji et al. [8] | ORFlab-1-LB | 0.000 | 0.000 |
| Mautner et al. [13] | OFR8-B2 | 3.77 | 3.77 |
| Mautner et al. [13] | OFR8-F1c | 0.0908 | 0.0908 |
| Rodriguez-Manzano et al. [15] | F2-N | 0.000 | 0.000 |
| Yoshikawa et al. [18] | LAMP_ORF1b-2_B2 | 0.000 | 0.000 |

Table S63: Potential drop-out primers for Omicron variant with partial coverage ($\Delta T_{\text{lim.}}$ = 5°C) below 5%.

| | | Omicron | |
|---|---|---|---|
| Set | Primer | $C_{\text{strict}}$ | $C_{\text{part.}}$ |
| Alves et al. [2] | E_Set1_B2 | 0.199 | 0.199 |
| Alves et al. [2] | N_Set2_F1c | 0.555 | 0.555 |
| Alves et al. [2] | N_Set2_F2 | 0.640 | 0.640 |
| Diego et al. [3] | M-B1c | 0.569 | 0.569 |
| Diego et al. [3] | N5-F3 | 0.555 | 0.555 |
| Diego et al. [3] | S555-B2 | 0.341 | 0.341 |
| Ganguli et al. [4] | N-P1-F2 | 0.000 | 0.000 |
| Ganguli et al. [4] | Orf8-P1-B3 | 0.000 | 0.000 |
| Ganguli et al. [4] | Orf8-P2-B1c | 0.000 | 0.000 |
| Ganguli et al. [4] | Orf8-P2-B2 | 0.000 | 0.000 |
| Ganguli et al. [4] | S-P1-B1c | 0.498 | 0.498 |
| Ganguli et al. [4] | S-P1-B3 | 0.341 | 0.341 |
| Ganguli et al. [4] | S-P1-Loop-B | 0.441 | 0.455 |
| Ganguli et al. [4] | S-P2-B2 | 0.341 | 0.341 |
| Ganguli et al. [4] | S-P3-B2 | 0.341 | 0.341 |
| Huang et al. [6] | N1-F1c | 0.555 | 0.555 |
| Huang et al. [6] | N1-F2 | 0.640 | 0.640 |
| Huang et al. [6] | S17-B2 | 0.341 | 0.341 |
| Jang et al. [7] | E-F2 | 0.185 | 0.185 |
| Ji et al. [8] | N2-B1c | 0.000 | 0.0854 |
| Ji et al. [8] | ORFlab-1-LB | 0.000 | 0.000 |
| Jiang et al. [9] | nCoV-N-F3 | 0.555 | 0.555 |
| Lalli et al. [10] | NEB_E1-F2 | 0.199 | 0.199 |
| Lau et al. [11] | B1c | 0.640 | 0.640 |
| Lau et al. [11] | B2 | 0.555 | 0.555 |
| Lau et al. [11] | BSP | 0.640 | 0.640 |
| Luo et al. [12] | PS1-B2 | 0.0854 | 0.0854 |
| Luo et al. [12] | PS2-F3 | 0.555 | 0.555 |
| Luo et al. [12] | PS3-F3 | 0.555 | 0.555 |
| Luo et al. [12] | PS4-B2 | 0.0854 | 0.0854 |
| Luo et al. [12] | PS5-B2 | 0.0854 | 0.0854 |
| Luo et al. [12] | PS6-F1c | 0.555 | 0.555 |
| Luo et al. [12] | PS6-F2 | 0.640 | 0.640 |
| Mohon et al. [14] | S2-F2 | 0.512 | 0.512 |
| Rodriguez-Manzano et al. [15] | F2-N | 0.000 | 0.000 |
| Yan et al. [16] | S-123LB | 0.726 | 0.726 |

(continued on next page)

(Table S63, Drop-out primers, continued)

| Set | Primer | Omicron | |
|---|---|---|---|
| | | $C_{\text{strict}}$ | $C_{\text{part.}}$ |
| Yang et al. [17] | CU-N2-F1c | 0.0854 | 0.0854 |
| Yoshikawa et al. [18] | LAMP_ORF1b-2_B2 | 0.000 | 0.000 |

Table S64: Potential drop-out primers for BA.2 subvariant with partial coverage ($\Delta T_{\text{lim.}}$ = 5°C) below 5%.

| | | BA.2 | |
|---|---|---|---|
| Set | Primer | $C_{\text{strict}}$ | $C_{\text{part.}}$ |
| Alves et al. [2] | E_Set1_B2 | 0.000 | 0.000 |
| Alves et al. [2] | N_Set2_F1c | 1.31 | 1.31 |
| Alves et al. [2] | N_Set2_F2 | 0.000 | 0.000 |
| Diego et al. [3] | M-B1c | 0.000 | 0.000 |
| Diego et al. [3] | N5-F3 | 1.31 | 1.31 |
| Diego et al. [3] | S555-B2 | 0.000 | 0.000 |
| Ganguli et al. [4] | N-P1-F2 | 0.000 | 0.000 |
| Ganguli et al. [4] | N-P3-B3 | 0.0676 | 0.0676 |
| Ganguli et al. [4] | Orf8-P1-B3 | 0.0135 | 0.0135 |
| Ganguli et al. [4] | Orf8-P2-B1c | 0.000 | 0.000 |
| Ganguli et al. [4] | Orf8-P2-B2 | 0.000 | 0.000 |
| Huang et al. [6] | N1-F1c | 1.31 | 1.31 |
| Huang et al. [6] | N1-F2 | 0.000 | 0.000 |
| Jang et al. [7] | E-F2 | 0.000 | 0.000 |
| Jang et al. [7] | RdRP-B1c | 0.0406 | 0.0406 |
| Ji et al. [8] | N2-B1c | 0.000 | 0.0947 |
| Ji et al. [8] | ORFlab-1-LB | 0.000 | 0.000 |
| Jiang et al. [9] | nCoV-N-F3 | 1.31 | 1.31 |
| Lalli et al. [10] | NEB_E1-F2 | 0.000 | 0.000 |
| Lalli et al. [10] | NEB_orf1a-A-B2 | 0.189 | 0.189 |
| Lau et al. [11] | B1c | 0.000 | 0.000 |
| Lau et al. [11] | B2 | 1.31 | 1.31 |
| Lau et al. [11] | BSP | 0.000 | 0.000 |
| Luo et al. [12] | PS1-B2 | 0.0947 | 0.0947 |
| Luo et al. [12] | PS2-F3 | 1.31 | 1.31 |
| Luo et al. [12] | PS3-F3 | 1.31 | 1.31 |
| Luo et al. [12] | PS4-B2 | 0.0947 | 0.0947 |
| Luo et al. [12] | PS5-B2 | 0.0947 | 0.0947 |
| Luo et al. [12] | PS6-F1c | 1.31 | 1.31 |
| Luo et al. [12] | PS6-F2 | 0.000 | 0.000 |
| Mohon et al. [14] | S2-F2 | 0.000 | 0.000 |
| Rodriguez-Manzano et al. [15] | F2-N | 0.000 | 0.000 |
| Yan et al. [16] | S-123LB | 0.0135 | 0.0135 |
| Yang et al. [17] | CU-N2-F1c | 0.0947 | 0.0947 |
| Yoshikawa et al. [18] | LAMP_ORF1b-2_B2 | 0.000 | 0.000 |

Table S65: Potential drop-out primers for BA.3 subvariant with partial coverage ($\Delta T_{\text{lim.}}$ = 5°C) below 5%.

| Set | Primer | BA.3 | |
|---|---|---|---|
| | | $C_{\text{strict}}$ | $C_{\text{part.}}$ |
| Alves et al. [2] | E_Set1_B2 | 0.000 | 0.000 |
| Alves et al. [2] | N_Set2_F1c | 0.862 | 0.862 |
| Alves et al. [2] | N_Set2_F2 | 1.15 | 1.15 |
| Diego et al. [3] | M-B1c | 0.287 | 0.287 |
| Diego et al. [3] | N5-F3 | 0.862 | 0.862 |
| Diego et al. [3] | S555-B2 | 0.000 | 0.000 |
| Ganguli et al. [4] | N-P1-F2 | 0.000 | 0.000 |
| Ganguli et al. [4] | N-P3-B3 | 1.15 | 1.15 |
| Ganguli et al. [4] | Orf8-P1-B3 | 0.000 | 0.000 |
| Ganguli et al. [4] | Orf8-P2-B1c | 0.000 | 0.000 |
| Ganguli et al. [4] | Orf8-P2-B2 | 0.000 | 0.000 |
| Ganguli et al. [4] | S-P1-B1c | 0.862 | 0.862 |
| Ganguli et al. [4] | S-P1-B3 | 0.862 | 0.862 |
| Ganguli et al. [4] | S-P1-Loop-B | 0.287 | 0.287 |
| Ganguli et al. [4] | S-P2-B2 | 0.862 | 0.862 |
| Ganguli et al. [4] | S-P3-B2 | 0.862 | 0.862 |
| Huang et al. [6] | N1-F1c | 0.862 | 0.862 |
| Huang et al. [6] | N1-F2 | 1.15 | 1.15 |
| Huang et al. [6] | S17-B2 | 0.862 | 0.862 |
| Jang et al. [7] | E-F2 | 0.000 | 0.000 |
| Jang et al. [7] | RdRP-B1c | 0.575 | 0.575 |
| Ji et al. [8] | N2-B1c | 0.000 | 0.000 |
| Ji et al. [8] | ORFlab-1-LB | 0.000 | 0.000 |
| Jiang et al. [9] | nCoV-N-F3 | 0.862 | 0.862 |
| Lalli et al. [10] | NEB_E1-F2 | 0.000 | 0.000 |
| Lalli et al. [10] | NEB_orf1a-A-B2 | 0.575 | 0.575 |
| Lau et al. [11] | B1c | 1.15 | 1.15 |
| Lau et al. [11] | B2 | 0.862 | 0.862 |
| Lau et al. [11] | BSP | 1.15 | 1.15 |
| Luo et al. [12] | PS1-B2 | 0.000 | 0.000 |
| Luo et al. [12] | PS2-F3 | 0.862 | 0.862 |
| Luo et al. [12] | PS3-F3 | 0.862 | 0.862 |
| Luo et al. [12] | PS4-B2 | 0.000 | 0.000 |
| Luo et al. [12] | PS5-B2 | 0.000 | 0.000 |
| Luo et al. [12] | PS6-F1c | 0.862 | 0.862 |
| Luo et al. [12] | PS6-F2 | 1.15 | 1.15 |

(continued on next page)

(Table S65, Drop-out primers, continued)

| Set | Primer | BA.3 | |
|---|---|---|---|
| | | $C_{\text{strict}}$ | $C_{\text{part.}}$ |
| Mohon et al. [14] | S2-F2 | 0.000 | 0.000 |
| Rodriguez-Manzano et al. [15] | F2-N | 0.000 | 0.000 |
| Yan et al. [16] | S-123LB | 0.000 | 0.000 |
| Yang et al. [17] | CU-N2-F1c | 0.000 | 0.000 |
| Yoshikawa et al. [18] | LAMP_ORF1b-2_B2 | 0.000 | 0.000 |

Table S66: Potential drop-out primers for BA.4 subvariant with partial coverage ($\Delta T_{\text{lim.}}$ = 5°C) below 5%.

| Set | Primer | BA.4 | |
|---|---|---|---|
| | | $C_{\text{strict}}$ | $C_{\text{part.}}$ |
| Alves et al. [2] | E_Set1_B2 | 0.159 | 0.159 |
| Alves et al. [2] | N_Set1_B3 | 0.954 | 0.954 |
| Alves et al. [2] | N_Set2_F1c | 0.318 | 0.318 |
| Alves et al. [2] | N_Set2_F2 | 0.477 | 0.477 |
| Diego et al. [3] | M-B1c | 0.000 | 0.000 |
| Diego et al. [3] | N5-F3 | 0.318 | 0.318 |
| Diego et al. [3] | S555-B2 | 0.000 | 0.000 |
| Ganguli et al. [4] | N-P1-F2 | 0.000 | 0.000 |
| Ganguli et al. [4] | N-P2-B3 | 0.954 | 0.954 |
| Ganguli et al. [4] | N-P3-B3 | 0.000 | 0.000 |
| Ganguli et al. [4] | Orf8-P1-B3 | 0.000 | 0.000 |
| Ganguli et al. [4] | Orf8-P2-B1c | 0.000 | 0.000 |
| Ganguli et al. [4] | Orf8-P2-B2 | 0.000 | 0.000 |
| Ganguli et al. [4] | S-P1-Loop-B | 3.02 | 3.18 |
| Huang et al. [6] | N1-F1c | 0.318 | 0.318 |
| Huang et al. [6] | N1-F2 | 0.477 | 0.477 |
| Jang et al. [7] | E-F2 | 0.159 | 0.159 |
| Jang et al. [7] | RdRP-B1c | 0.000 | 0.000 |
| Ji et al. [8] | N2-B1c | 0.000 | 0.000 |
| Ji et al. [8] | N2-F3 | 0.954 | 0.954 |
| Ji et al. [8] | ORFlab-1-LB | 0.000 | 0.000 |
| Jiang et al. [9] | nCoV-N-F3 | 0.318 | 0.318 |
| Lalli et al. [10] | NEB_E1-F2 | 0.159 | 0.159 |
| Lalli et al. [10] | NEB_geneN-A-B3 | 0.954 | 0.954 |
| Lalli et al. [10] | NEB_orf1a-A-B2 | 0.159 | 0.159 |
| Lalli et al. [10] | NEB_orf1a-A-B3 | 4.45 | 4.45 |
| Lau et al. [11] | B1c | 0.477 | 0.477 |
| Lau et al. [11] | B2 | 0.318 | 0.318 |
| Lau et al. [11] | BSP | 0.477 | 0.477 |
| Luo et al. [12] | PS1-B2 | 0.000 | 0.000 |
| Luo et al. [12] | PS1-F3 | 0.954 | 0.954 |
| Luo et al. [12] | PS2-F3 | 0.318 | 0.318 |
| Luo et al. [12] | PS3-F3 | 0.318 | 0.318 |
| Luo et al. [12] | PS4-B2 | 0.000 | 0.000 |
| Luo et al. [12] | PS5-B2 | 0.000 | 0.000 |
| Luo et al. [12] | PS6-F1c | 0.318 | 0.318 |

(continued on next page)

(Table S66, Drop-out primers, continued)

| Set | Primer | BA.4 | |
|---|---|---|---|
| | | $C_{\text{strict}}$ | $C_{\text{part.}}$ |
| Luo et al. [12] | PS6-F2 | 0.477 | 0.477 |
| Mohon et al. [14] | S2-F2 | 0.318 | 0.318 |
| Rodriguez-Manzano et al. [15] | F2-N | 0.000 | 0.000 |
| Yan et al. [16] | S-123LB | 0.636 | 0.795 |
| Yang et al. [17] | CU-N2-F1c | 0.000 | 0.000 |
| Yoshikawa et al. [18] | LAMP_ORF1b-2_B2 | 0.000 | 0.000 |

Table S67: Potential drop-out primers for BA.5 subvariant with partial coverage ($\Delta T_{\text{lim.}}$ = 5°C) below 5%.

| | | BA.5 | |
|---|---|---|---|
| Set | Primer | $C_{\text{strict}}$ | $C_{\text{part.}}$ |
| Alves et al. [2] | E_Set1_B2 | 0.162 | 0.162 |
| Alves et al. [2] | N_Set2_F1c | 0.975 | 1.14 |
| Alves et al. [2] | N_Set2_F2 | 0.569 | 0.569 |
| Diego et al. [3] | M-B1c | 0.000 | 0.000 |
| Diego et al. [3] | N5-F3 | 0.975 | 0.975 |
| Diego et al. [3] | S555-B2 | 0.244 | 0.244 |
| Ganguli et al. [4] | N-P1-F2 | 0.000 | 0.000 |
| Ganguli et al. [4] | N-P3-B3 | 0.0812 | 0.0812 |
| Ganguli et al. [4] | Orf8-P1-B3 | 0.000 | 0.000 |
| Ganguli et al. [4] | Orf8-P2-B1c | 0.000 | 0.000 |
| Ganguli et al. [4] | Orf8-P2-B2 | 0.000 | 0.000 |
| Ganguli et al. [4] | S-P1-Loop-B | 4.55 | 4.87 |
| Huang et al. [6] | N1-F1c | 0.975 | 1.14 |
| Huang et al. [6] | N1-F2 | 0.569 | 0.569 |
| Jang et al. [7] | E-F2 | 0.162 | 0.162 |
| Jang et al. [7] | RdRP-B1c | 0.0812 | 0.0812 |
| Ji et al. [8] | N2-B1c | 0.000 | 0.000 |
| Ji et al. [8] | ORFlab-1-LB | 0.000 | 0.000 |
| Jiang et al. [9] | nCoV-N-F3 | 0.975 | 0.975 |
| Lalli et al. [10] | NEB_E1-F2 | 0.162 | 0.162 |
| Lalli et al. [10] | NEB_orf1a-A-B2 | 0.325 | 0.325 |
| Lau et al. [11] | B1c | 0.569 | 0.569 |
| Lau et al. [11] | B2 | 0.975 | 0.975 |
| Lau et al. [11] | BSP | 0.569 | 0.569 |
| Luo et al. [12] | PS1-B2 | 0.000 | 0.000 |
| Luo et al. [12] | PS2-F3 | 0.975 | 0.975 |
| Luo et al. [12] | PS3-F3 | 0.975 | 0.975 |
| Luo et al. [12] | PS4-B2 | 0.000 | 0.000 |
| Luo et al. [12] | PS5-B2 | 0.000 | 0.000 |
| Luo et al. [12] | PS6-F1c | 0.975 | 1.14 |
| Luo et al. [12] | PS6-F2 | 0.569 | 0.569 |
| Mohon et al. [14] | S2-F2 | 0.325 | 0.325 |
| Rodriguez-Manzano et al. [15] | F2-N | 0.000 | 0.000 |
| Yan et al. [16] | S-123LB | 0.569 | 0.569 |
| Yang et al. [17] | CU-N2-F1c | 0.000 | 0.000 |
| Yoshikawa et al. [18] | LAMP_ORF1b-2_B2 | 0.000 | 0.000 |

Table S68: Percentage of mismatched base pairs at the first three positions of the 5′ end and the last three position of the 3′ end of FIP and BIP primers for 21665 genomes of SARS-CoV-2. $N_{\mathrm{al.}}$ is the number of alignments for each primer. Only those primers with $C_{\mathrm{strict}}(5°C) = 0$ and $C_{\mathrm{part.}}(5°C) > 90\%$.

| Primer | Gene Target | $C_{\mathrm{part.}}$ (%) | $N_{\mathrm{al.}}$ | 5′ end ⟶ (%) | | | | | ⟶ 3′ end (%) |
|---|---|---|---|---|---|---|---|---|---|
| As1_F1c [1] | Orf1a | 99.3 | 21512 | 0 | 0 | 0 | 0 | 0 | 100. |
| As1e_F1c [1] | Orf1a | 99.3 | 21564 | 0 | 0 | 0 | 0 | 99.7 | 99.7 |
| iLACO-F1c [1] | Orf1ab | 99.5 | 21547 | 0 | 0 | 0 | 0 | 0 | 100. |
| E-F1c [3] | E | 99.2 | 21508 | 0 | 0 | 0 | 0 | 0 | 99.9 |
| S447-B1c [3] | S | 99.4 | 21539 | 0 | 0 | 0 | 100. | 100. | 0 |
| N15-B1c [6] | N | 99.1 | 21475 | 0 | 0 | 0 | 0 | 99.9 | 0.084 |
| E-B2 [7] | E | 99.0 | 21486 | 0 | 0 | 99.8 | 0 | 0 | 0 |
| N1-B1c [8] | N | 99.4 | 21533 | 0 | 0 | 0 | 0 | 0 | 1.47 |
| N1-F1c [8] | N | 99.0 | 21547 | 0 | 0.032 | 0 | 99.8 | 99.8 | 0 |
| N2-F1c [8] | N | 99.1 | 21476 | 0 | 0 | 0 | 100. | 100. | 100. |
| ORFlab-1-F1c [8] | Orf1ab | 99.2 | 21534 | 0.032 | 0 | 0 | 0.028 | 99.8 | 99.8 |
| NEB_orf1a-A-F1c [10] | Orf1a | 98.9 | 21445 | 0.047 | 0 | 0 | 0 | 0 | 99.9 |
| ORF1e-B1c [17] | Orf1ab | 99.2 | 21534 | 0 | 0 | 0 | 0 | 0.014 | 99.7 |
| LAMP_ORF1b-2_F2 [18] | Orf1b | 98.8 | 21499 | 99.6 | 0 | 0 | 0 | 0 | 0 |

Table S69: Percentage of mismatched base pairs at the first three positions of the 5′ end and the last three position of the 3′ end of FIP and BIP primers for 7247 genomes of Alpha variant. $N_{al.}$ is the number of alignments for each primer. Only those primers with $C_{strict}(5°C) = 0$ and $C_{part.}(5°C) > 90\%$.

| Primer | Gene Target | $C_{part.}$ (%) | $N_{al.}$ | 5′ end ⟶ | | | | | ⟶ 3′ end |
|---|---|---|---|---|---|---|---|---|---|
| As1_F1c [1] | Orf1a | 99.4 | 7213 | 0 | 0 | 0 | 0 | 0 | 99.9 |
| As1e_F1c [1] | Orf1a | 99.5 | 7238 | 0 | 0 | 0 | 0 | 99.6 | 99.6 |
| iLACO-F1c [1] | Orf1ab | 99.7 | 7231 | 0 | 0 | 99.8 | 0 | 0 | 99.9 |
| E-F1c [3] | E | 99.6 | 7223 | 0 | 0 | 0 | 0 | 0 | 99.9 |
| S447-B1c [3] | S | 99.8 | 7234 | 0 | 0 | 0 | 99.9 | 99.9 | 0 |
| N15-B1c [6] | N | 99.6 | 7220 | 0.32 | 0 | 0 | 0 | 99.9 | 0.055 |
| E-B2 [7] | E | 99.5 | 7226 | 0 | 0 | 99.7 | 0 | 0 | 0 |
| N1-B1c [8] | N | 99.9 | 7240 | 0 | 0.028 | 0 | 0 | 0 | 0.41 |
| N1-F1c [8] | N | 99.8 | 7244 | 0 | 0 | 0 | 99.8 | 99.8 | 0 |
| N2-F1c [8] | N | 99.6 | 7219 | 0.014 | 0.32 | 0 | 100. | 100. | 100. |
| ORFlab-1-F1c [8] | Orf1ab | 99.5 | 7235 | 0 | 0 | 0 | 0 | 99.7 | 99.7 |
| NEB_orf1a-A-F1c [10] | Orf1a | 99.7 | 7230 | 0.18 | 0 | 0 | 0 | 0 | 99.9 |
| ORF1e-B1c [17] | Orf1ab | 98.9 | 7227 | 0 | 0 | 0 | 0 | 0 | 99.2 |
| LAMP_ORF1b-2_F2 [18] | Orf1b | 98.2 | 7168 | 99.2 | 0 | 0 | 0 | 0 | 0 |

Table S70: Percentage of mismatched base pairs at the first three positions of the 5′ end and the last three position of the 3′ end of FIP and BIP primers for 7497 genomes of Beta variant. $N_{\text{al.}}$ is the number of alignments for each primer. Only those primers with $C_{\text{strict}}(5°\text{C}) = 0$ and $C_{\text{part.}}(5°\text{C}) > 90\%$.

| Primer | Gene Target | $C_{\text{part.}}$ (%) | $N_{\text{al.}}$ | 5′ end ⟶ (%) | | | | | ⟶ 3′ end (%) |
|---|---|---|---|---|---|---|---|---|---|
| As1_F1c [1] | Orf1a | 98.7 | 7406 | 0 | 0 | 0 | 0 | 0 | 99.9 |
| As1e_F1c [1] | Orf1a | 98.8 | 7488 | 0 | 0 | 0 | 0 | 98.9 | 98.9 |
| iLACO-F1c [1] | Orf1ab | 99.1 | 7432 | 0 | 0.013 | 0.013 | 0 | 0 | 100. |
| E-F1c [3] | E | 99.3 | 7449 | 0 | 0 | 0 | 0 | 0 | 99.9 |
| S447-B1c [3] | S | 99.7 | 7479 | 0 | 0 | 0 | 99.9 | 99.9 | 0 |
| N15-B1c [6] | N | 96.6 | 7261 | 0.027 | 0 | 0 | 0 | 99.8 | 0.014 |
| E-B2 [7] | E | 99.1 | 7444 | 0 | 0 | 99.8 | 0.013 | 0 | 0 |
| N1-B1c [8] | N | 97.8 | 7333 | 0 | 0 | 0 | 0 | 0 | 0.76 |
| N1-F1c [8] | N | 98.4 | 7446 | 0 | 0 | 0 | 99.1 | 99.1 | 0 |
| N2-B1c [8] | N | 96.4 | 7239 | 0 | 0 | 0 | 0 | 99.8 | 99.8 |
| N2-F1c [8] | N | 96.7 | 7251 | 0.027 | 0.027 | 0 | 100. | 100. | 100. |
| ORFlab-1-F1c [8] | Orf1ab | 96.8 | 7413 | 0 | 0 | 0 | 0 | 97.9 | 97.9 |
| NEB_orf1a-A-F1c [10] | Orf1a | 99.3 | 7465 | 0.014 | 0 | 0 | 0 | 0 | 99.7 |
| ORF1e-B1c [17] | Orf1ab | 98.9 | 7418 | 0.108 | 0 | 0 | 0 | 0 | 99.9 |
| LAMP_ORF1b-2_F2 [18] | Orf1b | 94.1 | 7317 | 96.4 | 0 | 0 | 0 | 0 | 0 |

Table S71: Percentage of mismatched base pairs at the first three positions of the 5′ end and the last three position of the 3′ end of FIP and BIP primers for 2308 genomes of Gamma variant. $N_{al.}$ is the number of alignments for each primer. Only those primers with $C_{strict}(5°C) = 0$ and $C_{part.}(5°C) > 90\%$.

| Primer | Gene Target | $C_{part.}$ (%) | $N_{al.}$ | 5′ end ⟶ (%) | | | | | ⟶ 3′ end (%) |
|---|---|---|---|---|---|---|---|---|---|
| As1_F1c [1] | Orf1a | 99.7 | 2304 | 0 | 0 | 0 | 0 | 0 | 99.9 |
| As1e_F1c [1] | Orf1a | 99.8 | 2308 | 0 | 0 | 0 | 0 | 99.8 | 99.8 |
| iLACO-F1c [1] | Orf1ab | 99.5 | 2297 | 0 | 0 | 0.043 | 0 | 0 | 100. |
| E-F1c [3] | E | 99.8 | 2306 | 0 | 0 | 0 | 0 | 0 | 99.9 |
| S447-B1c [3] | S | 99.9 | 2308 | 0 | 0 | 0 | 99.9 | 99.9 | 0 |
| N15-B1c [6] | N | 98.6 | 2305 | 0 | 0 | 0 | 0 | 98.7 | 0.043 |
| E-B2 [7] | E | 99.7 | 2306 | 0 | 0 | 99.8 | 0 | 0 | 0 |
| N1-B1c [8] | N | 99.7 | 2300 | 0 | 0 | 0 | 0 | 0 | 0.043 |
| N1-F1c [8] | N | 98.7 | 2288 | 0 | 0 | 0 | 99.6 | 99.6 | 0 |
| N2-F1c [8] | N | 99.9 | 2305 | 0 | 0 | 0 | 100. | 100. | 100. |
| ORFlab-1-F1c [8] | Orf1ab | 99.1 | 2293 | 0 | 0 | 0 | 0.087 | 99.7 | 99.7 |
| NEB_orf1a-A-F1c [10] | Orf1a | 99.8 | 2303 | 0.087 | 0.217 | 0 | 0 | 0 | 100. |
| ORF1e-B1c [17] | Orf1ab | 100. | 2308 | 0 | 0 | 0 | 0 | 0.043 | 100. |
| LAMP_ORF1b-2_F2 [18] | Orf1b | 95.8 | 2232 | 99.1 | 0 | 0 | 0 | 0 | 0 |

Table S72: Percentage of mismatched base pairs at the first three positions of the 5′ end and the last three position of the 3′ end of FIP and BIP primers for 7943 genomes of Delta variant. $N_{\text{al.}}$ is the number of alignments for each primer. Only those primers with $C_{\text{strict}}(5°\text{C}) = 0$ and $C_{\text{part.}}(5°\text{C}) > 90\%$.

| Primer | Gene Target | $C_{\text{part.}}$ (%) | $N_{\text{al.}}$ | 5′ end ⟶ (%) | | | | | ⟶ 3′ end (%) |
|---|---|---|---|---|---|---|---|---|---|
| As1_F1c [1] | Orf1a | 99.2 | 7943 | 0 | 0 | 0 | 0.012 | 0.012 | 99.2 |
| As1e_F1c [1] | Orf1a | 100. | 7943 | 0 | 0.82 | 0 | 0 | 100. | 100. |
| iLACO-F1c [1] | Orf1ab | 100. | 7943 | 0 | 0.012 | 0.012 | 0 | 0 | 100. |
| E-F1c [3] | E | 99.9 | 7936 | 0 | 0 | 0 | 0 | 0 | 99.9 |
| S447-B1c [3] | S | 100. | 7942 | 0 | 0 | 0 | 100. | 100. | 0 |
| N15-B1c [6] | N | 100. | 7943 | 0 | 0 | 0 | 0 | 100. | 0 |
| E-B2 [7] | E | 99.6 | 7943 | 0 | 0 | 99.6 | 0 | 0 | 0 |
| N1-B1c [8] | N | 100. | 7941 | 0 | 0 | 0 | 0 | 0 | 0.43 |
| N1-F1c [8] | N | 99.8 | 7939 | 0 | 0.025 | 0 | 99.9 | 99.9 | 0 |
| N2-F1c [8] | N | 100. | 7942 | 0.012 | 0 | 0 | 100. | 100. | 100. |
| ORFlab-1-F1c [8] | Orf1ab | 99.8 | 7941 | 0.012 | 0 | 0 | 0.012 | 99.8 | 99.8 |
| NEB_orf1a-A-F1c [10] | Orf1a | 99.8 | 7939 | 0.012 | 0 | 0 | 0.012 | 0 | 99.9 |
| ORF1e-B1c [17] | Orf1ab | 100. | 7943 | 0 | 0 | 0 | 0 | 0.012 | 99.9 |
| LAMP_ORF1b-2_F2 [18] | Orf1b | 99.1 | 7886 | 99.9 | 0 | 0 | 0 | 0 | 0 |

Table S73: Percentage of mismatched base pairs at the first three positions of the 5′ end and the last three position of the 3′ end of FIP and BIP primers for 9340 genomes of Lambda variant. $N_{\text{al.}}$ is the number of alignments for each primer. Only those primers with $C_{\text{strict}}(5°C) = 0$ and $C_{\text{part.}}(5°C) > 90\%$.

| Primer | Gene Target | $C_{\text{part.}}$ (%) | $N_{\text{al.}}$ | 5′ end ⟶ (%) | | | ⟶ 3′ end (%) | | |
|---|---|---|---|---|---|---|---|---|---|
| As1_F1c [1] | Orf1a | 99.1 | 9332 | 0 | 0 | 0 | 0.032 | 0 | 99.1 |
| As1e_F1c [1] | Orf1a | 99.4 | 9338 | 0 | 0 | 0.043 | 0 | 99.4 | 99.4 |
| iLACO-F1c [1] | Orf1ab | 100. | 9337 | 0 | 0 | 0.043 | 0 | 0 | 100. |
| E-F1c [3] | E | 99.1 | 9268 | 0 | 0 | 0 | 0 | 0 | 99.9 |
| S447-B1c [3] | S | 99.7 | 9335 | 0 | 0 | 0 | 99.8 | 99.8 | 0 |
| E-B2 [7] | E | 98.4 | 9267 | 0 | 0 | 99.1 | 0 | 0 | 0 |
| N1-B1c [8] | N | 99.9 | 9335 | 0 | 0.021 | 0 | 0 | 0 | 0.30 |
| N1-F1c [8] | N | 99.5 | 9336 | 0 | 0.22 | 0.021 | 99.6 | 99.6 | 0 |
| N2-F1c [8] | N | 99.9 | 9329 | 0.021 | 0.011 | 0 | 100. | 100. | 100. |
| ORFlab-1-F1c [8] | Orf1ab | 99.4 | 9338 | 0.28 | 0 | 0 | 1.04 | 99.4 | 99.4 |
| NEB_orf1a-A-F1c [10] | Orf1a | 99.9 | 9331 | 0.15 | 0.021 | 0 | 0 | 0 | 99.9 |
| ORF1e-B1c [17] | Orf1ab | 99.7 | 9340 | 0 | 0.021 | 0 | 0 | 0.075 | 99.7 |
| LAMP_ORF1b-2_F2 [18] | Orf1b | 96.1 | 9116 | 98.4 | 0 | 0 | 0 | 0 | 0 |

Table S74: Percentage of mismatched base pairs at the first three positions of the 5′ end and the last three position of the 3′ end of FIP and BIP primers for 6610 genomes of Mu variant. $N_{al.}$ is the number of alignments for each primer. Only those primers with $C_{strict}(5°C) = 0$ and $C_{part.}(5°C) > 90\%$.

| Primer | Gene Target | $C_{part.}$ (%) | $N_{al.}$ | 5′ end ⟶ (%) | | | | | ⟶ 3′ end (%) |
|---|---|---|---|---|---|---|---|---|---|
| As1_F1c [1] | Orf1a | 99.7 | 6608 | 0.045 | 0 | 0 | 0.015 | 0 | 99.8 |
| As1e_F1c [1] | Orf1a | 99.9 | 6609 | 0.045 | 0 | 0 | 0 | 99.9 | 99.9 |
| iLACO-F1c [1] | Orf1ab | 100. | 6608 | 0 | 0.06 | 0.03 | 0 | 0 | 100. |
| E-F1c [3] | E | 99.5 | 6580 | 0 | 0 | 0 | 0 | 0 | 99.9 |
| S447-B1c [3] | S | 99.9 | 6607 | 0 | 0 | 0 | 100. | 100. | 0 |
| N15-B1c [6] | N | 99.8 | 6601 | 0.015 | 0 | 0 | 0 | 99.9 | 0.106 |
| E-B2 [7] | E | 99.4 | 6587 | 0 | 0 | 99.8 | 0 | 0 | 0 |
| N1-B1c [8] | N | 99.9 | 6602 | 6 | 0 | 0 | 0 | 0 | 2.68 |
| N2-B1c [8] | N | 99.3 | 6599 | 0 | 0 | 0 | 0 | 99.5 | 99.6 |
| N2-F1c [8] | N | 99.9 | 6601 | 0.303 | 0.015 | 0 | 100. | 100. | 100. |
| ORFlab-1-F1c [8] | Orf1ab | 99.7 | 6601 | 0 | 0 | 0 | 0.076 | 99.9 | 99.9 |
| NEB_orf1a-A-F1c [10] | Orf1a | 99.6 | 6604 | 0.35 | 0.015 | 0 | 0 | 0 | 99.6 |
| ORF1e-B1c [17] | Orf1ab | 99.6 | 6605 | 0 | 0 | 0 | 0 | 0 | 99.7 |
| LAMP_ORF1b-2_F2 [18] | Orf1b | 94.7 | 6373 | 98.2 | 0 | 0 | 0 | 0 | 0 |

Table S75: Percentage of mismatched base pairs at the first three positions of the 5′ end and the last three position of the 3′ end of FIP and BIP primers for 7029 genomes of Omicron variant. $N_{\text{al.}}$ is the number of alignments for each primer. Only those primers with $C_{\text{strict}}(5°C) = 0$ and $C_{\text{part.}}(5°C) > 90\%$.

| Primer | Gene Target | $C_{\text{part.}}$ (%) | $N_{\text{al.}}$ | 5′ end ⟶ (%) | | | ⟶ 3′ end (%) | | |
|---|---|---|---|---|---|---|---|---|---|
| As1_F1c [1] | Orf1a | 98.6 | 6930 | 0 | 0 | 0 | 0.014 | 0 | 100. |
| As1e_F1c [1] | Orf1a | 98.6 | 7020 | 0 | 0 | 0 | 0 | 98.7 | 98.7 |
| iLACO-F1c [1] | Orf1ab | 99.5 | 6996 | 0 | 0 | 0 | 0 | 0 | 100. |
| E-F1c [3] | E | 97.0 | 6818 | 0 | 0 | 0 | 0 | 0 | 100. |
| S447-B1c [3] | S | 99.8 | 7015 | 0 | 0 | 0 | 100. | 100. | 0 |
| N15-B1c [6] | N | 99.2 | 6977 | 0 | 0 | 0 | 0 | 100. | 0 |
| E-B2 [7] | E | 94.2 | 6625 | 0 | 0 | 99.9 | 0 | 0 | 0 |
| N1-B1c [8] | N | 94.0 | 6609 | 0 | 0 | 0 | 0 | 0 | 0.21 |
| N1-F1c [8] | N | 93.4 | 6567 | 0 | 0.015 | 0 | 99.9 | 99.9 | 99.3 |
| N2-F1c [8] | N | 99.3 | 6978 | 0 | 0 | 0 | 100. | 100. | 100. |
| ORFlab-1-F1c [8] | Orf1ab | 99.7 | 7013 | 0 | 0 | 0 | 0 | 99.9 | 99.9 |
| ORF1e-B1c [17] | Orf1ab | 99.0 | 6972 | 0 | 0 | 0 | 0 | 0 | 99.8 |
| LAMP_ORF1b-2_F2 [18] | Orf1b | 98.8 | 6955 | 99.8 | 0 | 0 | 0 | 0 | 0 |

Table S76: Percentage of mismatched base pairs at the first three positions of the 5′ end and the last three position of the 3′ end of FIP and BIP primers for 7393 genomes of BA.2 subvariant. $N_{\text{al.}}$ is the number of alignments for each primer. Only those primers with $C_{\text{strict}}(5°C) = 0$ and $C_{\text{part.}}(5°C) > 90\%$.

| Primer | Gene Target | $C_{\text{part.}}$ (%) | $N_{\text{al.}}$ | 5′ end ⟶ (%) | | | | | ⟶ 3′ end (%) |
|---|---|---|---|---|---|---|---|---|---|
| As1_F1c [1] | Orf1a | 100. | 7392 | 0.013 | 0 | 0 | 0 | 0 | 100. |
| As1e_F1c [1] | Orf1a | 100. | 7392 | 0.013 | 0 | 0 | 0 | 100. | 100. |
| iLACO-F1c [1] | Orf1ab | 100. | 7393 | 0 | 0 | 0.108 | 0.013 | 0 | 100. |
| E-F1c [3] | E | 99.9 | 7391 | 0 | 0 | 0 | 0 | 0 | 100. |
| S447-B1c [3] | S | 100. | 7393 | 0 | 0 | 0 | 100. | 100. | 0 |
| N15-B1c [6] | N | 99.9 | 7391 | 0 | 0 | 0 | 0 | 99.9 | 0 |
| E-B2 [7] | E | 99.9 | 7392 | 0 | 0 | 99.9 | 0 | 0 | 0 |
| N1-B1c [8] | N | 99.8 | 7381 | 0 | 0 | 0 | 0 | 0 | 0.203 |
| N1-F1c [8] | N | 98.9 | 7343 | 0 | 0 | 0 | 99.6 | 99.6 | 98.3 |
| N2-F1c [8] | N | 100. | 7392 | 0.013 | 0 | 0 | 100. | 100. | 100. |
| ORFlab-1-F1c [8] | Orf1ab | 99.9 | 7393 | 0 | 0 | 0 | 0.027 | 99.9 | 99.9 |
| NEB_orf1a-A-F1c [10] | Orf1a | 99.8 | 7390 | 0.068 | 0 | 0 | 0 | 0.013 | 99.8 |
| F1c [11] | N | 99.5 | 7390 | 0 | 0 | 0 | 0 | 0.013 | 0 |
| ORF1e-B1c [17] | Orf1ab | 100. | 7391 | 0 | 0 | 0 | 0 | 0 | 100. |
| LAMP_ORF1b-2_F2 [18] | Orf1b | 99.5 | 7388 | 99.6 | 0 | 0 | 0 | 0 | 0 |

Table S77: Percentage of mismatched base pairs at the first three positions of the 5′ end and the last three position of the 3′ end of FIP and BIP primers for 348 genomes of BA.3 subvariant. $N_{\text{al.}}$ is the number of alignments for each primer. Only those primers with $C_{\text{strict}}(5°C) = 0$ and $C_{\text{part.}}(5°C) > 90\%$.

| Primer | Gene Target | $C_{\text{part.}}$ (%) | $N_{\text{al.}}$ | 5′ end ⟶ (%) | | | ⟶ 3′ end (%) | | |
|---|---|---|---|---|---|---|---|---|---|
| As1_F1c [1] | Orf1a | 98.6 | 345 | 0 | 0 | 0 | 0 | 0 | 99.4 |
| As1e_F1c [1] | Orf1a | 99.1 | 348 | 0 | 0 | 0 | 0 | 99.1 | 99.1 |
| iLACO-F1c [1] | Orf1ab | 100. | 348 | 0 | 0 | 0 | 0 | 0 | 100. |
| E-F1c [3] | E | 97.7 | 340 | 0 | 0 | 0 | 0 | 0 | 100. |
| S447-B1c [3] | S | 99.7 | 347 | 0 | 0 | 0 | 100. | 100. | 0 |
| N15-B1c [6] | N | 99.7 | 347 | 0 | 0 | 0 | 0 | 100. | 0 |
| E-B2 [7] | E | 98.0 | 341 | 0 | 0 | 100. | 0 | 0 | 0 |
| N1-B1c [8] | N | 94.3 | 328 | 0 | 0 | 0 | 0 | 0 | 0 |
| N1-F1c [8] | N | 95.1 | 331 | 0 | 0 | 0 | 100. | 100. | 98.8 |
| N2-F1c [8] | N | 99.7 | 347 | 0 | 0 | 0 | 100. | 100. | 100. |
| ORFlab-1-F1c [8] | Orf1ab | 99.4 | 346 | 0 | 0 | 0 | 0 | 100. | 100. |
| NEB_orf1a-A-F1c [10] | Orf1a | 99.1 | 346 | 0 | 0 | 0 | 0 | 0 | 99.7 |
| ORF1e-B1c [17] | Orf1ab | 98.6 | 343 | 0 | 0 | 0 | 0 | 0 | 100. |
| LAMP_ORF1b-2_F2 [18] | Orf1b | 98.6 | 345 | 99.4 | 0 | 0 | 0 | 0 | 0 |

Table S78: Percentage of mismatched base pairs at the first three positions of the 5′ end and the last three position of the 3′ end of FIP and BIP primers for 629 genomes of BA.4 subvariant. $N_{\text{al.}}$ is the number of alignments for each primer. Only those primers with $C_{\text{strict}}(5°\text{C}) = 0$ and $C_{\text{part.}}(5°\text{C}) > 90\%$.

| Primer | Gene Target | $C_{\text{part.}}$ (%) | $N_{\text{al.}}$ | 5′ end ⟶ (%) | | | | | ⟶ 3′ end (%) |
|---|---|---|---|---|---|---|---|---|---|
| As1_F1c [1] | Orf1a | 99.5 | 628 | 0 | 0 | 0 | 0 | 0 | 99.7 |
| As1e_F1c [1] | Orf1a | 99.8 | 629 | 0 | 0 | 0 | 0 | 99.8 | 99.8 |
| iLACO-F1c [1] | Orf1ab | 100. | 629 | 0 | 0 | 0 | 0 | 0 | 100. |
| E-F1c [3] | E | 100. | 629 | 0 | 0 | 0 | 0 | 0 | 100. |
| S447-B1c [3] | S | 100. | 629 | 0 | 0 | 0 | 100. | 100. | 0 |
| N15-B1c [6] | N | 98.9 | 624 | 0 | 0 | 0 | 0 | 99.8 | 0 |
| E-B2 [7] | E | 99.2 | 626 | 0 | 0 | 99.7 | 0 | 0 | 0 |
| N2-F1c [8] | N | 99.4 | 625 | 0 | 0 | 0 | 100. | 100. | 100. |
| ORFlab-1-F1c [8] | Orf1ab | 100. | 629 | 0 | 0 | 0 | 0 | 100. | 100. |
| NEB_orf1a-A-F1c [10] | Orf1a | 99.4 | 625 | 0.16 | 0 | 0 | 0 | 0 | 100. |
| ORF1e-B1c [17] | Orf1ab | 99.8 | 629 | 0 | 0 | 0 | 0 | 0 | 99.8 |
| LAMP_ORF1b-2_F2 [18] | Orf1b | 99.8 | 628 | 100. | 0 | 0 | 0 | 0 | 0 |

Table S79: Percentage of mismatched base pairs at the first three positions of the 5′ end and the last three position of the 3′ end of FIP and BIP primers for 1231 genomes of BA.5 subvariant. $N_{\text{al.}}$ is the number of alignments for each primer. Only those primers with $C_{\text{strict}}(5°C) = 0$ and $C_{\text{part.}}(5°C) > 90\%$.

| Primer | Gene Target | $C_{\text{part.}}$ (%) | $N_{\text{al.}}$ | 5′ end ⟶ (%) | | | ⟶ 3′ end (%) | | |
|---|---|---|---|---|---|---|---|---|---|
| As1_F1c [1] | Orf1a | 99.7 | 1229 | 0 | 0 | 0 | 0 | 0 | 99.8 |
| As1e_F1c [1] | Orf1a | 99.8 | 1231 | 0 | 0 | 0 | 0 | 99.7 | 99.7 |
| iLACO-F1c [1] | Orf1ab | 99.9 | 1230 | 0 | 0 | 0 | 0 | 0 | 100. |
| E-F1c [3] | E | 99.5 | 1227 | 0 | 0 | 0 | 0 | 0 | 99.8 |
| S447-B1c [3] | S | 99.8 | 1230 | 0 | 0 | 0 | 99.9 | 99.9 | 0 |
| N15-B1c [6] | N | 98.9 | 1221 | 0 | 0 | 0 | 0 | 99.7 | 0 |
| E-B2 [7] | E | 99.8 | 1229 | 0 | 0 | 99.9 | 0 | 0 | 0 |
| N1-B1c [8] | N | 99.0 | 1219 | 0 | 0 | 0 | 0 | 0 | 0.33 |
| N2-F1c [8] | N | 99.4 | 1223 | 0 | 0 | 0 | 100. | 100. | 100. |
| ORFlab-1-F1c [8] | Orf1ab | 99.9 | 1231 | 0 | 0 | 0 | 0 | 99.9 | 99.9 |
| NEB_orf1a-A-F1c [10] | Orf1a | 97.1 | 1197 | 0 | 0 | 0 | 0.083 | 0 | 99.8 |
| ORF1e-B1c [17] | Orf1ab | 96.0 | 1223 | 0 | 0 | 0 | 0 | 0 | 96.6 |
| LAMP_ORF1b-2_F2 [18] | Orf1b | 98.8 | 1226 | 99.2 | 0 | 0 | 0 | 0 | 0 |
